# N_2_O-respiring bacteria in biogas digestates for reduced agricultural emissions

**DOI:** 10.1101/2020.11.16.384990

**Authors:** Kjell Rune Jonassen, Live H. Hagen, Silas H.W. Vick, Magnus Ø. Arntzen, Vincent G.H. Eijsink, Åsa Frostegård, Pawel Lycus, Lars Molstad, Phillip B. Pope, Lars R. Bakken

## Abstract

Inoculating agricultural soils with N_2_O-respiring bacteria (NRB) can reduce N_2_O-emissions, but would be impractical as a standalone operation. Here we demonstrate that digestates obtained after biogas production are suitable substrates and vectors for NRB. We show that indigenous NRB in digestates grew to high abundance during anaerobic enrichment under N_2_O. Gas-kinetics and meta-omic analyses showed that these NRB's, recovered as metagenome-assembled genomes (MAGs), grew by harvesting fermentation intermediates of the methanogenic consortium. Three NRB's were isolated, one of which matched the recovered MAG of a *Dechloromonas*, deemed by proteomics to be the dominant producer of N_2_O-reductase in the enrichment. While the isolates harbored genes required for a full denitrification pathway and could thus both produce and sequester N_2_O, their regulatory traits predicted that they act as N_2_O sinks in soil, which was confirmed experimentally. The isolates were grown by aerobic respiration in digestates, and fertilization with these NRB-enriched digestates reduced N_2_O emissions from soil. Our use of digestates for low-cost and large-scale inoculation with NRB in soil can be taken as a blueprint for future applications of this powerful instrument to engineer the soil microbiome, be it for enhancing plant growth, bioremediation, or any other desirable function.

## Introduction

Nitrous oxide is an intermediate in the nitrogen cycle and a powerful greenhouse gas emitted in large volumes from agricultural soils, accounting for ~1/3 of total anthropogenic N_2_O emissions (Tian et al 2020). Reduced emissions can be achieved by minimizing the consumption of fertilizer nitrogen through improved agronomic practice and reduction of meat consumption (Snyder et al 2014, Sutton et al, 2011), but such measures are unlikely to do more than stabilize the global consumption of fertilizer N (Erisman et al 2008). This calls for more inventive approaches to reduce N_2_O emissions, targeting the microbiomes of soil (D’Hondt et al 2021), in particular the physiology and regulatory biology of the organisms involved in production and consumption of N_2_O in soil (Bakken and Frostegård 2020).

N_2_O turnover in soil involves several metabolic pathways, controlled by a plethora of fluctuating physical and chemical variables (Butterbach-Bahl et al 2013, Hu et al 2015). Heterotrophic denitrification is the dominant N_2_O source in most soils, while autotrophic ammonia oxidation may dominate in well drained calcareous soils (Song et al 2018 and references therein). Heterotrophic denitrifying organisms are both sources and sinks for N_2_O because N_2_O is a free intermediate in their stepwise reduction of nitrate to dinitrogen (NO_3_^−^→NO_2_^−^→NO→N_2_O→N_2_). Denitrification involves four enzymes collectively referred to as denitrification reductases: nitrate reductase (Nar/Nap), nitrite reductase (Nir), nitric oxide reductase (Nor) and nitrous oxide reductase (Nos), encoded by the genes *nar*/*nap*, *nir*, *nor* and *nosZ*, respectively. Oxygen is a strong repressor of denitrification, both at the transcriptional and the metabolic level (Zumft 1997, Qu et al 2016). Many organisms have truncated denitrification pathways, lacking from one to three of the four reductase genes (Shapleigh 2013, Lycus et al 2017), and truncated denitrifiers can thus act as either N_2_O producers (organisms without *nosZ*) or N_2_O reducers (organisms with *nosZ* only). The organisms with *nosZ* only, coined non-denitrifying N_2_O-reducers (Sanford et al 2013), have attracted much interest as N_2_O sinks in the environment (Hallin et al 2018). Of note, organisms with a full-fledged denitrification pathway may also be strong N_2_O sinks depending on the relative activities and regulation of the various enzymes in the denitrification pathway (Lycus et al 2018; Mania et al 2020). Despite their promise, feasible ways to utilize N_2_O-reducing organisms to reduce N_2_O emissions have not yet emerged.

A soil with a strong N_2_O-reducing capacity will emit less N_2_O than one dominated by net N_2_O producing organisms, as experimentally verified by Domeignoz-Horta et al (2016), who showed that soils emitted less N_2_O if inoculated with large numbers (10^7^ - 10^8^ cells g^−1^ soil) of organisms expressing Nos as their sole denitrification reductase. As a standalone operation, the large-scale production and distribution of N_2_O-respiring bacteria would be prohibitively expensive and impractical. However, the use of N_2_O-respiring bacteria could become feasible if adapted to an existing fertilization pipeline, such as fertilization with the nitrogen- and phosphate-rich organic waste (digestate) generated by biogas production in anaerobic digesters. Anaerobic digestion (**AD**) is already a core technology for treating urban organic wastes, and is expected to treat an increasing proportion of the much larger volumes of waste produced by the agricultural sector (**Figure 1**), as an element of the roadmap towards a low-carbon circular economy (Scarlat et al 2018). This means that digestates from **AD** are likely to become a major organic fertilizer for agricultural soils, with a huge potential for reducing N_2_O emissions if enriched with N_2_O-respiring bacteria prior to application.

**Figure 1.**
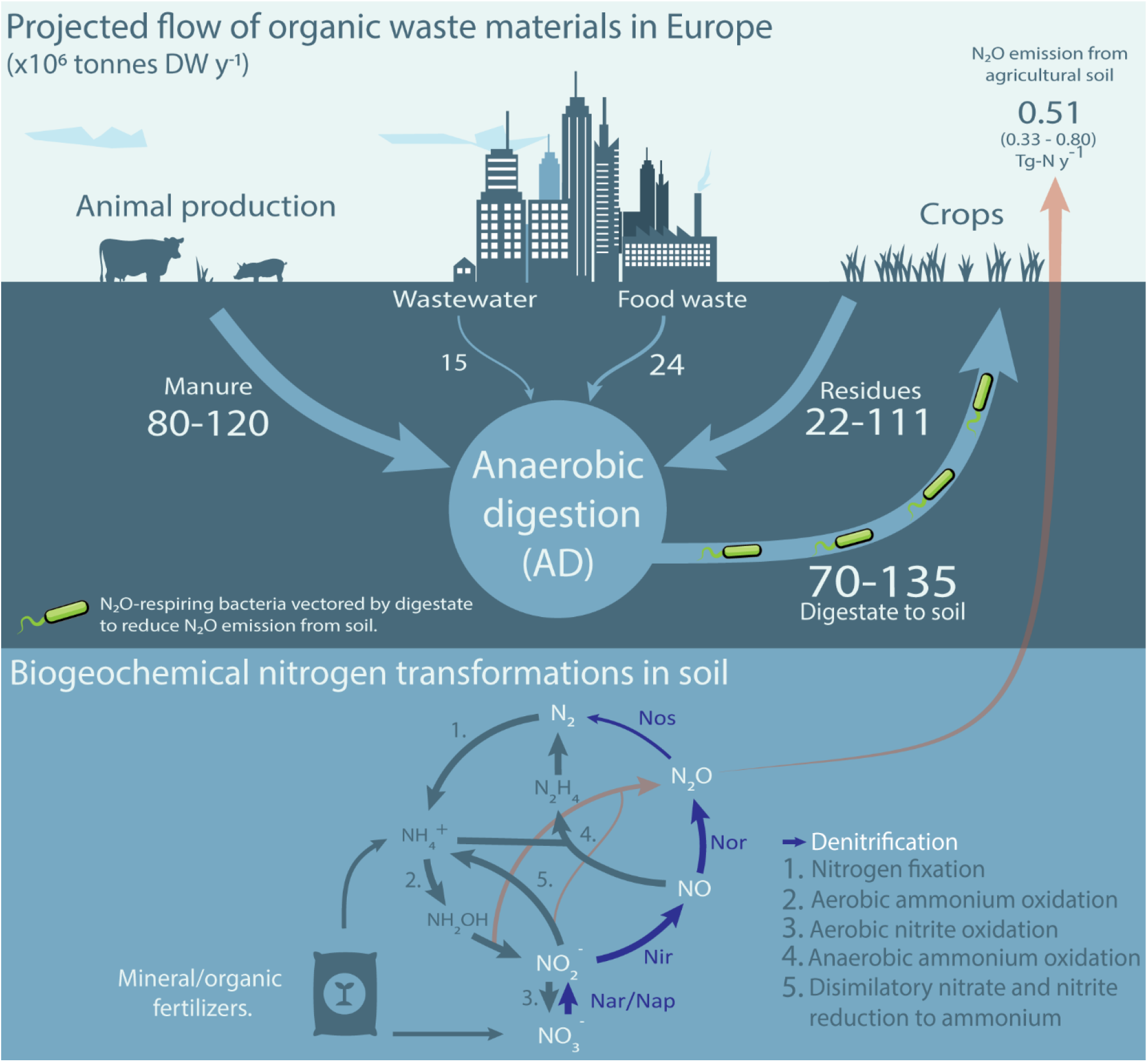
Possible biomass streams in a future circular economy with a central role for anaerobic digestion. Solid arrows (top section) show streams of biomass available for anaerobic digestion (AD). Numbers indicate known estimates of currently used or potentially available amounts in Europe, in million tonnes dry-weight (DW) per year (Foged et al 2011, Holm-Nielsen et al 2009, Stenmarck et al 2016, Meyer et al 2018). The arrow from anaerobic digestion to agricultural soil, indicates a credible pathway for digestate enriched with N_2_O-reducing bacteria (assuming enrichment at **AD** site); fertilization with such enriched digestates strengthens the N_2_O sink capacity of the soil, hence reducing N_2_O emissions. N_2_O emissions from agricultural soil in Europe are estimated at 0.51 tG per year (min 0.33 – max 0.80), representing some 48 % of total European N_2_O emissions (Tian et al 2020), which account for approximately 3.5 % of the global warming effect from European greenhouse gas emissions and 35 % of the global warming effect from European agriculture (Eurostat 2018). The lower half of the picture shows the microbial nitrogen transformations underlying these N_2_O emissions, which are fed by fertilizers. Today, **AD** is primarily used for treating urban organic wastes, which comprise only ~10 % of the biomass potentially available for **AD**. The amount of biomass treated by **AD** is expected to increase by an order of magnitude when adopted on a large scale in the agricultural sector. This would generate 70-135 Mt DW of digestate annually (assuming 50% degradation by AD), which is equivalent to 400-780 kg DW ha^−1^ y^−1^ if spread evenly on the total farmland of Europe (173 million ha).

Here we provide the first proof of this promising concept. Firstly, we demonstrate selective enrichment and isolation of fast-growing digestate-adapted N_2_O-respiring bacteria using a digestate from a wastewater treatment plant. Secondly, we demonstrate that the use of digestates enriched with such organisms as a soil amendment reduces the proportion of N leaving soil as N_2_O, confirming the suitability of such digestates for this purpose. Analysis of the enrichment process with multi-omics and in-depth monitoring of gas kinetics provides valuable insights into Nos-synthesis by the various enriched taxa, and the metabolic pathways of the anaerobic consortium providing substrates for these enriched N_2_O-respiring organisms.

## Materials and methods

<u>The digestates</u> were taken from two anaerobic digesters, one mesophilic (37 °C) and one thermophilic (52 °C), which were running in parallel, producing biogas from sludge produced by a wastewater treatment. The sludge was a poly-aluminum chloride (PAX-XL61™, Kemira) and ferric chloride (PIX318™, Kemira) precipitated municipal wastewater sludge, with an organic matter content of 5.6% (w/w). Both digestors reduced the organic matter by approximately 60%, producing digestates containing ~2.1 % organic matter, 1.8-1.9 g NH_4_ ^+^-N L^−1^, ~16 and 32 Meq VFA L^−1^, pH=7.6-7.8 and 8.2; mesophilic and thermophilic, respectively (see Suppl Methods 1 for further details). The digestates were transported to the laboratory in 1 L insulated steel-vessels and used for incubation experiments 3-6 hours after sampling.

<u>The robotized incubation system</u> developed by Molstad et al (2007, 2016) was used in all experiments where gas kinetics was monitored. The system hosts 30 parallel stirred batches in 120 mL serum vials, crimp sealed with gas tight butyl rubber septa, which are monitored for headspace concentration of O_2_, N_2_, N_2_O, NO, CO_2_ and CH_4_ by frequent sampling. After each sampling, the system returns an equal volume of He, and elaborated routines are used to account for the gas loss by sampling to calculate the production/consumption-rate of each gas for each time interval between two samplings. More details are given in Suppl Methods 2.

<u>Enrichment culturing</u> of N_2_O-respiring bacteria (NRB) in digestate was done as stirred (300 rmp) batches of 50 mL digestate per vial. Prior to incubation, the headspace air was replaced with Helium by repeated evacuation and He-filling (Molstad et al 2007), and supplemented with N_2_O, and N_2_O in the headspace was sustained by repeated injections in response to depletion. Liquid samples (1 mL) were taken by syringe, for metagenomic and metaproteomic analyses, and for quantification of volatile fatty acids (VFA) and 16srDNA abundance. The samples were stored −80 °C before analyzed. The growth of NRB in the enrichments was modelled based on the N_2_O reduction kinetics. The modelling and the analytic methods (quantification of VFA and 16srDNA abundance) are described in detail in Suppl Methods 3.

<u>Metagenomics and metaprotomics</u>: Sequencing of DNA (Illumina HiSec4000), and the methods for Metagenome-Assembled Gemome (MAG) binning, and the phylogenetic placement of the MAGs is described in detail in Suppl Methods 4. Proteins were extracted and digested to peptides, which were analyzed by nanoLC-MS/MS, and the acquired spectra were inspected, using the metagenome-assembled genomes (149 MAGs) as a scaffold (Suppl Methods 5).

<u>Isolation of N_2_O-respiring bacteria (NRB) (</u>Suppl Methods 6). NRB present in the enrichment cultures were isolated by spreading diluted samples on agar plates with different media composition, then incubated in an anaerobic atmosphere with N_2_O. Visible colonies were re-streaked and subsequently cultured under aerobic conditions, and 16s-sequenced. Three isolates, **AS** (*Azospira* sp), **AN** (Azonexus sp) and **SP** (*Pseudomonas* sp) (names based on their 16s sequence), were selected for genome sequencing, characterization of their denitrification phenotypes, and for testing their effect as N_2_O sinks in soil.

<u>Genome sequencing and phenotyping of isolates.</u> Three isolates were genome sequenced and compared with MAG's of the enrichment culture (Suppl Methods 7). The isolates' ability to utilize various organic C substrates was tested on BiOLOG Phenotype MicroArray**^TM^** microtiter plates, and their characteristic regulation of denitrification was tested through a range of incubation experiments as in previous investigations (Bergaust et al 2010, Liu et al 2014, Lycus et al 2018, Mania et al 2020), by monitoring the kinetics of O_2_, N_2_, N_2_O, NO and NO_2_^−^ throughout the cultures' depletion of O_2_ and transition from aerobic to anaerobic respiration in stirred batch cultures with He + O_2_ (+/- N_2_O) in the headspace (Suppl Methods 8). The kinetics of electron flow throughout the oxic and anoxic phase in these experiments were used to assess if the organisms were *bet hedging*, as demonstrated for *Paracoccus denitrificans* (Lycus et al 2018), i.e. that only a minority of cells express nitrate- and/or nitrite-reductase, while all express Nos, in response in response to oxygen depletion. Putative *bet hedging* was corroborated by measuring the abundance of nitrate-, nitrite- and nitrous oxide reductase (Suppl Methods 9).

<u>N_2_O mitigation experiments</u> (Suppl Methods 9). To assess the capacity of the isolates to reduce the N_2_O emission from soil, they were grown aerobically in sterilized digestate, which was then added to soil in microcosms, for measuring the NO-, N_2_O- and N_2_- kinetics of denitrification in the soil. For comparison, the experiments included soils amended with sterilized digestate, live digestate (no pretreatment), and digestate in which N_2_O-reducing bacteria had been enriched by anaerobic incubation with N_2_O (as for the initial enrichment culturing).

### Data availability

The sequencing data for this study have been deposited in the European Nucleotide Archive (ENA) at EMBL-EBI under accession number PRJEB41283 (isolates AN, AS and PS) and PRJEB41816 (metagenome) (https://www.ebi.ac.uk/ena/browser/view/PRJEBxxxx). Functionally annotated MAGs and metagenomic assembly are available in FigShare (DOI: 10.6084/m9.figshare.13102451 and 10.6084/ m9.figshare.13102493). The proteomics data has been deposited to the ProteomeXchange Consortium (http://proteomecentral.proteomexchange.org) via the PRIDE partner repository (Vizcaino et al 2013) with the dataset identifier PXD022030* and PXD023233** for the metaproteome and proteome of *Azonexus* sp. AN, respectively.

## Results and Discussion

### Enrichment of indigenous N_2_O-respiring bacteria (NRB) in digestates

We hypothesized that suitable organisms could be found in anaerobic digesters fed with sewage sludge, since such sludge contains a diverse community of denitrifying bacteria stemming from prior nitrification/denitrification steps (Lu et al 2014). We further hypothesized that these bacteria could be selectively enriched in digestates by anaerobic incubation with N_2_O. We decided to enrich at 20 °C, rather than at the temperatures of the anaerobic digesters (37 and 52 °C), to avoid selecting for organisms unable to grow within the normal temperature range of soils.

The digestates were incubated anaerobically as stirred batch cultures with N_2_O in the headspace (He atmosphere), and the activity and apparent growth of N_2_O reducers was assessed by monitoring the N_2_O-reduction to N_2_. **Figure 2A** shows the results for the first experiment, where culture vials were liquid samplws were taken at three time points (0, 115 and 325 h) for metagenomics, metaproteomics, and quantification of 16S rDNA and volatile fatty acids (VFAs). N_2_O was periodically depleted (100-140 h) in this experiment, precluding detailed analysis of the growth kinetics throughout. This was avoided in the second enrichment, for which complete gas data are shown in **Figure 2BC**. Apart from the deviations caused by the temporary depletion of N_2_O in the first experiment, both experiments showed very similar N_2_ production rates (**Figures 2B and S1B**). The gas kinetics of the second enrichment are discussed in detail below.

**Figure 2:**
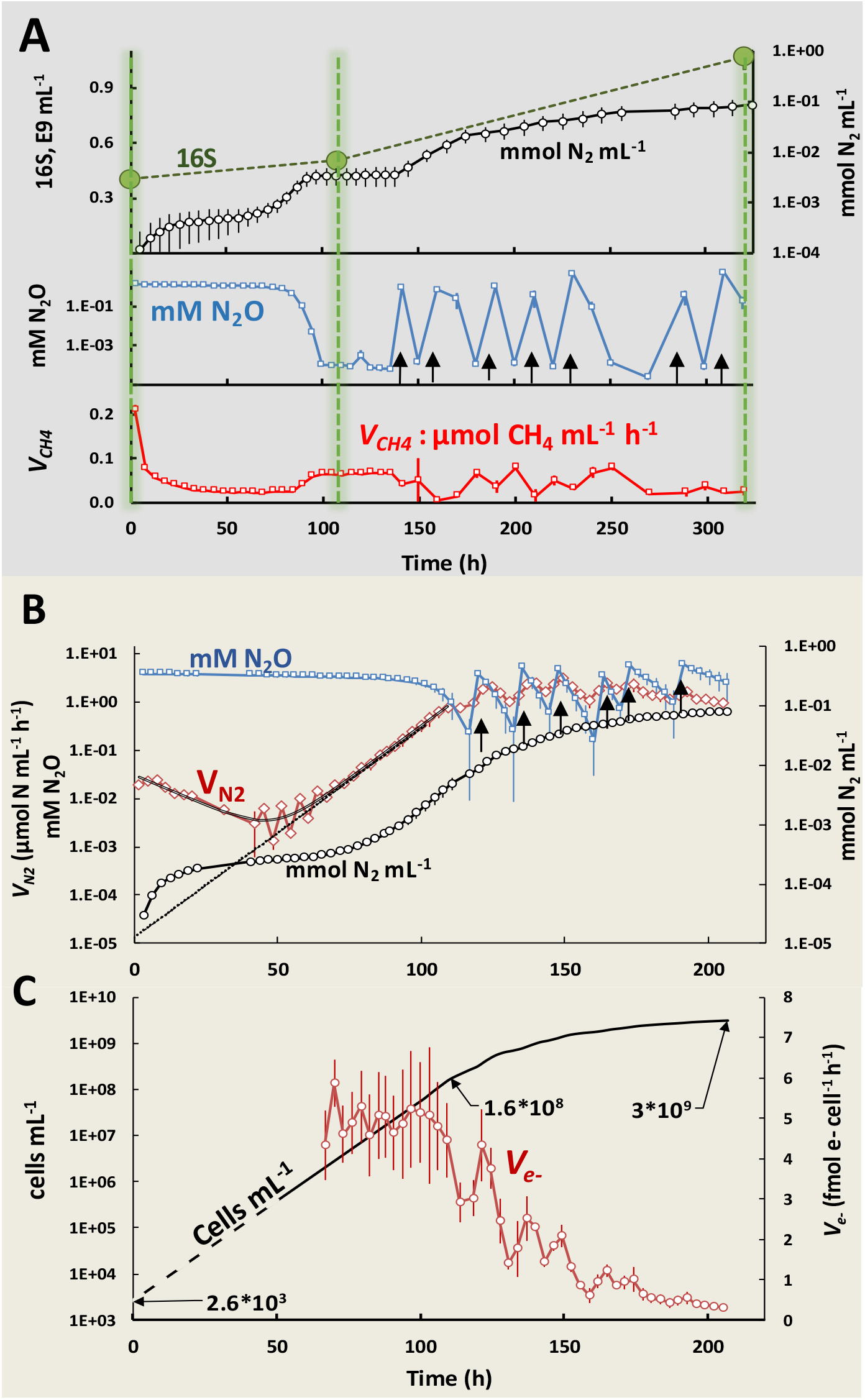
Gas kinetics in anaerobic enrichment cultures with digestate. <u>Panel A</u> shows results for the enrichment culture (triplicate culture vials) sampled for metagenomics, metaproteomics, quantification of volatile fatty acids (VFAs) and 16S rDNA abundance (sampling times = 0, 115 and 325 hours, marked by vertical green lines). The top panel shows the amounts of N_2_ produced (mmol N_2_ L^−1^ digestate, log scale) and 16S rDNA copy numbers. The mid panel shows the concentration of N_2_O in the digestate (log scale), which was replenished by repeated injections from t=140 h and onwards (indicated by black arrows). The bottom panel shows the rate of methane production. Standard deviations (n=3) are shown as vertical lines in all panels. <u>Panel B and C</u> show the results of a repeated enrichment experiment where N_2_O-depletion (as seen at t=100-140 h in panel A) was avoided, to allow more precise assessment and modelling of growth kinetics. <u>Panel B</u>: N_2_O concentration in the digestate (mM N_2_O), rate of N_2_-production (*V_N2_*) and N_2_ produced (mmol N_2_ mL^−1^ digestate), all log scaled. The curved black line shows the modelled *V_N2_* assuming two populations, one growing exponentially (μ = 0.1 h^−1^), and one whose activity was dying out gradually (rate = - 0.03 h^−1^). The dotted black line is the activity of the exponentially growing population extrapolated to time=0. <u>Panel C</u> shows the modelled density (cells mL^−1^) of cells growing by N_2_O respiration, extrapolated back to t=0 h (dashed line), and the cell specific respiratory activity (*V_e_^−^*, fmol electrons cell^−1^ h^−1^), which declined gradually after 110 h. Standard deviations (n = 3) are shown as vertical lines. **Figure S1** provides additional data for the experiment depicted in Panel A, as well as a detailed description of the modelling procedures and their results.

**Figure 2B** shows declining rates of N_2_-production (*V_N2_*) during the first 50 h, followed by exponential increase. This was modelled as the activity of two groups of NRB, one growing exponentially from low initial abundance, and one which was more abundant initially, but whose activity declined gradually (further explained in **Figure S1**). The modelling, indicated that the cell density of the growing NRB increased exponentially (specific growth rate, μ = 0.1 h^−1^) from a very low initial density (~2.5·10^3^ cells mL^−1^) to 1.6·10^8^ cells mL^−1^ after 110 h, and continued to increase at a gradually declining rate to reach ~3·10^9^ cells mL^−1^ at the end of the incubation period (215 h). The modelled cell-specific electron flow rate (*V_e_^−^*, **Figure 2C**) was sustained at around 5 fmol e^−^ cell^−1^ h^−1^ during the exponential growth, and declined gradually thereafter, as the number of cells continued to increase, while the overall rate of N_2_O-respiration remained more or less constant (*V_N2_*, **Figure 2B**). Enrichment culturing as shown in **Figure 2BC** was repeated three times, demonstrating that the characteristic N_2_ production kinetics was highly reproducible (**Figure S2**).

The provision of substrate for the N_2_O-respiring bacteria can be understood by considering the enrichment culture as a continuation of the metabolism of the anaerobic digester (**AD**), albeit slowed down by the lower temperature (20 °C, versus 37 °C in the digester). In **AD**, organic polymers are degraded and converted to CO_2_ and CH_4_ through several steps, conducted by separate guilds of the methanogenic microbial consortium: 1) hydrolysis of polysaccharides to monomers by organisms with carbohydrate-active enzymes, 2) primary fermentation of the resulting monomers to volatile fatty acids (VFAs), 3) secondary fermentation of VFAs to acetate, H_2_ and CO_2_, and 4) methane production from acetate, CO_2_, H_2_, and methylated compounds. By providing N_2_O to this (anaerobic) system, organisms that respire N_2_O can tap into the existing flow of carbon, competing with the methanogenic consortium for intermediates, such as monomeric sugars, VFAs (such as acetate) and hydrogen (Stams et al 2003). Thus, the respiration and growth of the N_2_O-respiring bacteria is sustained by a flow of carbon for which the primary source is the depolymerization of organic polymers. It is possible that the retardation of growth after ~100 h of enrichment was due to carbon becoming limiting. Thus, at this point, the population of N_2_O-respiring organisms may have reached high enough cell densities to reap most of the intermediates produced by the consortium.

Parallel incubations of digestates without N_2_O confirmed the presence of an active methanogenic consortium, sustaining a methane production rate of ~0.2 μmol CH_4_ mL^−1^ h^−1^ throughout (**Figure S3**). Methane production was inhibited by N_2_O, and partly restored in periods when N_2_O was depleted (**Figure 2A, Figures S3**&**S4**). We also conducted parallel incubations with O_2_ and NO_3_^−^ as electron acceptors. These incubations showed that methanogenesis was completely inhibited by NO_3_-, and partly inhibited by O_2_ (concentration in the liquid ranged from 20 to 90 μM O_2_) (**Figures S3**). The rates of O_2_ and NO_3_^−^ reduction indicated that the digestate contained a much higher number of cells able to respire O_2_ and NO_3_^−^ than cells able to respire N_2_O (**Figure S5A-C**). During the enrichment culturing with NO_3_^−^, almost all reduced nitrogen appeared in the form of N_2_O during the first 50 h (**Figure S5E**), another piece of evidence that in the digestate (prior to enrichment culturing), the organisms reducing NO_3_^−^ to N_2_O outnumbered those able to reduce N_2_O to N_2_. The measured production of CH_4_ and electron flows to electron acceptors deduced from measured gases (N_2_, O_2_ and CO_2_) were used to assess the effect of the three electron acceptors (N_2_O, NO_3_^−^ and O_2_) on C-mineralization. While oxygen appeared to have a marginal effect, NO_3_^−^ and N_2_O caused severe retardation of C-mineralization during the first 50 and 100 h, respectively (**Figure S5A-D**). This retarded mineralization is plausibly due to the inhibition of methanogenesis, causing a transient accumulation of H_2_ and VFAs until the N_2_O-reducing bacteria reach a cell density that allowed them to effectively reap these compounds. This was corroborated by measurements of H_2_ and VFAs (**Figure S13**).

To track the origin of the enriched N_2_O-respiring bacteria in the digestate, we considered the possibility that these are indigenous wastewater-sludge bacteria that survive the passage through the anaerobic digester, which had a retention time of 20-24 days. We assessed survival of N_2_O-respiring bacteria by comparing the N_2_O reduction potential of wastewater sludge and the digestate. The results indicated that ≤ 1/3 of the N_2_O-respiring bacteria in the sludge survived the passage (**Figure S6**). We also did enrichment culturing with a digestate from a thermophilic digester (52 °C) operated in parallel with the mesophilic digester (provided with the same feed), and found that it too contained N_2_O reducers that could be enriched, although the estimated initial numbers were orders of magnitude lower than in the mesophilic digestate **(Figure S7)**.

### MAG-centric metaproteomic analysis of the enrichment cultures

We analyzed the metagenome and metaproteome at three timepoints (0, 115 and 325 h, **Figure 2A**), to explore the effect of the anaerobic incubation with N_2_O on the entire microbial consortium, and to identify the organisms growing by N_2_O reduction. Metagenomic sequences were assembled and resultant contigs assigned to 278 metagenome-assembled genomes (MAGs), of which 149 were deemed to be of sufficient quality (completeness > 50% and contamination < 20%, Supplementary Data S1) for downstream analysis. The phylogenetic relationship and the relative abundance of the MAGs throughout the enrichment are summarized in **Figure 3**, which also shows selected features revealed by the combined metagenomic and metaproteomic analyses, including information about genes and detected proteins involved in N_2_O reduction, other denitrification steps, methanogenesis, syntrophic acetate oxidation and methane oxidation.

**Figure 3:**
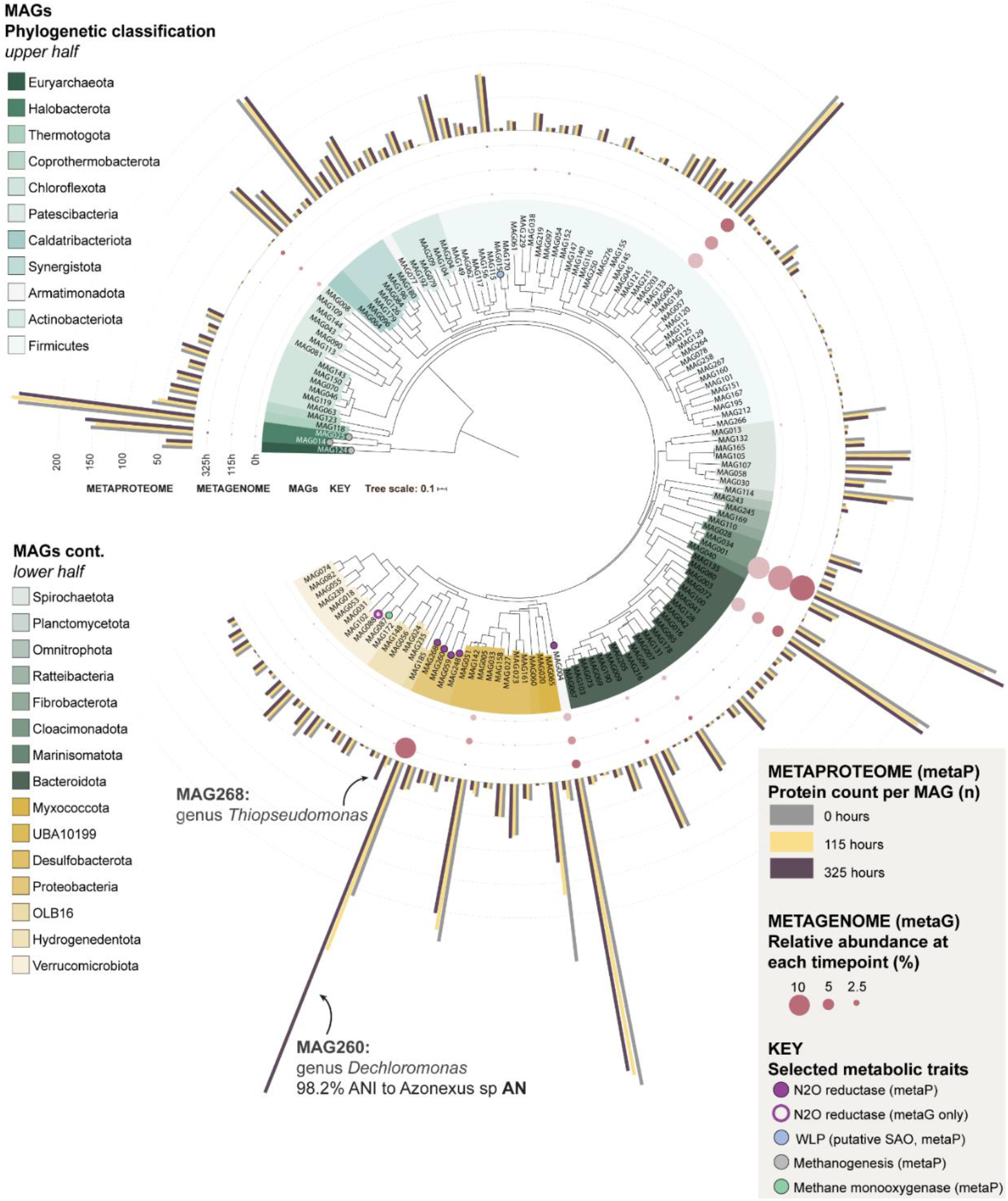
MAGs from the anaerobic enrichment culture with the mesophilic digestate. The figure shows a maximum likelihood tree indicating the phylogenetic placement of MAGs from the anaerobic enrichment. The tree was constructed from a concatenated set of protein sequences of single-copy genes. Taxonomic classification of the MAGs was inferred using the Genome Taxonomy Database (GTDB) and is displayed at the phylum level by label and branch coloring. Branch label decorations indicate the presence of genes involved in selected metabolic traits in the MAGs. The relative abundance of the MAG in the community as calculated from sequence coverage is indicated by bubbles at branch tips and bar charts indicate the number of detected proteins affiliated with each MAG at the three time points during incubation. Four of the 149 MAGs that met the completeness and contamination threshold for construction of the metaproteome database were lacking the universal single-copy marker genes and were omitted from the tree. Total protein counts per MAG were calculated by aggregating both secretome and cell-associated proteomes.

Closer inspections of the abundance of individual MAGs, based on their coverage in the metagenome and metaproteome, showed that the majority of the MAGs had a near constant population density throughout the incubation, while two MAGs (260 and 268) increased substantially (**Figure 4**; further analyses in **Supplementary Section B, Figures S8-S11**). The stable abundance of the majority indicates that the methanogenic consortium remained intact despite the downshift in temperature (20 °C versus 37 °C) and the inhibition of methanogenesis by N_2_O. Only 9 MAGs showed a consistent decline in abundance throughout the enrichment (**Table S1**). These MAGs could theoretically correspond to microbes whose metabolism is dependent on efficient H_2_ scavenging by methanogens (Schink 1997), but we found no genomic evidence for this, and surmise that organisms circumscribed by the declining MAGs were unable to adapt to the temperature downshift from 37 °C to 20 °C.

**Figure 4:**
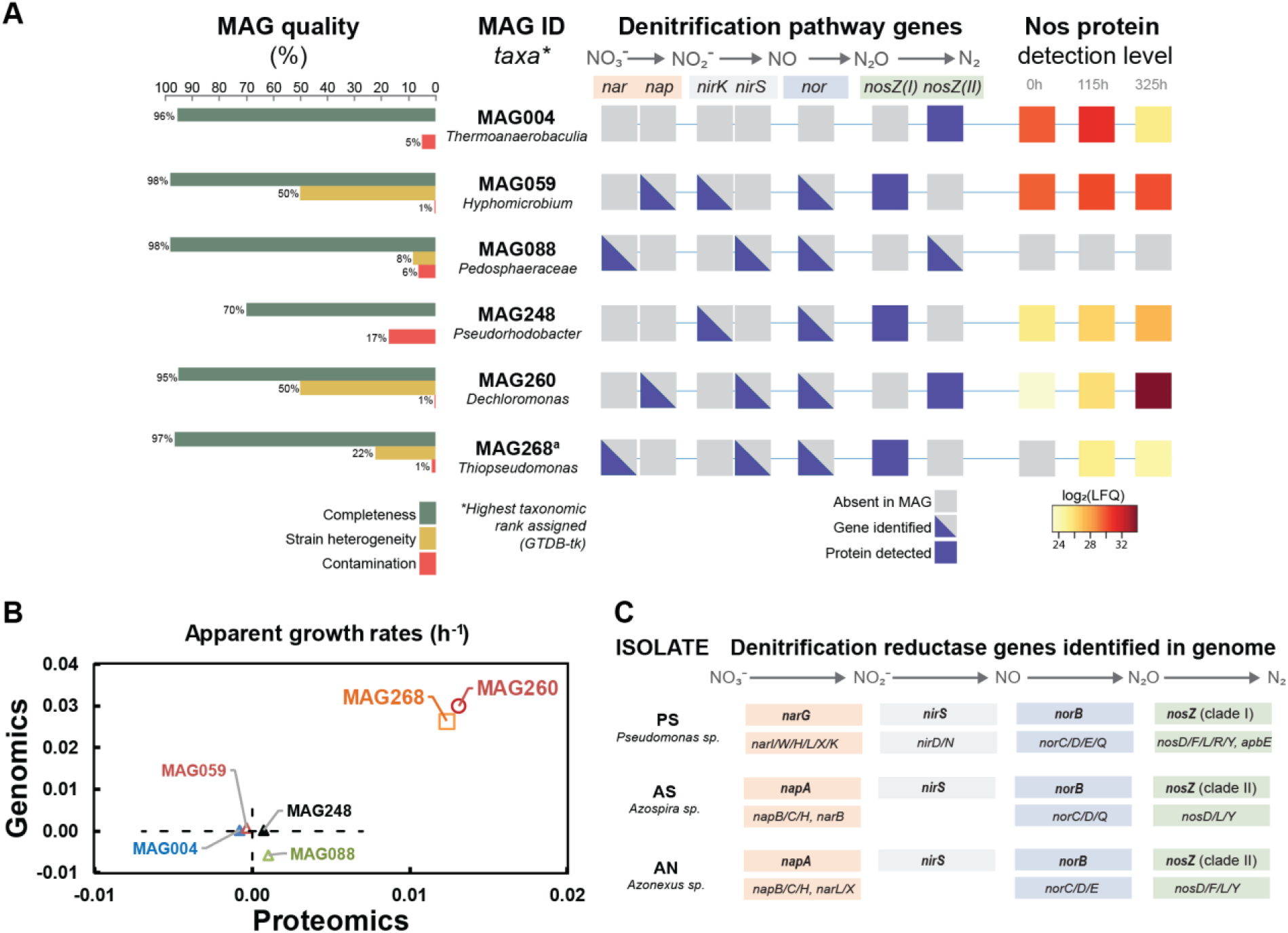
Overview of MAGs with *nosZ* and denitrification genes in isolated organisms. <u>Panel A</u> shows the quality (completeness, strain heterogeneity and contamination), taxonomic classification based on GTDB and NCBI, presence of denitrifying genes and proteins, and the detected levels of Nos (N_2_O reductase, encoded by *nosZ*) throughout the enrichment culturing for the six MAGs containing the *nosZ* gene (**Supplementary Data S1** and **S2**). Nos was detected in the proteome of five MAGs, but the detection level increased significantly throughout for only MAG260 and 248, respectively. None of the MAGs produced detectable amounts of the other denitrification reductases. ^a)^ LFQ values for one of the two detected predicted Nos proteins for MAG268 is shown. <u>Panel B</u> shows the apparent growth rates of the MAGs, based on their coverage in the metagenome and metaproteome (regression of ln(N) against time; see **Figure S11** for more details). <u>Panel C</u> shows the taxonomic classification (16S rDNA), working names (abbreviations) and denitrification genotypes of three isolates from the enrichment culturing. The genes coding catalytic subunits of denitrification reductases are shown in bold, above the accessory genes (Vaccaro et al 2016) that were also identified. More information about accessory genes is presented in **Figure S14**. The isolate **AN** has 98.2 % ANI to MAG260.

Six MAGs, including the two that were clearly growing (MAG260 & MAG268) contained the *nosZ* gene and thus had the genetic potential to produce N_2_O-reductase (Nos) (**Figure 4**). Nos proteins originating from five of these MAGs were detected in the metaproteome. Importantly, while all but one of these MAGs contained genes encoding the other denitrification reductases, none of these were detected in the metaproteome, suggesting that the organisms can regulate the expression of their denitrification machinery to suit available electron acceptors, in this case N_2_O. Three of the MAGs with detectable Nos in the proteome (MAG004, MAG059, MAG248) appeared to be non-growing during the enrichment. The detected levels of their Nos proteins remained more or less constant, and their estimated abundance in the metagenome and -proteome did not increase (**Figure 4B**). It is conceivable that these three MAGs belong to the initial population of N_2_O reducers whose N_2_O-reduction activity was present initially but gradually decreased during the early phase of the enrichment (**Figure 2A**). The two growing MAGs (MAG260 and MAG268) showed increasing Nos levels and increasing abundance both in terms of coverage and metaproteomic detection (**Figure 4B**), in proportion with the N_2_ produced (**Figure S11**). MAG260 reached the highest abundance of the two and accounted for 92% of the total detectable Nos pool at the final time point. MAG260 is taxonomically most closely affiliated with the genus *Dechloromonas* (GTDB, 97.9% amino acid similarity). Interestingly, Nap rather than Nar takes the role of nitrate reductase in MAG260 (**Figure 4**), which makes it a promising organism for N_2_O mitigation since organisms with Nap only (lacking Nar) preferentially channel electrons to N_2_O rather than to NO_3_^−^ (Mania et al 2020). MAG260, MAG004 and MAG088 contain a clade II *nosZ*, characterized by a *sec*-dependent signal peptide, in contrast to the more common *tat*-dependent clade I *nosZ*. The physiological implications of clade I versus clade II *nosZ* remains unclear. Organisms with *nosZ* Clade II have high growth yield and high affinity (low *k_s_*) for N_2_O, compared to thise with *nosZ* Clade II (Yoon et al 2016), suggesting a key role of *nosZ* Clade II organisms for N_2_O reduction in soil, but this was contested by Conthe et al (2018), who found that Clade I organisms had higher “catalytic efficiency” (*V_max_*/k_s_) than those with Clade II.

The apparent inhibition of methanogenesis by N_2_O seen in the present study has been observed frequently (Andalib et al 2011) and is probably due to inhibition of coenzyme M methyltransferase (Kengen et al 1988), which is a membrane bound enzyme essential for methanogenesis and common to all methanogenic archaea (Fischer et al 1992). The gas kinetics demonstrate that the inhibition of was reversible, being partly restored whenever N_2_O was depleted (**Figure 2**). In the enrichment culture where metagenomics and metaproteomics was monitored, several such incidents of N_2_O depletion occurred (**Figure 2A**) and during these periods CH_4_ accumulated to levels amounting to 10% of levels in control vials without N_2_O (**Figure S4B**). These observations suggest that methanogens would be able to grow, albeit sporadically, during the enrichment, which is corroborated by the sustained detection of the complete methanogenesis pathway, including the crucial coenzyme M methyl-transferase, of *Methanothrix* (MAG025), *Methanoregulaceae* (MAG014) and *Methanobacterium* (MAG124) at high levels in the metaproteome. In fact, both MAG coverage data and 16S rDNA copy numbers assessed by ddPCR suggested that the majority of the original methanogenic consortium continued to grow (**Supplementary Section B**). A tentative map of the metabolic flow of the methanogenic consortium, including the reaping of intermediates (monosaccharides, fatty acids, acetate and H_2_) by N_2_O-respiring bacteria is shown in **Figure S12.** Since methane production was inhibited from the very beginning of the incubation, while it took ~100 hours for the N_2_O-respiring bacteria to reach high enough numbers to become a significant sink for intermediates (**Figure 2**), one would expect transient accumulation of volatile fatty acids and H_2_, which was corroborated by measurements of these metabolites (**Figure S13**).

Of note, we detected methane monooxygenase and methanol dehydrogenase proteins from MAG087 and MAG059, respectively, in the metaproteome. This opens up the tantalizing hypothesis of N_2_O-driven methane oxidation, a process only recently suggested to occur (Valenzuela et al 2020; Cheng et al 2019). However, a close inspection of the N_2_O- and CH_4_-kinetics indicated that N_2_O-driven methane oxidation played a minor role (**Figure S4CD**).

### Isolation of N_2_O-respiring bacteria and their geno- and phenotyping

Whilst this enrichment culture could be used directly as a soil amendment, this approach is likely to have several disadvantages. First, it would require the use of large volumes of N_2_O for enrichment, a process which would be costly and require significant infrastructure. An alternative approach would be to introduce an axenic or mixed culture of digestate-derived, and likely digestate-adapted, N_2_O-respiring bacteria to sterilized/sanitized digestates. This approach has multiple benefits: 1) it would remove the need for N_2_O enrichment on site as isolates could be grown aerobically in the digestate material, 2) one could chose organisms with favorable denitrification genotypes and regulatory phenotypes, 3) the sanitation would eliminate the methanogenic consortium hence reducing the risk of methane emissions from anoxic micro-niches in the amended soil, and 4) sanitation of digestates aligns with current practices that require such a pretreatment prior to use for fertilization. For these reasons an isolation effort was undertaken to obtain suitable digestate-adapted N_2_O-respiring microorganisms from the N_2_O-enrichment cultures. These efforts resulted in the recovery of three axenic N_2_O-respiring bacterial cultures, which were subjected to subsequent genomic and phenotypic characterization.

The isolates were phylogenetically assigned to *Pseudomonas* sp. (**PS**), *Azospira* sp. **(AS)** and *Azonexus* sp. (**AN**) (working names in bold) based on full length 16S rDNA obtained from the sequenced genomes (accessions ERR4842639 - 40, **Table S2**, phylogenetic trees shown in **Figure S14**). All were equipped with genes for a complete denitrification pathway (**Figure 4C**). **AN** and **AS** carried *napAB*, encoding the periplasmic nitrate reductase (Nap) and *nosZ* clade II, whilst **PS** carried genes for the membrane bound nitrate reductase (Nar), encoded by *narG*, and *nosZ* clade I. All had *nirS* and *norBC*, coding for nitrite reductase (NirS) and and nitric oxide reductase (Nor), respeictivey. Pairwise comparison of average nucleotide identities (ANI) with MAGs from the enrichment metagenomes showed that the isolate **AN** matched the *Dechloromonas*-affiliated MAG260 with 98.2 % ANI, suggesting the isolate is circumscribed by MAG260 (Richter and Resselló-Móra 2009). Given the GTDB phylogeny of **AN** and MAG260 and the 16S rDNA gene homology of **AN** (95.2 % sequence identity to *Azonexus hydrophilus* DSM23864, **Fig S14C**), we conclude that **AN** likely represents a novel species within the *Azonexus* lineage. Unfortunately, the 16S rDNA gene was not recovered in MAG260, preventing direct comparison with related populations. No significant ANI matches in our MAG inventory were identified for the genomes of **PS** and **AS.**

The carbon catabolism profiles of the isolates were assayed using Biolog™ PM1 and PM2 microplates, to screen the range of carbon sources utilized (**Supplementary Section E)**. **PS** utilized a wide spectrum of carbon sources (amino acids, nucleic acids, volatile fatty acids (VFA), alcohols, sugar alcohols, monosaccharides and amino sugars), but only one polymer (laminarin). **AN** and **AS** could only utilize small VFAs (*eg*. acetate, butyrate), intermediates in the TCA cycle and/or the β-oxidation/methyl malonyl-CoA pathways of fatty acid degradation (*eg*. malate, fumarate, succinate), and a single amino acid (glutamate). Thus, all three would be able to grow in a live digestate by reaping the VFA's produced by the methanogenic consortium. While the utilization of VFAs as C-substrates is one of several options for **PS**, **AN** and **AS** appear to depend on the provision of VFAs. This was confirmed by attempts to grow the three isolates in an autoclaved digestate: while **PS** grew well and reached high cell densities without any provision of extra carbon sources, **AN** and **AS** showed early retardation of growth unless provided with an extra dose of suitable carbon source (glutamate, acetate, pyruvate or ethanol) (**Figure S25 and S26**). A high degree of specialization and metabolic streamlining may thus explain the observed dominance of **AN** (MAG260) during enrichment culturing.

To evaluate the potentials of these isolates to act as sinks for N_2_O, we characterized their denitrification phenotypes, by monitoring kinetics of oxygen depletion, subsequent denitrification and transient accumulation of denitrification intermediates (NO_2_^−^, NO, N_2_O). The experiments were designed to assess properties associated with strong N_2_O reduction such as 1) *bet hedging*, i.e. that all cells express N_2_O reductase while only a fraction of the cells express nitrite- and/or nitrate-reductase, as demonstrated for *Paracoccus denitrificans* (Lycus et al 2018); 2) strong metabolic preference for N_2_O-reduction over NO_3_^−^-reduction, as demonstrated for organisms with periplasmic nitrate reductase (Mania et al 2020). **Supplementary section F** provides the results of all the experiments and a synopsis of the findings. In short: *Azonexus* sp. (**AN**) had a clear preference for N_2_O over NO_3_^−^ reduction, but not over NO_2_^−^ reduction, ascribed to *bet hedging* with respect to the expression of nitrate reductase (a few cells express Nap, while all cells express Nos), which was corroborated by proteomics: the Nos/Nap abundance ratio was ~25 during the initial phase of denitrification (**Figure S17**). *Azospira* sp. (**AS**) had a similar preference for N_2_O over NO_3_ reduction, albeit less pronounced than in **AN**, and no preference for N_2_O over NO_2_^−^. *Pseudomonas* sp. (**PS**) showed a phenotype resembling that of *Paracoccus denitrificans* (Lycus et al 2018), with denitrification kinetics indicating that Nir is expressed in a minority of cells in response to O_2_ depletion, while all cells appeared to express N_2_O reductase. This regulation makes **PS** a more robust sink for N_2_O than the two other isolates, since it kept N_2_O extremely low even when provided with NO_2_^−^.

In summary, **PS** appeared to be the most robust candidate as a sink for N_2_O in soil for two reasons; 1) it can utilize a wide range of carbon substrates, and 2) its N_2_O sink strength is independent of the type of nitrogen oxyanion present (NO_2_^−^ or NO_3_^−^). In contrast, **AN** and **AS** appear to be streamlined for harvesting intermediates produced by anaerobic consortia, hence their metabolic activity in soil could be limited. In addition, they could be sources rather than sinks for N_2_O if provided with NO_2_^−^, which is likely to happen in soils, at least in soils of neutral pH, during hypoxic/anoxic spells (Lim et al 2018).

### Effects on N_2_O emissions

To assess if fertilization with digestates containing N_2_O-reducing bacteria could reduce N_2_O emissions from denitrification in soil, we conducted a series of incubation experiments with soils fertilized with digestates with and without N_2_O-reducing bacteria. The fertilized soils were incubated in closed culture vials containing He + 0.5 vol % O_2_, and O_2_, NO, N_2_O and N_2_ were monitored during oxygen depletion and anaerobic growth. The experiments included soils amended with digestates in which indigenous N_2_O-reducing bacteria had been enriched by anaerobic incubation with N_2_O (**Figure 2**), as well as autoclaved digestates in which the isolates from the current study had been grown by aerobic cultivation (see **Figures S25** & **S26** for cultivation details). The experiments included two types of control digestates: Live digestate (directly from the digester), and live digestate heated to 70 ^°^C for 2 hours (to eliminate most of the indigenous consortium). In all cases, 3 mL of digestate was added to 10 g of soil. Since soil acidity has a pervasive effect on the synthesis of functional N_2_O reductase (Liu et al 2014), we tested the digestates with two soils from a liming experiment (Nadeem et al 2020) with different pH (pH_CaCl2_ = 5.5 and 6.6).

The transient N_2_O accumulation during denitrification was generally higher in the acid than in the near-neutral soil (**Figure 5**), which was expected since the synthesis of functional Nos is hampered by low pH (Bergaust et al 2010, Liu et al 2014). Based on the kinetics of both N_2_ and N_2_O (see **Figure S27 and S28**), we calculated the N_2_O-index (***I_N2O_***) which is a measure of the molar amounts of N_2_O relative to N_2_+N_2_O in the headspace for a specific period (0-T), see equation at top of **Figure 5**). Low values of ***I_N2O_*** indicate efficient N_2_O-reduction. In this case, we calculated ***I_N2O_*** for the incubation period until 40% of the available NO_3_-had been recovered as N_2_+N_2_O (=***I_N2O_ 40***) and for the incubation period until 100% was recovered (***I_N2O_ 100***).

**Figure 5:**
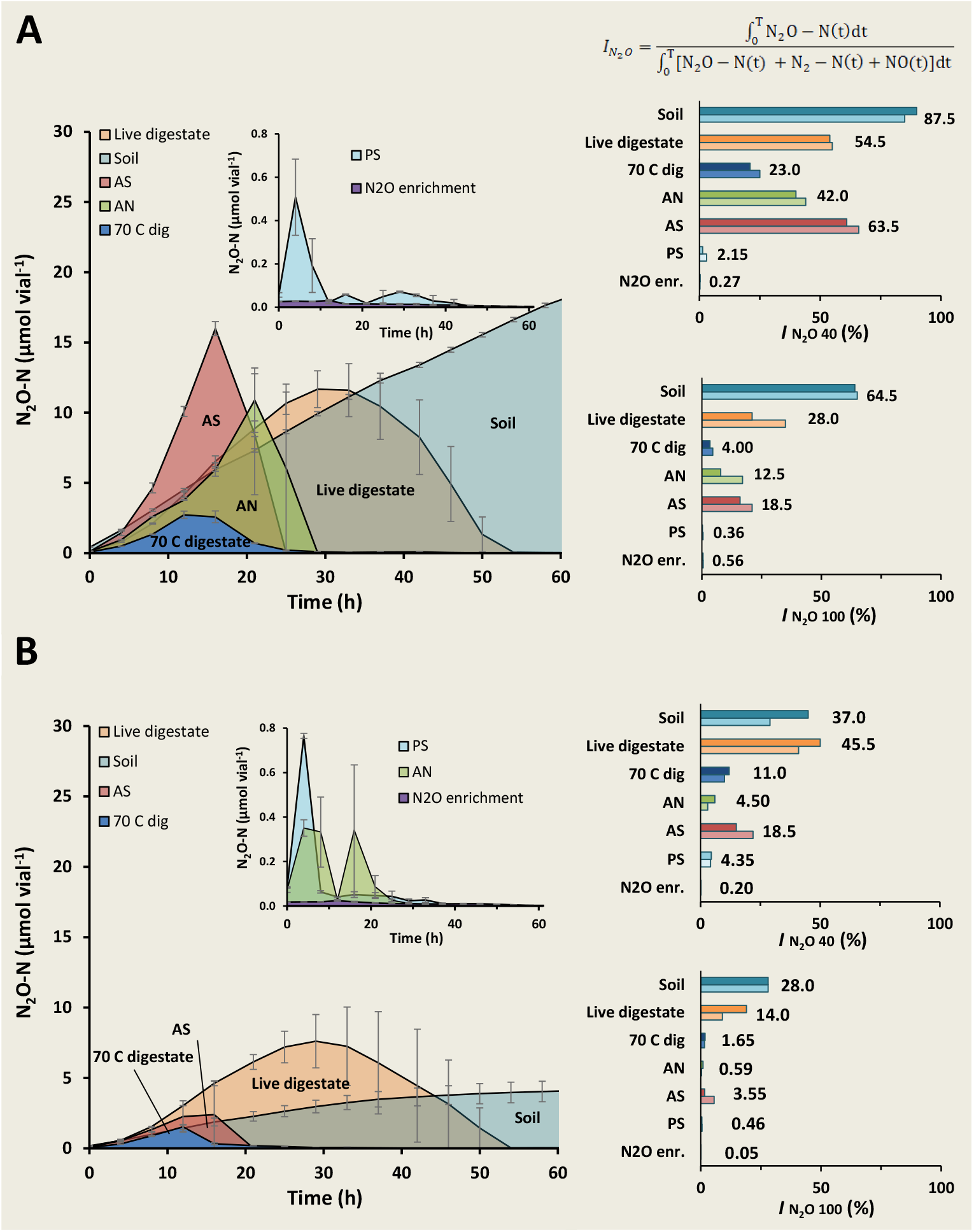
Soil incubations. N_2_O kinetics during incubation of soils amended with six different digestates and a control sample (soil only). Panel A shows results for the pH 5.5 soil, while panel B the pH 6.6 soil. The digestates treatments are: “Live digestate”, digestate directly from the anaerobic digester; “70 C dig”, live digestate heat treated to 70 °C for two hours; AN, AS and PS: autoclaved digestate on which isolates AN, AS and PS had been grown aerobically (see Figure **S25&S26** for details on the cultivation); “N_2_O enr”= digestate enriched with N_2_O-respiring bacteria (as in **Fig 2**). The left panels show the N_2_O levels observed during each treatment; the insets, with altered scaling, show N_2_O levels for treatments that resulted in very low N_2_O levels (the PS and N_2_O enr. treatments). The bar graphs to the right show the N_2_O indexes (***I_N2O_***, bar height = single culture vial values, numerical value = average of duplicate culture vials), which are calculated by dividing the area under the N_2_O-curve by the sum of the areas under the N_2_O and N_2_-curve, expressed as % (see equation in the figure and Liu et al 2014; the N_2_ curves are provided in **Figures S27&S28**). ***I_N2O_*** have proven to be a robust proxy for potential N_2_O emission from soil (Russenes et al 2016). Two ***I_N2O_*** values are shown: one for the timespan until 40% of the NO_3_^−^ -N was recovered as N_2_+N_2_O+NO (**I_N2O_ 40%**), and one for 100% recovery (**I_N2O_ 100%**). More details (including N_2_ and NO kinetics) are shown in **Figure S27 and S28.**

Extremely low ***I_N2O_*** values were recorded for the treatments with digestate in which N_2_O-reducing bacteria were enriched by anaerobic incubation with N_2_O, even in the acid soil. This is in line with the current understanding of how pH affects N_2_O-reduction: low pH slows down the synthesis of functional Nos, but once synthesized, it remains functional even at low pH (Bergaust et al 2010). Functional Nos had already been expressed during the enrichment and was evidently active after amendment to the soils.

***I_N2O_*** values were generally high for treatment with live digestate, which probably reflects that the digestate is dominated by N_2_O-producing organisms (**Figure S5E**). This interpretation is corroborated by the observed effect of heat-treating the live digestate; this lowered ***I_N2O_*** substantially.

The presence of the isolates in the digestates had clear but variable effects on ***I_N2O_***. Compared to the heat treated digestate (“70 C dig” Fig 5), **AN** and **AS** increased the ***I_N2O_***-values in the soil with pH=5.5, while in the soil with pH 6.6, their effect was marginal. The high ***I_N2O_*** for **AN** and **AS** in the acid soil plausibly reflect that the isolates were grown aerobically in the digestate, hence synthesizing their denitrification enzymes after transfer to soil, which would be hampered by low pH. In contrast to AN and AS, **PS** resulted in very low ***I_N2O_*** values in both soils, suggesting that this organism has an exceptional capacity to synthesize functional Nos at low pH.

These results show that the emission of N_2_O from soil fertilized with digestates can be manipulated by tailoring the digestate microbiome. Interestingly, measurements of methane in these soil incubations showed that the methanogenic consortia in digestates that had not been heat treated (i.e. the live digestate and the N_2_O enrichment) remained metabolically intact in the soil, and started producing methane as soon as N_2_O and nitrogen oxyanions had been depleted, while no methane was produced in the soils amended with autoclaved digestate, and that heated to 70 °C (**Figure S29**).

In an effort to determine the survival of the N_2_O-scavenging capacity of a digestate enriched with N_2_O reducers, we also tested its effect on soil N_2_O emissions after a 70-hour aerobic storage period (in soil or as enrichment culture, at 20 °C). These experiments demonstrated a sustained beneficial effect on ***I_N2O_*** after 70 hours of aerobic storage (**Figure S30**). This result indicates that the enrichment strategies discussed here are robust, although long-lasting storage experiments as well as field trials are needed.

## Concluding remarks

This feasibility study identifies an avenue for large scale cultivation of N_2_O reducers for soil application, which could be low cost if implemented as an add-on to biogas production systems. Further efforts should be directed towards selecting organisms that are both strong sinks for N_2_O and able to survive and compete in soil, to secure long-lasting effects on N_2_O emissions. A tantalizing added value would be provided by selecting organisms (or consortia of organisms) that are not only strong N_2_O-sinks, but also promote plant growth and disease resistance (Gao et al 2016, 2017).

Gas kinetics, metagenomics and metaproteomics revealed that the methanogenic consortium of the digestate remains active during anaerobic incubation with N_2_O, and that bacteria with an anaerobic respiratory metabolism grew by harvesting fermentation intermediates. The inhibition of methanogenesis by N_2_O implies that the respiring organisms would have immediate access to the electron donors that would otherwise be used by the methanogens, i.e. acetate and H_2_, while they would have to compete with fermentative organisms for the “earlier” intermediates such as alcohols and VFA. The importance of fermentation intermediates as a carbon source for the N_2_O-respiring bacteria would predict a selective advantage for organisms with a streamlined (narrow) catabolic capacity, i.e. limited to short fatty acids, and our results lend some support to this: the catabolic capacity of the organism that became dominant (MAG260, isolate **AN**) was indeed limited, as was also the case for isolate **AS**. Such organisms are probably not ideal N_2_O-sinks in soil because their ability to survive in this environment would be limited. Organisms with a wider catabolic capacity, such as the isolated *Pseudomonas* sp. (**PS**), are stronger candidates for long term survival and N_2_O-reducing activity in soil. The ideal organisms are probably yet to be found, however, and refinements of the enrichment culturing process are clearly needed.

The digestate used in this study contained N_2_O-respiring bacteria, most likely survivors from the raw sludge, which however, were clearly outnumbered by bacteria that are net producers of N_2_O. We surmise that the relative amounts of N_2_O-producers and N_2_O-reducers in digestates may vary, depending on the feeding material and configuration for the anaerobic digestion. This could explain the observed large variation of digestates on N_2_O emission from soils (Baral et al 2017, Herrero et al 2016). The high abundance of both NO_3_^−^ - and O_2_-respiring organisms in digestates has practical implications for the attempts to grow isolated strains in digestates: they could be outnumbered by the indigenous NO_3_^−^ - and O_2_-respiring organisms (**Figure S5**). Hence, we foresee that future implementation of this strategy will require a brief heat treatment or other sanitizing procedure. A bonus of such sanitation is that it eliminates methane production by the digestate in soil.

We failed to enrich organisms lacking all other denitrification genes than *nosZ*; the only reconstructed genome with *nosZ* only (**MAG004**) did not grow at all. Failure to selectively enrich such organisms by anaerobic incubation with N_2_O was also experienced by Conthe et al (2018). The organisms that did grow by respiring N_2_O in our enrichment, were all equipped with genes for the full denitrification pathway, although the only denitrification enzyme expressed/detected during the enrichment was Nos. This agrees with the current understanding of the gene regulatory network of denitrification; *nosZ* is the only gene whose transcription does not depend on the presence of NO_3_^−^, NO_2_^−^ or NO (Spiro 2016), which were all absent during the enrichment.

Two of the reconstructed MAGs had periplasmic nitrate reductase (*nap*), as was the case for two of the three isolates (AN and AS). This in itself would predict preference for N_2_O-over NO_3_^−^ reduction at a metabolic level (Mania et al 2020), but otherwise their potential for being N_2_O sinks cannot be predicted by their genomes. The phenotyping of the isolates revealed conspicuous patterns of *bet hedging* as demonstrated for *Paracoccus denitricans* (Lycus et al 2018). The *bet hedging* in *P. denitrificans* is characterized by expression of Nir (and Nor) in a minority of the cells, while Nos is expressed in all cells, in response to oxygen depletion, hence the population as a whole is a strong sink for N_2_O. The isolated *Pseudomonas* sp. (PS) displayed denitrification kinetics that closely resembles that of *P. denitrificans*. The two other isolates (**AN** and **AS**) showed indications of *bet hedging* as well, but of another sort: Nap appears to be expressed in a minority of the cells. This different regulatory phenotype had clear implications for the ability of organisms to function as N_2_O-sinks: while all isolates were strong N_2_O sinks when provided with NO_3_-only, **AN** and **AS** accumulated large amounts of N_2_O if provided with NO_2_^−^.

The N_2_O sink capacity of the organisms was tested by fertilizing soils with digestates with and without the organisms, and monitoring the gas kinetics in response to oxygen depletion, thus imitating the hot spots/hot moments of hypoxia/anoxia induced by digestates in soil (Kuzyakov and Blagodatskaya 2015). Since the isolates were raised by aerobic growth in autoclaved digestates, they would have to synthesize all denitrification enzymes in the soil, hence the synthesis of functional Nos was expected to be hampered by low pH (Liu et al 2014). The results for isolates **AS** and **AN** lend support to this (high ***I_N2O_*** in the soil with pH=5.5). **AN** was also dominating in the digestate enrichment culture, and in this case the organism had a strong and pH-independent effect on N_2_O emission, plausibly due synthesis of Nos prior to incorporation into the soils.

In summary, we have demonstrated that a digestate from biogas production can be transformed into an effective agent for mitigating N_2_O emission from soil, simply by allowing the right bacteria to grow to high cell densities in the digestate prior to fertilization. The technique is attractive because it can be integrated in existing biogas production systems, and hence is scalable. If we manage to treat a major part of waste materials in agroecosystems by AD, the resulting digestates would suffice to treat a large share of total farmland, as illustrated by **Figure 1**. Estimation of the potential N_2_O-mitigation effect is premature, but the documented feasibility and the scalability of the approach warrant further refinement as well as rigorous testing under field condition. Our approach suggests one avenue for a much needed valorization of organic wastes (Peng and Pivato 2019) via anaerobic digestion. Future developments of this approach could extend beyond the scope of climate change mitigation and include the enrichment of microbes for pesticide- and other organic pollutant degradation (Sun et al 2018), plant growth promotion (Backer et al 2018) and inoculation of other plant symbiotic bacteria (Poole et al 2018).

## Footnotes

* Reviewer access: Username. Password: GdTR3biE

** Reviewers access: Username: Password: nMz62S8O

