## Supplementary material for "N_2_O-respiring bacteria in biogas digestates for reduced agricultural emissions": Suppl Methods (from another version of this preprint: version 2, 2021-05-11)

#### 1. Digestates

The digestate material used in this study originated from mesophilic (37 $^{o}$C) and thermophilic (52 $^{o}$C) anaerobic digesters operated semi continuously and in parallel at a 750 000-person equivalent wastewater treatment plant (WWTP) (Oslo, Norway). The total individual reactor volume was 6 000 $m^3$ with a normal operation level of 5000 ± 300 $m^3$ of digestate. Mixing/stirring was maintained by intermittent recirculation of produced biogas (average 150 $Nm^3/h$) through lances releasing gas at the bottom of the reactor, and by continuous recirculation of digested sludge (216 – 432 $m^3/h$) from the bottom and back to the top. The substrate (raw sludge), top fed to the digesters, was a poly aluminum chloride (PAX-XL61™, Kemira) and ferric chloride (PIX318™, Kemira) precipitated municipal wastewater sludge, dewatered, by addition of 1.6 ± 0.5 kg $ton^{-1}$ total solids cationic polyacrylamide based polymer flocculant (Zetag 7550®, Kemetyl) by decantation of free water through rotary drum filters, and buffered in a stirred holding tank (retention time ~24 hours) prior to anaerobic digestion. The total solids content (TS %) of the raw sludge was 7.1 ± 0.5 %, and loss of ignition (LOI, % of dry weight) was 79 ± 3 % (measured by the WWTP, given as yearly average). Yearly average operational parameters of the digesters were provided by the WWTP and are shown in Table 1.

Chemical and physical properties of the digestates, the slurry of anaerobically digested wastewater sludge, were analyzed in the NS-EN ISO/IEC 17025 accredited laboratory belonging to the WWTP. Total alkalinity, pH and total volatile fatty acid (VFA) concentration in the digestates were determined using a two-point titration procedure described in EN12176:1998. Total solids (TS %) and volatile solids (VS %) were determined according to EN15934 and EN15935, respectively. The sum of $NH_3$ and $NH_4^+$ was measured as described by Greenberg et al (1980), using a ThermoOrion Model 95-12 ammonia electrode. Instrument drifting and reproducibility were controlled by carrying out parallel measurements on reference materials.

Yearly average digestate characteristics were provided by the WWTP and are shown in Table 1 together with corresponding digestate and operational characteristics at the time of sampling for the digestate material used for enrichment culturing.

**Table 1** Operational parameters for the mesophilic and the thermophilic anaerobic digesters and digestate characteristics at the time of sampling for enrichment culturing. Enrichment culturing was repeated several times with digestate from the mesophilic AD, each with freshly sampled digestate (Sampling 1-7).

| | **Digestate characteristics** | | | | | | | **AD operational parameters** | | |
|---|---|---|---|---|---|---|---|---|---|---|
| | pH | % dry weight [a] | LOI [b] (% of DW) | TAK [c] (meq $L^{-1}$) | VFA [c] (meq $L^{-1}$) | VFA/TAK | $NH_3$+$NH_4^+$ (mg-N $L^{-1}$) | $CH_4$ prod. [d] rate (mmol $L^{-1}$ $h^{-1}$) | HRT [e] (d) | VS loading rate[f] (kgVS $m^{-3}d^{-1}$) |
| **Mesophilic** (WWTP average): | **7.6** | **3.84** | **54.2** | **185** | **16.1** | **0.087** | **1824** | **1.5** | **24.4** | **2.4** |
| Sample 1[g] | 7.6 | 3.85 | 55.8 | 188 | 16.0 | 0.085 | n.d. | 1.48 | 17.2 | 3.3 |
| Sample 2[g] | 7.7 | 3.81 | 57.0 | 187 | 15.2 | 0.081 | n.d. | 1.55 | 22.0 | 2.4 |
| Sample 3[g] | 7.7 | 3.81 | 57.0 | 187 | 15.2 | 0.081 | n.d. | 1.55 | 22.0 | 2.4 |
| Sample 4[g] | 7.8 | 3.79 | 58.1 | 184 | 17.1 | 0.082 | n.d. | 1.21 | 28.2 | 1.9 |
| Sample 5[g] | 7.6 | 3.70 | 56.1 | 188 | 15.1 | 0.080 | n.d. | 1.25 | 30.8 | 1.9 |
| Sample 6[g] | 7.8 | 3.91 | 56.4 | 184 | 16.6 | 0.084 | n.d. | 1.36 | 21.1 | 2.1 |
| Sample 7[g] | 7.6 | 3.69 | 57.6 | 198 | 16.3 | 0.082 | n.d | 1.24 | 25.4 | 2.1 |
| **Thermophilic** (WWTP average): | **8.1** | **3.79** | **54.7** | **207** | **31.2** | **0.150** | **1922** | **1.64** | **22.8** | **2.5** |
| Sample[h] | 8.2 | 3.74 | 52.1 | 237 | 36.9 | 0.156 | n.d. | 1.61 | 17.3 | 3.3 |

[a] Dry weight % expressed as percentage of wet weight.
[b] Loss of ignition as percentage of dry weight.
[c] VFA = volatile fatty acids. TAK = total alkalinity.
[d] $m^3$ gas (1 bar, 0 $^0$C), 45% $CO_2$, 55% $CH_4$.
[e] Hydraulic retention time (days)
[f] VS = volatile solids= fraction of organic material, determined by ignition (LOI)
[g] Sample 1 (date: 2017.04.26) was used for the enrichment analyzed by genomics and proteomics (Figure 2A), Sample 2 (date 2017.12.12): repeated experiment shown in Figure 2B&C, Sample 3 (date 2018.05.01): third repeat of the enrichment (Figure S2), Sample 4 (date 2020.05.02): oxic cultivation of isolates (Figure S25, S26), Samples 5 and 6 (dates:2020.05.05, 2020.05.15): digestate used for soil inoculation (Figures S27, S28, S29), Sample 7 (date2020.08.26): final enrichment where $H_2$ was monitored (Figure S13).
[h] Thermophilic digestate was only used in the first enrichment experiment (Figure S6 and S7)

Samples of digestates for enrichment culturing were taken from the mesophilic and thermophilic anaerobic digestors at sampling points located on the recirculation loop of the digesters. The digestates were transported to the laboratory in 1 L vacuum isolated steel vessels which were filled completely to minimize the exposure to $O_2$ and used in enrichment culturing within 3-6 hours after sampling. During the 3-6 hours between sampling and initiation of enrichment culturing, the temperature in the thermos fell to ~20 °C. For each enrichment culturing, we took new samples, and the operational parameters and digestate characteristics for each case are shown in Table 1. Raw wastewater sludge was used in some experiment, and this was taken downstream of the buffer tank (see above) and transported in 1 L steel vessels a described for digestates.

### 2 Incubation- and gas measurement system, calculation of concentrations and rates of transformations.

In all incubations described below, we used a temperature controlled robotized incubation system, as described by Molstad et al (2007, 2016). This system samples the headspace of 120 mL closed vials at intervals, and analyses $O_2$, $N_2$, $N_2O$, NO, $CO_2$ and $CH_4$ in a single gas sample, using a gas chromatograph (789A GC-System, Agilent Technologies) and a chemiluminescence NO analyzer (Model 200A, Teledyne Instruments). The sampled gas is replaced by an equal volume of He. Mass loss due to sampling, and leakage of $N_2$ and $O_2$ through tubing, valves and septa are accounted for when estimating the rates of production/consumption for each time increment between two samplings. The measured gas concentrations in the headspace can be converted to concentrations in the liquid, based on the solubility of the individual gases and the empirically determined transport coefficient for gas exchange between the headspace and the liquid (explained in detail by Molstad et al 2007). The concentration of $N_2O$ is reported as mol $N_2O$ $L^{-1}$ in the liquid, rather than the concentration in the headspace, since the concentration in the liquid is what the organisms experience. The amounts of $N_2$ produced (and $N_2O$ reduced) are expressed either as mol $N_2$ and $N_2O$, or as mol N ($N_2$-N and $N_2O$-N). The latter is a convention in denitrification research which simplifies nitrogen mass balance calculations. An example excel spreadsheet with dummy data, but otherwise identical to the ones used in this work, transparent with respect to all calculations regarding gas kinetics and solubility of gases, in addition to accompanying e-learning videos, is available (see Bakken 2020).

**3. Enrichment culturing and samples for molecular analyzes and VFA quantification**

Within 3-6 hours after sampling digestates from the WWTP (Table 1) triplicates of 50 mL mesophilic, thermophilic and a heat treated mesophilic digestate (heat treated at 55 °C for two hours in a temperature controlled water bath) were transferred to 120 mL glass vials with a 23 mm Teflon coated triangular magnet. The vials were crimp sealed with a butyl rubber septum, and headspace air was removed and replaced by helium by repeated evacuation and He-filling (“He-washing”, see Molstad et al 2007). The procedure of filling and helium washing took ~1 hour. The vials were then placed on a magnetic stirring plate (stirring speed 300 rpm) in the thermostatic water-bath (20 $^{o}$C) of the incubation robot and the He overpressure was released after temperature equilibration with the water bath (~10 minutes). Then 3 mL medical grade $N_2O$ (Aga, Norway) was injected to the vials, and the gas kinetics was monitored by frequent sampling of the headspace. Additional $N_2O$ was injected several times throughout the incubation, in response to depletion. Negative controls without injection of $N_2O$ were included.

Samples for metagenomics and metaproteomics were taken at three time points during the enrichment culturing, using a syringe flushed with helium to minimize oxygen contamination. The samples were placed in an ultra-freezer (-80 °C) immediately after sampling. The first samples (1 mL vial$^{-1}$, sample name “0h” used throughout) were taken prior to the first injection of $N_2O$. Subsequent samples were taken after 115 h (0.2 mL vial$^{-1}$) sample name “115h” used throughout the text), and at the end of the incubation (t = 325 hours, 1 mL vial$^{-1}$) sample name “325h” used throughout the text).

This enrichment culturing experiment was repeated several times (each time using freshly sampled digestate, Table 1) to check reproducibility of the gas kinetics, while metagenomic and metaproteomic analyses were done only in the first experiment (Figure 2A). The repeated enrichment experiments were done to refine the analyses of the gas kinetics, and to explore inhibition of methanogenesis by $N_2O$. In addition, we incubated mesophilic digestates provided with either $NO_3^-$ or $O_2$ (no $N_2O$) to assess the potential for $NO_3^-$ and $O_2$- consumption, and the effects of these electron acceptors on methanogenesis. In the final enrichment culturing, we used an improved version of the incubation robot, equipped with an extra detector (Plasma Emission Detector, LDetek) for quantification of $H_2$, thus testing if the inhibition of methanogenesis by $N_2O$ resulted in $H_2$ accumulation.

We also conducted $N_2O$ enrichments with mesophilic digestates (fresh as well as heat treated) amended with raw sludge (1 mL in 50 mL digestate per vial) to assess the potential for $N_2O$ reduction in raw sludge versus the digestate, hence implicitly assessing to which degree $N_2O$-reducing organisms in the raw sludge survive in the digester.

*DNA extraction and quantification of 16S copy numbers*

The samples taken from the enrichment culturing from mesophilic vials (after 0, 115 and 325 hours) and from heat treated mesophilic vials (0h and 325h) were thawed at room temperature and centrifuged at 10 000 × g for 3 minutes. DNA extraction from the resulting pellet was performed using PowerLyzer® PowerSoil® DNA Isolation kit (QIAGEN) following the manufacturer's protocol. Extracted DNA was stored at -20 °C prior to high throughput metagenome sequencing (see *Metagenomics* paragraph below) and quantification of the 16S gene copy number with quantitative digital droplet PCR (ddPCR). ddPCR was performed on technical triplicates of DNA preparations from each mesophilic sample. The ddPCR reaction mix was prepared according to the manufacturer's instructions. Each sample contained 10 µL QX200 ddPCR EvaGreen Supermix (Bio-Rad), 2 µL of DNA template, and 100 nM final concentration of the universal primer-pair PRK341F (5'-CCTACGGGRBGCASCAG-3') and PRK806R (5'-GGACTACYVGGGTATCT-3') (Eurofins Genomic) targeting the V3-V4 region of 16S rDNA (Yu et al 2005). Oil droplets where generated in a QX200 droplet generator from 20 µL reaction mix and 70 µL droplet generation oil for EvaGreen (Bio-Rad) and 40 µL of the oil droplet suspension was transferred to a well of a 96 well twin.tec plate (Eppendorf) that was heat sealed with aluminum foil (PX1™ PCR plate sealer (Bio-Rad)). The PCR reaction was conducted in a 2720 Thermal Cycler (Applied Biosystems) with 2 °C $s^{-1}$ ramp rate, a lid temperature of 105 °C, and ran for 40 cycles, as recommended by the supplier, with temperature settings: 95 °C for 30 seconds (denaturation), 55 °C for 30 seconds (annealing) and 45 seconds at 72 °C (extension). The last cycle was followed by 5 minutes at 4 °C and 5 minutes at 90 °C (for signal stabilization). PCR products where analyzed in a QX200 droplet reader (Bio-Rad), and the data was analyzed using the Quantasoft™ Analysis Pro 1.0.596 software (Bio-Rad).

*VFA quantification*

Samples taken during enrichment culturing (sample: 0h, 115h and 325h) were stored frozen (-80$^{o}$C) until analyzed for VFA. In addition, we analyzed VFA in freshly sampled, i.e. digestate that was frozen (-80 °C) immediately after sampling, from the anaerobic digester. The frozen samples were thawed in room temperature and centrifuged at 12 000 × g for 5 minutes.

Thereafter the supernatant was pH adjusted to ~2.5 using concentrated $H_2SO_4$ and centrifugated for 1 minute at 12 000 × g. The supernatant of individual samples was divided in three aliquots (technical triplicates). Quantification of VFAs (formate, acetate, propionate, iso-butyrate, valerate and iso-valerate) was done with high pressure liquid chromatography (HPLC) using a Dionex Ultimate 3000 system (Dionex, USA), operated at 40 °C with flowrate 0.3 mL $min^{-1}$, equipped with a UV detector (210 nm) and a Zorbax Eclipse Plus C18 column (Agilent, USA) (150 x 2.1 mm, 3.5 µm particles) and a guard column (12.5 x 2.1 mm; 5 µm particles) (Agilent, USA) for all samples. Standards covered the range (mM); 0.8–330, 0.5–220, 0.5–170, 0.7–135, 0.4–135, 0.4– 113 and 0.3-112 for formate, acetate, propionate, iso-butyrate, valerate and iso-valerate, respectively. The sample volume was 1 µL. Separation was achieved by applying a gradient of 2.5 µM $H_2SO_4$ and methanol as outlined in Table 2.

**Table 2** Elution profile, VFA quantification.

| Time (min) | Methanol (anhydrous) (%) | 2.5 mM $H_2SO_4$ (%) |
|---|---|---|
| (0.0 – 2.5) | 0 | 100 |
| (2.5 – 25) | 15 | 85 |
| (25 – 35) | 0 | 100 |

*Modelling growth of $N_2O$ reducing bacteria based on the measured $N_2O$ kinetics*

Growth of $N_2O$-respiring organisms in the enrichment cultures was estimated from the measured kinetics of $N_2O$ reduction, using parameters for anaerobic growth of the model denitrifying bacterium *Paracoccus denitrificans* as determined in our laboratory (Bergaust et al 2010, 2012): Cell dry-weight = 310 (+/-50) fg $cell^{-1}$, growth yield, $Y_{e-}$ = 1.9 · $10^{13}$ cells $mol^{-1}$ electrons to $N_2O$ (= 5.7 g cell dry-weight $mol^{-1}$ $e^-$), growth rate = 0.1 $h^{-1}$ (the cell specific rate of electron flow at this growth rate, $V_{emax}$, = 5.26 · $10^{-15}$ mol $e^-$ $h^{-1}$ ). These parameters were determined in experiments with succinate as the sole C source and at 20 °C (i.e. the same temperature as in all enrichment cultivations). Details of the modelling are explained in the legend of **Figure S2**. It should be noted that the calculated cell numbers are expressed as "*Paracoccus* equivalents" i.e. cells with 310 · $10^{-15}$ g dry weight $cell^{-1}$. The estimated cell densities can be converted to cell dry weights $mL^{-1}$ with reasonable confidence because different denitrifying organisms have fairly similar growth yields in terms of g dry weight $mol^{-1}$ $e^-$ to $N_2O$, namely 4-6 g cell dry weight $mol^{-1}$ $e^-$ (Hein et al 2017, Yoon et al 2016).

### 4 Metagenomics

*Metagenomics*

Isolated DNA from the mesophilic enrichment (samples 0h, 115h and 325h) was sequenced on an Illumina HiSeq4000 system, using TruSeq PCR-free library preparation. This also included DNA from the parallel enrichment with a pre-heated (55 °C for 2 hours) mesophilic digestate (Figure S6), to improve downstream analysis (i.e. binning) by increasing the differential coverage (Albertsen et al 2013). All reads were trimmed using Trimmomatic v0.36 (Bolger et al 2014) in pair end mode (Bolger et al 2014), before assembly with metaSPADes v3.10.1 (Nurk et al 2017). Both individual assemblies and co-assemblies of all samples from the enrichment were carried out, of which the co-assemblies were evaluated to give a better result according to metaQuast v4.5 (Mikheenko et al 2016). Metagenome assembled genomes (MAGs) was recovered from the co-assemblies (contigs > 500 bp) using MaxBin2 v2.2.1 (Yu-Wei et al 2016), and the quality of the MAGs was evaluated using CheckM v1.0.13 (Parks et al 2014). The binning effort resulted in 278 MAGs, of which 149 were considered to be of sufficient quality (completeness >50 %, contamination <20 %) for downstream analysis (**Supplementary Data S1**). Gene calling and functional annotation of the metagenomes were carried out using Prodigal v2.6.1 (Hyatt et al 2010) and InterProScan5 v5.32-71.0 (Jones et al 2014). Raw reads from each experimental sample were mapped to a concatenated fasta file of the 149 MAGs (Bowtie2 v2.3.4.1 (Langmead and Salzberg 2012) and Samtools v1.3.1) (Li et al 2009) and the relative abundance of each MAG was calculated using CoverM v0.3.2 (https://github.com/wwood/CoverM) requiring a minimum read identity of 95% and minimum read alignment of 75%.

*Phylogenetic placement and taxonomic classification*

A set of 16 universal single-copy ribosomal proteins (L2-L6, L14-L16, L18, L22, L24, S3, S8, S10, S17 and S18) was used to build a phylogenetic tree consisting of the 149 MAGs. The ribosomal proteins were identified within the functionally annotated MAGs. Four MAGs lacked all 16 protein sequences, and these were excluded from the phylogenetic tree but included in the taxonomic classification described below, which analyzes 100+ marker genes. Separate alignments were built for every ribosomal protein using MUSCLE v3.8.31 (Edgar 2004). The alignments were manually checked for misalignments and all conserved, single-copy ribosomal protein sequences that occurred more than once in a MAG were excluded GBlocks (Castresana 2000, Talavera and Castresana 2007) (parameters: -b2=50, -b3=20, -b4=2) was used to find conserved regions in each alignment, and the aligned regions for single proteins were then concatenated in an alignment of 2528 residues. Maximum likelihood phylogenies were built with RAxML-ng (Stamatakis 2014) using the PROTGAMMAWAG method, and the consensus tree was visualized using iTOL (Letunic and Bork 2019). A complete version of the tree is available in Newick format as Supplementary Data S3. The taxonomic classification of the MAGs was inferred using a set of 120 bacterial and 122 archaeal marker genes via the Genome Taxonomy Database GTDB v1.0.2 (Chaumeil et al 2019) using *classify_wf* with default parameters.

**5.Quantitative metaproteomics**

Samples were prepared by an initial centrifugation of mesophilic digestate samples in replicates (taken at 0, 115 and 325 hours) to separate the fiber fraction from the secretome. The secretome was filtered with a 0.22 µm sterile filter to remove cells and debris and treated with TCA (10 % final concentration) to precipitate the proteins. The fiber fraction was resuspended in dissociation buffer (1 % methanol, 1 % tert-butanol, 0.1 % Tween-80, pH 2) and, after gently mixed for 30 s at room temperature, released material (including cells) was separated from the plant material via a gentle spin (100 × *g*, 30 sec) and the supernatant retained in a fresh tube (Frank et al 2016). This procedure was repeated three times to increase the yield. Cell lysates were prepared by bead-beating in lysis buffer (50 mM tris-HCl,

200 mM NaCl, 1 mM DTT, 0.1% Trition X-100, pH 7.5), using glass beads (diameter ≤ 106 µm), followed by centrifugation (16.000 × *g*, 15 minutes) to spin down beads and cellular debris. The proteins in the collected lysate were precipitated using TCA as above.

The TCA precipitated proteins were resuspended in Laemmli sample buffer and subjected to SDS-PAGE (270V, 4 minutes). Each sample (secretome, cell lysate) was excised from the gels in four fractions = gel pieces). After washing the gel pieces with 25 mM ammonium biocarbonate, pH 7.8 in 50% acetonitrile, proteins were reduced by incubation in 10 mM DTT for 30 minutes at 56 °C, followed by carbamidomethylation by incubation with 55 mM iodoacetamide for 30 minutes at room temperature. Subsequently, the proteins were digested into peptides using 300 ng trypsin per sample and incubation at 37 °C, overnight. The peptides were desalted using C18 ZipTips (Merch Millipore, Darmstadt, Germany), according to the manufacturer's instructions, and analysed by nanoLC-MS/MS using a Dionex Ultimate 3000 UHPLC (Thermo Scientific) coupled to a Q-Exactive hybrid quadupole orbitrap mass spectrometer (Thermo Scientific, Bremen, Germany). Peptides were separated using an analytical column (Acclaim PepMap RSLC C18, 2 µm, 100 Å, 75 µm i.d. × 50 cm, nanoViper) with a 90-minutes gradient from 3.2 to 44 % [v/v] acetonitrile in 0.1 % [v/v] formic acid) at flow rate 300 nL/min. The Q-Exactive mass spectrometer was operated in data-dependent mode acquiring one full scan (400-1500 m/z) at R=70000 followed by (up to) 10 dependent MS/MS scans at R=35000.

The acquired MS/MS spectra were searched against the proteome of the 149 MAGs recovered from the abovementioned metagenomics data (785 999 protein sequences), using MaxQuant version 1.6.3.3 (Cox and Mann 2008). Common contaminants, such as human keratins, trypsin and bovine serum albumin were concatenated to the sample specific database as well as reversed sequences of all protein entries for estimation of false discovery rates. Proteins were quantified using the MaxLFQ algorithm in MaxQuant (Cox et al 2014). Protein N-terminal acetylation and oxidation of methionine were used as variable modifications, while carbamidomethylation of cysteine residues was used as a fixed modification. Tolerance levels for peptide identifications were 4.5 ppm and 20 ppm for MS and MS/MS, respectively, and two missed cleavages of trypsin were allowed. Additional quality filtering and downstream interpretation were performed in the software platform Perseus version 1.6.0.7 (Tyanova et al 2016). This included removal of contaminations, hits to reversed sequences and hits based on a single modified peptide. Furthermore, all identifications were filtered in order to achieve a protein false discovery rate (FDR) of 1% using the target-decoy strategy. For a protein group to be considered valid, we required the protein group to have at least one unique peptide and be detected in at least two of the three replicates for samples taken at 0 and 325 hours. For the sample taken after 115 hours, only duplicates were available, in this case proteins were only considered valid if they were identified in both replicates, to preserve high confidence. The putative functionality of the detected protein groups was assigned using the abovementioned InterProScan annotation of the protein sequences in the database as well as with the dbCAN2 meta server (Zhang et al 2018) (using CAZy-HMMs version 8) to detect putative carbohydrate-active enzymes.

To construct metaproteome-based metabolic maps (**Figure S12**), we scanned the detected proteins affiliated to each MAG (**Supplementary Data S2**) for enzymes involved in specific metabolic pathways (Frank et al 2015) In brief, to predict that a given population utilized monomeric sugars, we detected genes associated with the Embden-Meyerhof-Parnas pathway (glycolysis), including phosphofructokinases. Gluconeogenesis was predicted in a given population if a representative unidirectional fructose diphosphatase gene was detected in the proteome. The detection of both a phosphate acetyl/butyryl transferase-enzyme (contains phosphotransacetylase) and an acetokinase, or Acetyl-CoA hydrolase/transferase, was used to predict acetate metabolism. For predicting the ability of given populations to oxidize fatty acids, we detected key proteins encoded on a methylmalonyl-CoA (MMC) gene cluster and all four enzymes inferred in beta-oxidation (acyl-CoA dehydrogenase, enoyl-CoA hydratase, hydroxy acyl-CoA dehydrogenase, and ketoacyl-CoA thiolase). The prediction of an active Wood-Ljungdahl pathway was assessed as follows: the combination of a highly expressed CO dehydrogenase/acetyl-CoA synthase cluster, electron transfer complex, aldehyde ferredoxin oxidoreductase for potential acetate activation, and enzymes for beta-oxidation of fatty acids. The methanogen-affiliated proteomes were scanned for the detection of Coenzyme M methyl-transferase, a key enzyme in the methanogenesis. Nos (EC:1.7.2.4) levels were visualized using ggplot2 in RStudio. The proteome size of individual MAGs was estimated in two different manner, either by the number of proteins detected per MAG or as the sum of LFQ-values for all proteins belonging to one MAG normalized to the total LFQ for the sample.

### 6. Isolation of $N_2O$ reducing bacteria

Three types of media were used for incubation of cultures in liquid medium or on agar plates (1.5 w. % agar). Unless otherwise stated, the media were brought to desired strength by diluting stock solutions or dry powders in milliQ $H_2O$, pH adjusted by addition of KOH/HCl to pH = 7.0 and autoclaved at 121 °C for 20 minutes. Sistrom's succinate medium (**SS**), contained ($L^{-1}$) 3.48 g $K_2HPO_4$, 0.195 g $NH_4Cl$, 4 g succinic acid, 0.10 g glutamic acid, 0.04 g aspartic acid, 0.5 g NaCl, 0.2 g nitrolotriacetic acid, 0.3 g $MgSO_4 \cdot 7H_2O$, 0.015 g $CaCl_2 \cdot 7H_2O$, 0.002 g $FeSO_2 \cdot 7H_2O$, 0.1 mL trace element solution and 0.1 mL vitamin solution. The trace element solution contained (g $L^{-1}$): 17.65 g EDTA (triplex 3), 109.5 g $ZnSO_4 \cdot 7H_2O$, 50 g $FeSO_4 \cdot 7H_2O$, 15.4 g $MnSO_4 \cdot H_2O$, 3.92 g $CuSO_4 \cdot 5H_2O$, 2.48 g $Co(NO_3)_2 \cdot 6H_2O$ and 1.14 g $H_3BO_3$; $H_2SO_4$ was added until the solution cleared. The vitamin solution contained (g $L^{-1}$) 10.0 g nicotinic acid, 5.0 g thiamine HCl and 0.10 g Biotin. Digestate medium (**D**) was prepared by centrifuging digestate from the VEAS WWTP at 8000 × g for 30 minutes, after which the supernatant was pH adjusted to ~6.5 and autoclaved (121 °C for 20 minutes). The heat treatment led to loss of dissolved $CO_2$ and a pH increase, giving a final pH of 7.5. Anaerobe basal medium (**AB**; OXOID CM0957, Thermo Scientific) contained ($L^{-1}$) 1.6 g peptone, 0.7 g yeast extract, 0.5 g sodium chloride, 0.1 g starch, 0.1 g arginine, 0.05 g sodium succinate, 0.05 g L-cysteine hydrochloride, 0.04 g sodium bicarbonate, 0.05 g ferric pyrophosphate, 0.01 g dithiothreitol, 0.05 g sodium thioglycolate, 0.0005 g haemin and 0.00004 g vitamin K.

Dilution series of dispersed $N_2O$ enriched mesophilic digestate were prepared and spread on agar plates (50 µL diluted suspension per plate) on all media (SS, D and AB) shortly after ending the enrichment culturing. In order to select for $N_2O$ reducing strains the agar plates were incubated anoxically in 8.6 L anaerobe boxes which were first sparged with $N_2$, followed by injecting ~8 vol % $N_2O$. To secure anoxic conditions anaerobic boxes were equipped with two oxygen scavenger bags (3.5 L AnaeroGen™, OXOID). The plates incubated at 20 °C and inspected after ~2 weeks, and a selection of visible colonies were picked and re-streaked on daughter plates with corresponding media and incubated anaerobically with $N_2O$ as described. Growing colonies were picked and re-streaked on corresponding plates and incubated under aerobic conditions to avoid continuation of growth of obligate fermentative bacteria. The 16S gene of single colonies growing on aerobic plates was amplified by PCR using the DreamTaq™ Green PCR Master Mix (Thermo Scientific) using the bacteria specific primer 27F (3'-AGAGTTTGATCMTGGCTCAG-5') (Lane 1991, Invitrogen)  and the universal primer 1492R  (5'-GGTTACCTTGTTACGACTT-3') (Stackerbrandt and Liesack 1993, Invitrogen) in a 2720 Thermal Cycler (Applied Biosystems) with 2 °C $s^{-1}$ ramp rate and a lid temperature of 105 °C. The temperature parameters of the 30 amplification cycles were 98 °C for 10 seconds, 55 °C for 30 seconds and 72 °C for 1 minute. The last cycle was followed by a 1-minute final elongation at 72 °C and a 4 °C hold step. The presence of contaminants was evaluated based on inspection of Sanger sequencing chromatograms of the 16S PCR amplicons (LightRUN™ sequencing services, Eurofins Genomics, Germany) and by inspection/microscopy of colony- and cell morphology. The 16S analyses showed that a large majority of the SS-, D- and AB agar plates had growing colonies related to *Azonexus* sp..

One colony of *Azonexus* sp., growing on SS agar, was selected for further work and was given the working name "**AN**". Another culture, related to *Pseudomonas sp*., growing on SS-agar, was obtained and given the working name "**PS**". Continuation of aerobic growth of AN on new SS agar plates revealed a minor contamination (contaminant was not visible in Sanger sequencing chromatograms of 16S amplicons obtained of the mother colony). Re-streaking and purification of the contaminated culture revealed that the contaminant had almost identical morphological features as **AN** when growing as single colonies, and when inspected under the light microscope. The contaminant, related to *Azospira* sp. by 16S, was given the working name "**AS**".

The cultures of **AS** and **PS** where grown aerobically at 20 °C in stirred (700 rpm) SS liquid media to $OD_{660nm}$ ~ 1 (UV-1280 UV-VIS spectrophotometer, Shimadzu), and aliquots were snap frozen as glycerol stocks (15 wt. %) in liquid nitrogen and stored as precultures at -80 °C. **AN** was not revivable after freezing, and was kept as $N_2O$ raised colonies on SS agar slabs stored at 4 °C.

**7. Genome sequencing of isolates and comparison average nucleotide identity (ANI) with metagenome-assembled genomes**

The isolates were recovered from snap frozen glycerol (15 vol%) stocks of aerobically grown cultures in SS liquid medium (AS and PS), or from single colonies grown on $N_2O$ on SS-agar slabs stored at 4 °C (AN), and grown aerobically (stirred at 700 rpm, air atmosphere) at 20 °C in SS medium to late exponential phase ($OD_{660} \approx 1.0$). The cell suspensions were centrifuged at 10000 × *g* for 10 minutes and DNA was extracted from the pellet using the PowerLyzer™ Soil DNA extraction kit (QIAGEN) following a modified kit protocol (bead beating for 45 s at 4.5 $m.s^{-1}$ in a FastPrep®-24 (M.P. Biomedicals) substituted the vortexing step in the manufacturers protocol). Paired end MiSeq sequencing on extracted DNA was performed at the Norwegian Sequencing Center on a MiSeq v2 nano 250 PE platform with Nextera™ DNA Flex Tagmentation sample preparation for the isolates AN and AS. PS was sequenced at Novogene Co., Ltd., Hongkong on a HiSeq4000 platform 150 PE. Raw reads were quality checked with FastQC v0.11.5 (Andrews, 2010). Removal of low-quality sequences and ambiguous reads was done using Trimmomatic (Bolger et al 2014) with the following settings: sliding window 4:15; adapter clipping options: enabled for adapters NexteraPE-PE (for AN and AP only); seed mismatches 2; palindrome clip threshold 30; simple clip threshold 10; head crop length 12 (AN and AP only). Contig assembly was done with SPAdes (Nurk et al 2013) using default parameters. Quality assessment of the assembled contigs was done in Quast (Gurevich et al 2014) with the following settings: unaligned part size 1000; extensive mix size 1000; min alignment 50; min identity 80. Prokka v1.12 (Seemann 2014) and RAST (Aziz et al 2008) were used for annotation of the assembled contigs with default parameters. The OrthoANIu tool (Yoon et al 2017) was used to compare and to calculate average nucleotide identities the sequenced genomes and the metagenome assembled genomes.

### 8. Phenotyping of isolates

The capacity of the isolates to utilize a variety of carbon substrates was tested using PM1 and PM2 BiOLOG Phenotype MicroArray™ plates (BiOLOG Inc. Hayward, CA). The BiOLOG test method is based on the irreversible reduction of tetrazolium violet to formazan as in indicator of active metabolism (Bochner et al 2001). The isolates were raised on Merck Nutrient broth agar plates (20 g agar $L^{-1}$) and transferred to the BiOLOG plates according to the instructions of the manufacturer. The plates were incubated at 30 °C and analyzed by spectrophotometry after 72 hours. The experiments included control plates without inoculum, and 3 replicate plates for each isolate.

The characteristic regulation of denitrification (regulatory phenotypes) by the isolated cultures was determined as in previous investigations (Bergaust et al 2011, Liu et al 2013, Lycus et al 2018, Mania et al 2016) by monitoring the kinetics of $O_2$, $N_2$, $N_2O$, NO and $CO_2$ throughout the cultures' depletion of $O_2$ and transition from aerobic to anaerobic respiration in stirred batch cultures with He + $O_2$ (+/- $N_2O$) in the headspace.

The cells to inoculate these vials were raised under strict aerobic conditions to avoid synthesis of denitrification enzymes prior to inoculation for testing the regulatory phenotypes: 1 mL frozen pre-culture of the isolates **AS** and **PS**, and cells from a single colony of **AN** (this culture did not survive freezing), were raised in 50 mL liquid SS medium (initial $OD_{660nm}$ of ~0.02 for **AS** and **PS** and <0.01 for **AN**) under oxic conditions (air, the 120 mL serum vials covered with Al-foil) at 20 °C with rapid stirring (700 rpm). When $OD_{660nm}$ reached ~0.2, 1 mL was transferred to new vials containing 50 mL SS medium for continuation of aerobic growth.

When the cultures reached $OD_{660nm}$ 0.05 – 0.1, they were used to inoculate the phenotype test vials.

The phenotyping was conducted in triplicate or duplicate 120 mL capped vials containing 50 mL SS medium supplemented with either $NO_2^-$ (1 mM, 50 µmol), $NO_3^-$ (2 mM, 100 µmol) or both. The headspace (He) was supplemented either with 1 mL $O_2$, or 1 mL $O_2$ + 1 mL $N_2O$. The vials were inoculated with 0.1 – 0.5 mL (depending of the OD) of the aerobically raised cultures (added using a sterile syringe) and monitored for gas kinetics while incubated at 20 $^oC$ (stirred, 700 rpm).

Nitrite concentrations were measured at various timepoints throughout the incubations, by taking 10 µL liquid samples which were injected immediately into a purging device containing 1% w/v NaI in 50% acetic, which converts nitrite instantaneously to NO, which is transported (by $N_2$-flow) to a chemiluminescence NO analyzer (Sievers 280i, GE Analytical Instruments) (Cox 1980, MacArthur et al 2007.

**9. Protein extraction and quantitative proteomics in *Azonexus* sp. AN.**

*A* cell culture of *Azonexus* sp. AN was raised from a single colony incubated at aerobic conditions at 20 °C with rapid stirring (700 rpm) in 50 mL SS liquid medium. When the pre-culture reached an $OD_{660nm}$ of 0.2, 1 mL cell suspension was transferred to a new vial containing 50 mL Sistrom medium for continuation of aerobic growth at the same conditions. To determine the relative expression of $N_2O$ reductase and nitrate reductase in the isolate AN in response to the transition to anoxia, six sterile 120 mL vials containing 50 mL SS liquid medium supplemented with 2 mM $NO_3^-$ (0.1 mL 1M KNO3) and 1 mL $O_2$ in helium atmosphere (headspace volume 70 mL), was incubated at 20 °C and inoculated with 1 mL cell culture of *Azonexus* sp. AN ($OD_{660nm}$ = 0.150) using a sterile syringe. At intervals throughout the incubation, single vials were subjected to destructive sampling: the vial was removed from the incubation robot and immediately cooled down, with stirring, in ice-cold water. The entire culture volume was then transferred to a 50 mL sterile Falcon™ tube and centrifuged at 10 000 × *g* for 10 minutes at 4°C. The supernatant was gently poured of and the cell pellet immediately frozen ( -80 °C). The frozen cell pellets were thawed on ice, resuspended in lysis buffer (20 mM Tris-HCl pH 8.0, 0.1 % v/v Triton X-100, 200 mM NaCl, 1 mM DTT, 4% SDS) and treated with 3 × 45 s bead beating with glass beads (particle size ≤106 µm, Sigma) at maximum power and cooling on ice between the cycles (MP Biomedicals™ FastPrep- 24™, Thermo Fischer Scientific Inc). Cell debris was removed by centrifugation (10 000 × *g*; 5 min) and the supernatant, containing water soluble proteins, was used for proteome analysis using the NanoLC- Orbitrap-MS, as described above. Data analysis was performed in MaxQuant 1.6.2.3 (Cox and Mann 2008). The raw data was matched against the proteome of the type strain *Azonexus hydrophilus* 418702 (Uniprot, https://www.uniprot.org/proteomes/UP000187526) supplemented with sequences obtained from the predicted proteins of periplasmic nitrate reductase (Nap), nitrite reductase (Nir), nitric oxide reductase (Nor) and nitrous oxide reductase (Nos) obtained from the genome sequence of *Azonexus sp*. AN. The denitrification reductases were quantified by expressing their LFQ values. Since Nos is a homo-dimer and Nap a monomer, the LFQ values for Nos were divided by two to obtain the correct number of putatively functional enzymes.

**10. Incubations of soils for determining if digestates with $N_2O$-reducing bacteria can reduce the $N_2O$ emission from soil denitrification.**

Two agricultural clay loam soils (pH = 5.5 and 6.5) were used, taken from a long-term liming experiment at Ås, Norway (described by Nadeem et al 2020). Prior to incubations, the soils were sieved through a 3 mm metal mesh, air dried at room temperature, and stored in plastic containers at 4 °C for 4 months. The nitrate content of the two soils (when used for experiments) was 1.32 (± 0.01) and 1.13 (± 0.01) µmol $g^{-1}$ dry weight (standard error in parenthesis, n=3). The nitrite content of the soil was <5 nmol $g^{-1}$ (below detection limit).

The soils were inoculated with digestates that were pretreated in various ways to assess the effect of 1) the indigenous bacteria (in digestates as taken directly from the anaerobic digester), 2) indigenous $N_2O$-reducing bacteria enriched by anaerobic incubation with $N_2O$, and 3) isolated cultures grown aerobically in digestates. The digestates used were (working names in bold italics):

1) ***Live digestate*** = digestate directly from the anaerobic digester of the WWTP (sampled ~3 hours before inoculation of soils)

2) ***70 °C dig*** = Live digestate heated to 70°C for 2 h to kill most of the indigenous denitrifying bacteria in the live digestate

3) ***AN, AS*** and ***PS*** = autoclaved digestates in which isolated *Azonexus sp.*, *Azospira sp.* and *Pseudomonas sp.,* respectively, were grown by aerobic respiration.

4) **$N_2O$ enr.** = digestates in which $N_2O$-reducing bacteria were enriched by anaerobic cultivation with $N_2O$ (repeat of the enrichment culturing shown in Figure 2).

The procedure for aerobic cultivation of the isolated cultures in autoclaved digestate (point 3 above) was: freshly sampled digestate was autoclaved (121 °C, 20 min), which increased the pH to ~9.8 (due to removal of $CO_2$), and sparged with sterile filtered air for 24 hours. The air sparging was necessary because the WWTP adds ferric chloride as a precipitation chemical post anaerobic digestion, which is reduced to ferrous iron ($Fe^{2+}$) during AD (Cheng et al 2015): abiotic oxidation of $Fe^{2+}$ obscured measurements of oxygen consumption by respiration, and the abiotic oxidation may inhibit aerobic respiration due to formation of reactive oxygen species (Winterbourn 1995). After sparging, pH was adjusted to 7.5 by addition of HCl. The isolates where raised from frozen stocks (AS and PS), or from a single $N_2O$-raised colony picked from SS agar slabs stored at 4 °C (AN), in 50 mL SS medium at 20 °C under oxic conditions (air) with rapid stirring (700 rpm). At $OD_{660nm}$ ~0.2 the cultures where transferred to new vials containing 50 mL SS medium and growth continued under the same conditions, and when $OD_{660}$ reached ~1, 1 mL of each culture was added to 120 mL vials containing 50 mL autoclaved and the pre-aerated digestate, which were then incubated (stirred, 700 rpm) at 20 °C in the robotized incubation system. $O_2$ was injected several times throughout the incubation to maintain >10 vol % $O_2$ in the headspace and ended after 160 hours. At this time point, ***AN, AS*** and ***PS*** had consumed 16.3, 22.9 and 22.4 µmol $O_2$ $mL^{-1}$, respectively, which implies cell densities of 2.5-3.5*$10^9$ $mL^{-1}$, or 0.7-1 mg cell dry-weight $mL^{-1}$ , if assuming 300 fg dry weight $cell^{-1}$ and growth yield= 15*$10^{13}$ cells $mol^{-1}$ $O_2$, as determined for *Paracoccus denitrificans* (Bergaust et al 2010, 2012).

Digestate amendments of soils was set up as duplicate 120 mL vials with 10 g soil, amended with 3 mL digestate+ 0.1 mL 0.5 M $KNO_3$, which was spread as small droplets over the soil surface (~19 $cm^2$ surface area) using a syringe, resulting in ~61 % waterfilled pore space (bulk density = 1.1). The vials were then capped (butyl rubber septa), He-washed (repeated evacuation and He-filling), and 1 mL pure $O_2$ was injected with a syringe. The vials were then placed in the water-bath (20 $^0$C) of the incubation robot and monitored by frequent sampling of the headspace.

The $N_2O$ production index ($I_{N_2O}$) was calculated for each individual by

$$I_{N_2O} = \frac{\int_0^T \mathrm{N_2O{-}N(t)dt}}{\int_0^T [\mathrm{N_2O{-}N(t)} + \mathrm{N_2{-}N(t)} + \mathrm{NO(t)}]\mathrm{dt}} \qquad (1)$$

where $\int_0^T N_2O - N(t)dt$ is the area under the curve (trapezoidal rule) for measured $N_2O$-N (µ mol N $vial^{-1}$ h) and $\int_0^T [N_2O - N(t) + N_2 - N(t) + NO(t)]dt$ is the area under the curve for measured $N_2$+$N_2O$+NO-N (µ mol N $vial^{-1}$ h), both for the time period 0-T (h). $I_{N_2O}$ was calculated for two time periods: $I_{N_2O\ 40\%}$ is the index for the period (0-T) until 40% of the available $NO_3^-$- N was recovered as N-gas (NO+$N_2O$+$N_2$), $I_{N_2O\ 100\%}$ is the index for the time period (0-T) until 100% was recovered. $I_{N_2O}$ was used by Liu et al (2014) as a proxy for the relative propensity of a soil to emit $N_2O$ from denitrification, and its predictive capacity verified by Russenes et al (2016). The experiments included control treatment where distilled water replaced digestates (duplicate vials for both soils).
