## Supplementary for "N_2_O-respiring bacteria in biogas digestates for reduced agricultural emissions" (from another version of this preprint: version 2, 2021-05-11)

This Supplementary Online Material contains 7 sections:

| | Page |
|---|---|
| A. Gas kinetics during enrichment culturing | S2 |
| B. Growth and decline of members of the methanogenic consortium | S9 |
| C. Metabolism of the methanogenic consortium | S14 |
| D. Genetics of isolated organisms | S18 |
| E. Carbon substrate utilization by isolated organisms | S20 |
| F. Denitrification phenotypes of isolated organisms | S22 |
| G. Aerobic growth in sterilized digestate, and the effect of the enriched digestate on $N_2O$ emissions | S35 |
| H. References | S41 |

### A. Gas kinetics during enrichment culturing

This section provides details of the gas kinetics and modelled bacterial growth for anaerobic enrichment culturing with $N_2O$, as well as gas kinetics in additional (control) experiments with other electron acceptors and a different (thermophilic) digestate.

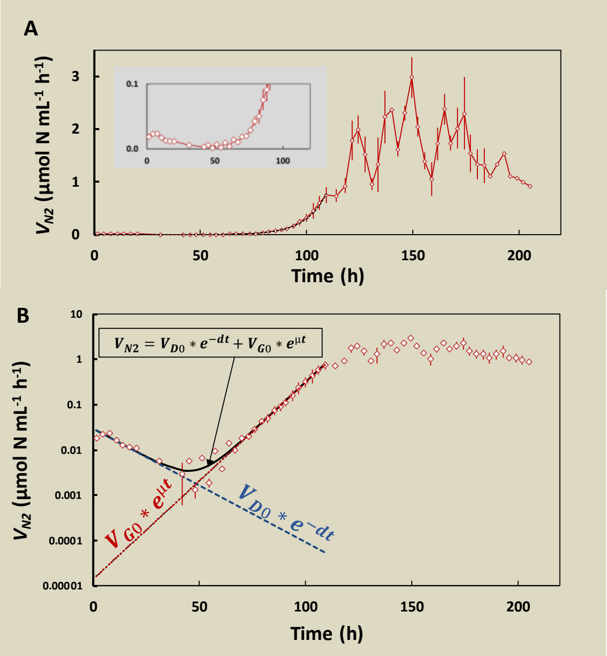


**Figure S1: Modelling growth of $N_2O$ reducing bacteria during enrichment by anaerobic incubation of a mesophilic digestate with $N_2O$, based on measured rates of $N_2O$ reduction to $N_2$.**

Panel A shows measured rates of $N_2O$ reduction to $N_2$ (***$V_{N2}$***, µmol N $mL^{-1}$ $h^{-1}$) in the second enrichment culture experiment (Figure 2BC). The values plotted are average of three replicate enrichment vials, with standard deviation as vertical lines. The insert is scaled to visualize the rates during the first 100 h. The fluctuations after 120 hours are due to episodes of $N_2O$ depletion and subsequent injections of $N_2O$ (see Figure 2B). Panel B shows the same data plotted on a log scale, illustrating that the rate declined exponentially during the first 30-40 h, and increased exponentially from 60 to 110 h. The kinetics suggested the presence of two groups of $N_2O$ respiring organisms: One whose respiration died out gradually during the enrichment (**D**), and one which was growing by respiring $N_2O$ (**G**). To assess the respiration kinetics of **D** and **G**, the following model was used:

$$V_{N2} = V_D + V_G = V_{D0} * e^{-dt} + V_{G0} * e^{\mu t}$$

where ***$V_D$*** is the rate of $N_2O$-reduction (µmol N $mL^{-1}$ $h^{-1}$) by **D**, **$V_{D0}$** is their rate at time zero, and ***d*** is the first order rate of decline ($h^{-1}$); ***$V_G$*** is the rate of $N_2O$-reduction by **G,** and ***µ*** is the growth rate of **G** ($h^{-1}$). The parameters were estimated by fitting the modeled to the measured data for 0-110 h, using the Generalized Reduced Gradient Solver in Excel, yielding the following results: ***$V_{D0}$***= 30 nmol N $mL^{-1}h^{-1}$, ***$V_{G0}$*** = 0.014 nmol N $mL^{-1}$ $h^{-1}$, ***d***= 0.03 $h^{-1}$, ***µ***=0.1 $h^{-1}$. The modelled **$V_{N2}$** (black line in panel B) fits very well with the data ($r^2$ = 0.997). The rates of $N_2O$ reduction by the growing and declining groups are shown as red and blue dotted lines.

The estimated growth rate of **G** (***µ***=0.1 $h^{-1}$) equals the maximum anaerobic growth rate of the model strain *Paracoccus denitrificans* at 20 $^0$C, as measured by Bergaust et al (2010, 2012), who also determined the growth yield (***Y*** =1.9*$10^{13}$ cells per mol electrons), and the cell specific electron flow rate at µ=0.1 $h^{-1}$ (***$V_{e\text{-}max}$***= 5.26*$10^{-15}$ mol $e^-$ $cell^{-1}$ $h^{-1}$). These parameters were used to estimate the number of "*Paracoccus* equivalent" cells in the enrichment culture as shown in Figure 2C in the main paper. For the period 0-110 hours with exponential growth, the cell density was calculated as ***$N_G(t)$***= ***$V_G(t)$/$V_{e\text{-}max}$*** where ***$N_G(t)$*** is the number of cells $mL^{-1}$ at time t (h) after initiation of the enrichment culturing, ***$V_G(t)$*** is the rate of $N_2O$-reduction (mol N $mL^{-1}$ $h^{-1}$) at time t, and ***$V_{e\text{-}max}$*** = 5.26*$10^{-15}$ mol $e^-$ $cell^{-1}$ $h^{-1}$. The estimated initial number of growing cells (***$N_G(0)$***) was 2.7*$10^3$ cells $mL^{-1}$, and the number after 110 hours (***$N_G(110)$***) was 1.6*$10^8$ cells $mL^{-1}$ (numbers are given in Figure 2C in the main paper). After 110 h, the $N_2O$-reduction rate ceased to increase exponentially, presumably because the provision of electron donors was insufficient to sustain a growth rate of 0.1 $h^{-1}$ for cell densities >1.6*$10^8$ cells $mL^{-1}$. Further growth was thus estimated by ***$N_G(t)$***= ***$N_G(110)$*** + **Δ$N_2$(t)*Y**, where ***$N_G(110)$*** is the cell density reached at t=110 h (1.6*$10^8$$mL^{-1}$), **Δ$N_2$(t)** is the cumulated of $N_2$ produced from 110 h and onwards (mol N $mL^{-1}$), and **Y** is the growth yield = 1.9*$10^{13}$ cells $mol^{-1}$ N. The cell specific rate of electron flow (***$V_{e-}$***, mol $e^-$ $cell^{-1}$ $h^{-1}$) throughout the enrichment culturing (Figure 2C of the main paper) was calculated by ***$V_{e-}(t)$***= ***$V_{N2}(t)$/$N_G(t)$*** where **$V_{N2}$(t)** is the measured rate of $N_2$ production (mol N $mL^{-1}$ $h^{-1}$) and ***$N_G(t)$*** is the estimated cell density, both at time t.

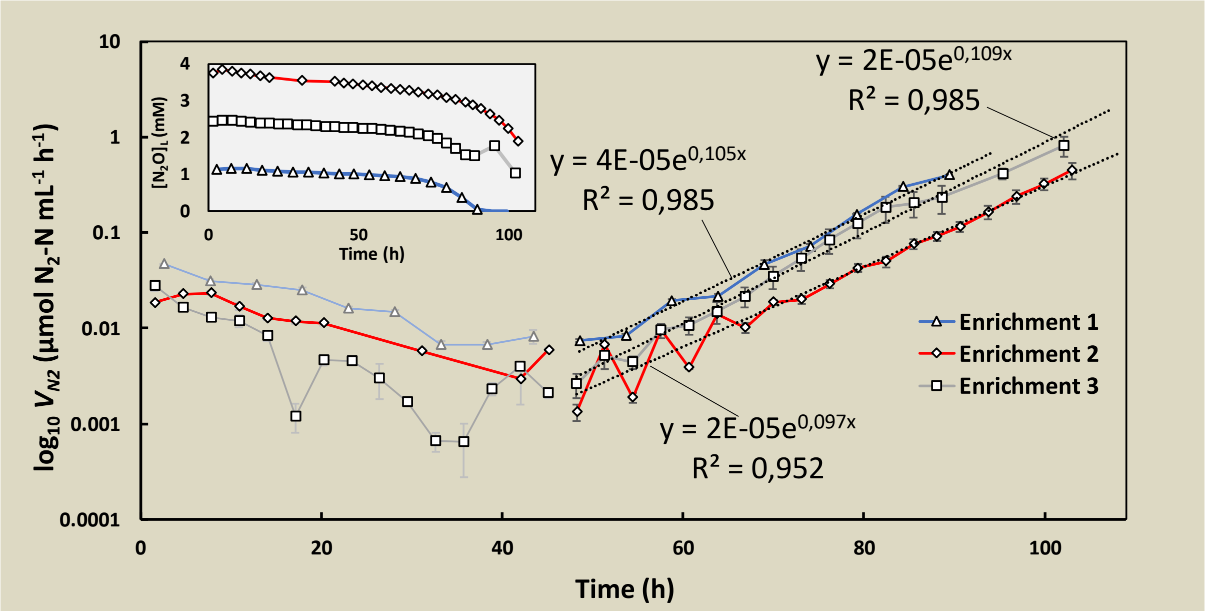


**Figure S2: Reproducible $N_2$ production rates during anaerobic incubation of a mesophilic digestate with $N_2O$.** Several enrichment experiments were run with mesophilic digestates, showing essentially identical $N_2$ production kinetics, i.e. declining rates during the first 50 hours, followed by exponential increase during the next 50 h. Enrichment 1 is the experiment used for metagenomics and metaproteomics (Figure 2A, main paper). Enrichment 2 is the experiment shown in Figure 2BC (main paper). All three enrichment experiments were equal except for the different initial concentrations of $N_2O$ which were 4, 15 and 9 vol% $N_2O$ in headspace for experiment 1, 2 and 3, respectively. The equilibrium concentrations in the digestate, given the temperature = 20 $^0$C, are 1.1, 4.2 and 2.5 mM $N_2O$ (Experiment 1, 2 and 3 respectively). In all enrichment experiments there were 3 replicates (vials), and the plotted values are average, with standard deviation as vertical lines. Different initial $N_2O$ concentrations were used (ranging from 1-4 mM in the digestate; see inserted panel), without any significant effect on the $N_2$ kinetics. The insert shows the concentrations of $N_2O$ in the digestate during the first 100 hours of each enrichment experiment.

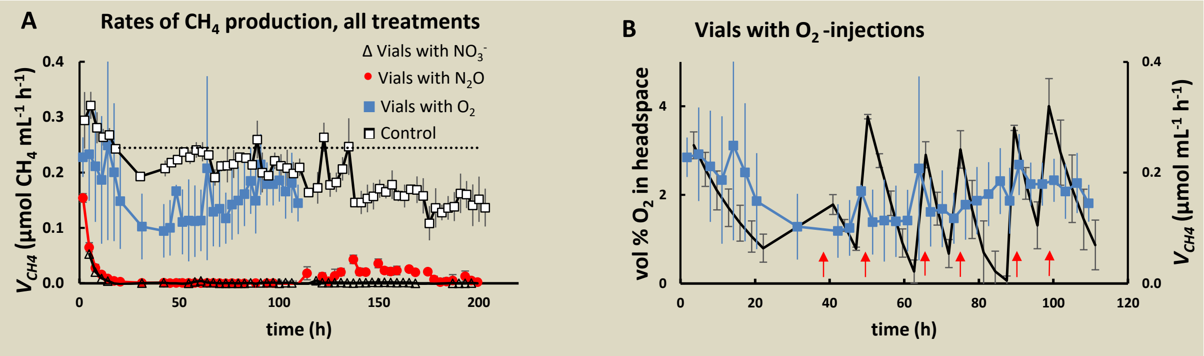


**Figure S3: Effects of $O_2$, $NO_3^-$ and $N_2O$ on methane production.** Freshly sampled digestate from the mesophilic anaerobic digester (37 °C) was incubated (20 °C) as stirred batches (50 mL in 120 mL vials, three replicates for each treatment), provided with either $O_2$, $N_2O$, $NO_3^-$, or without any electron acceptors added (=control). Oxygen and $N_2O$ concentrations were sustained by repeated injections, while $NO_3^-$ was supplied by peristaltic pumping of $KNO_3$ via a needle through the septum (1.57M $KNO_3$, flow rate 0.02 mL h$^{-1}$ => 31.4 µmol $NO_3^-$ h$^{-1}$). The vials were monitored for gas concentrations ($N_2$, NO, $N_2O$, $O_2$, $CH_4$, $CO_2$) in the headspace. Panel A shows the methane production rate ($V_{CH4}$) in all treatments (standard deviation shown as vertical lines). While $CH_4$ production was effectively suppressed by $NO_3^-$ and $N_2O$, $V_{CH4}$ in the vials with $O_2$ was 50-75% of that in the control vials (i.e. vials without any electron acceptors, basically resembling the anaerobic digester). The methane production in the mesophilic anaerobic digester was 1.48 µmol $CH_4$ mL$^{-1}$ h$^{-1}$ (Materials and Methods), which is an order of magnitude higher than the measured production rate in the control treatment. The temperature difference between the digester and the vials could account for this difference; indeed based on the apparent activation energy (Ea) for methane production in anaerobic digesters, determined by Elsgaard et al (2016) to be 80 kJ mol$^{-1}$, one would predict that the rate declines from 1.48 to 0.24 µmol $CH_4$ mL$^{-1}$ h$^{-1}$ by the downshift from 37°C (in the digester) to 20 °C (in the vials). This temperature-extrapolated rate of methane production is shown as a dashed line in Panel A. The similarity between the predicted and observed methane production rates in the control vials shows that a competent methanogenic consortium was maintained during culturing at 20 °C. Panel B shows the methane production together with the $O_2$ concentration in the headspace (vol % $O_2$) for the vials with oxygen in the headspace (standard deviation as vertical lines). The oxygen concentrations in the headspace fluctuated between 0 and 4 vol % ($O_2$ injection events marked by red arrows). The low rate of stirring in these experiments (300 rpm), implies relatively slow transport of $O_2$ from headspace to the liquid, and probably uneven distribution of $O_2$ within the liquid volume. Hence, there may have been anaerobic zones and microsites within the liquid, which could explain the sustained methanogenesis. The coexistence of aerobic and anaerobic metabolism, including methanogenesis is bioreactors has often been observed, and the inhibitory effect of low concentrations of oxygen on the methanogenesis in bioreactors may be marginal (Botheju and Bakke 2011).

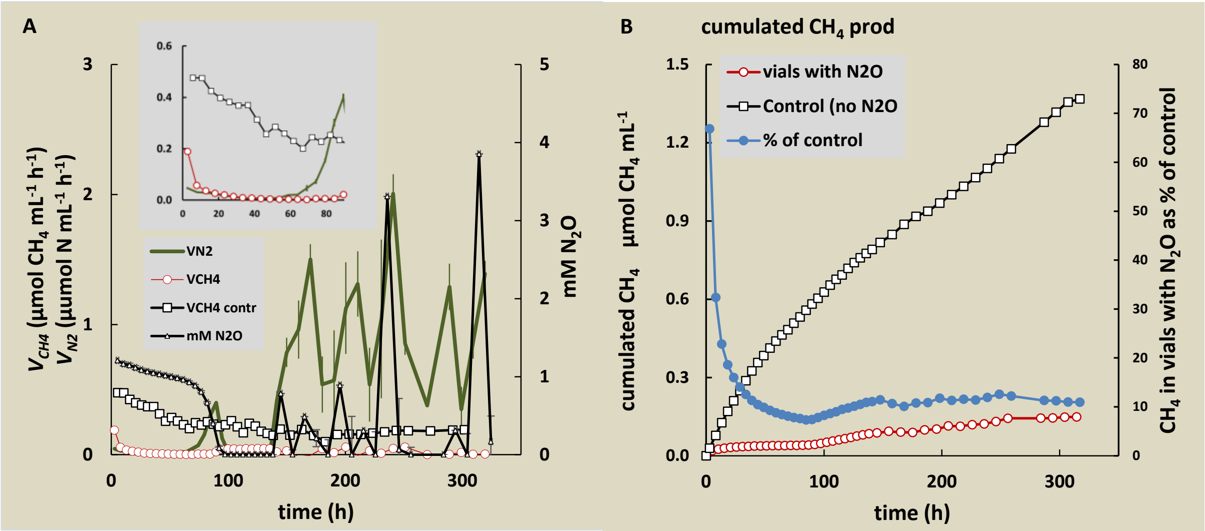


**Figure S4: Methane production in the enrichment culture used for metagenomics and metaproteomics.** In parallel with the enrichment culturing of $N_2O$ reducers which was used for -omics analyses **(Figure 2A in the main paper),** we monitored control vials (n=3), i.e. vials without $N_2O$ in headspace. Panel A shows the $N_2O$ concentration in the digestate (mM $N_2O$), the rates of $N_2$ production (***$V_{N2}$***) and methane production (***$V_{CH4}$***) in the vials with $N_2O$ in the headspace, as well as the rate of methane production in vials without $N_2O$ (***$V_{CH4\,control}$***). Panel B shows cumulated $CH_4$ production in the control vials without $N_2O$ (µmol $CH_4$ vial$^{-1}$) and in the vials with $N_2O$, as well as the latter expressed as % of $CH_4$ accumulation in the control. The data in panel B show that inhibition of methanogenesis by $N_2O$ was incomplete, the total methane production in the $N_2O$ enrichment vials being ~10 % of that in the control vials. The insert in panel A shows $V_{CH4}$, $V_{CH4control}$ and $V_{N2}$ (symbols the same as in the main panel) for the first 90 hours. Hypothetically, the apparent $N_2O$ inhibition of methanogenesis could be caused by $N_2O$-driven methanotrophy ($N_2O$ replacing $O_2$ as co-substrate for methane monooxygenase), but if so, the oxidation of 1 mole $CH_4$ would reduce 2 mol of $N_2O$ to $N_2$, i.e. ***$V_{N2}$*** $=2*($***$V_{CH4control}$*** $-$**$V_{CH4}$**$)$. The inserted panel shows that the measured rates of $N_2O$-reduction was clearly insufficient, and the hypothesis must be rejected.

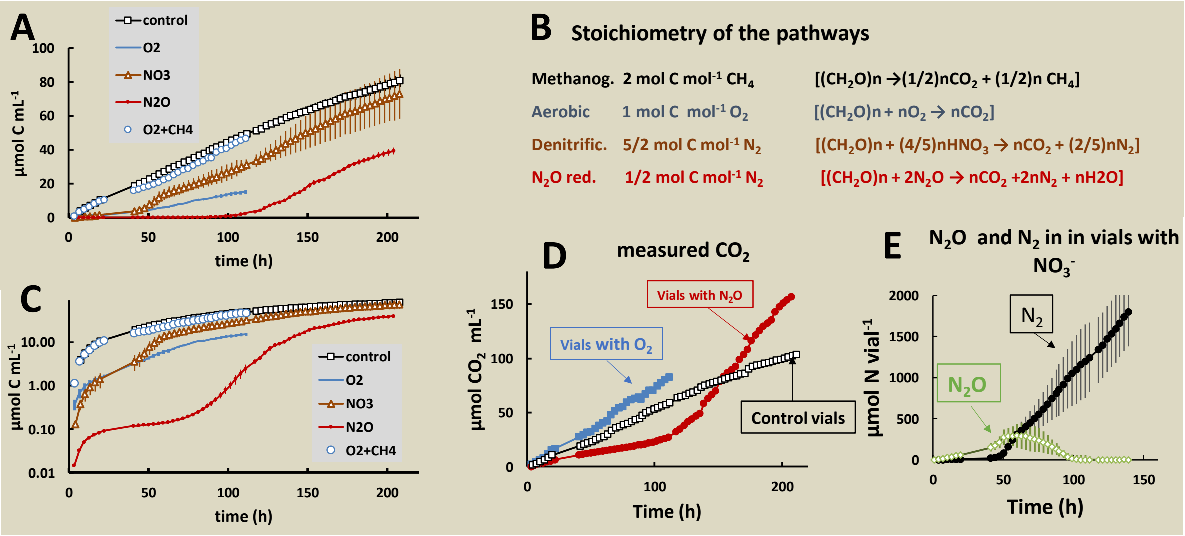


**Figure S5: Estimated carbon mineralization with different terminal electron acceptors: $O_2$, $N_2O$, $NO_3^-$ and $CO_2$.** The Figure shows the effect of $O_2$, $N_2O$ and $NO_3^-$ on apparent C-mineralization rates for the experiment presented in Figure S3. Panels A and C show the estimated C-mineralization for the different pathways (linear and logarithmic scale, in A and C, respectively), based on measured gas consumption/production, and the stoichiometry of the pathways (Panel B). For the vials with $NO_3^-$, the stoichiometry was corrected for the transient accumulation of $N_2O$ (Panel E), since $NO_3^- \rightarrow 1/2\ N_2$ consumes 5 electrons mol$^{-1}$ N, while $NO_3^- \rightarrow 1/2\ N_2O$ consumes only 4. While $N_2O$ and $NO_3^-$ effectively inhibited methane production, this was not the case for $O_2$ (See Figure S3). For this treatment, two curves are shown (Panel A & C): one for the aerobic pathway alone (blue, marked $O_2$ in the legend), and one for the sum of aerobic respiration and methanogenesis (marked $O_2$+$CH_4$ in legend). This shows that aerobic respiration accounts for approximately 50% of the C mineralization.

Panel D: Estimated $CO_2$ production based on measure $CO_2$ in headspace. The values are uncertain, because the digestate contained large amounts of $CO_2$ and $HCO_3^-$ when sampled (high partial pressure of $CO_2$ in the digester), and the pH in the digestate was high (7.6), which means that minor changes in pH throughout the incubation would affect the proportion of $CO_2$ present as $HCO_3^-$ in the liquid. Nevertheless, the estimated of $CO_2$ production showed similar contrasts between treatments as the estimates based on stoichiometry (Panel A): both show a retarded mineralization in the $N_2O$ treatment during the first 100 h compared to the control and the oxic treatment. No estimates could be made for the treatment with $NO_3^-$ because denitrification raised the pH (measured only at the end), causing declining $CO_2$ concentrations in the headspace.

Panel E: Transient accumulation of $N_2O$ during the incubation with $NO_3^-$. During the first 50 h, $NO_3^-$ was reduced to $N_2O$ exclusively, reflecting that in the original digestate, bacteria that reduce $NO_3^-$ to $N_2O$ outnumber those that are able to reduce $N_2O$ to $N_2$. This is corroborated by the estimated C kinetics shown in Panel A-D (early onset of mineralization based on $NO_3^-$ reduction while that based on $N_2O$ reduction was initially very slow).

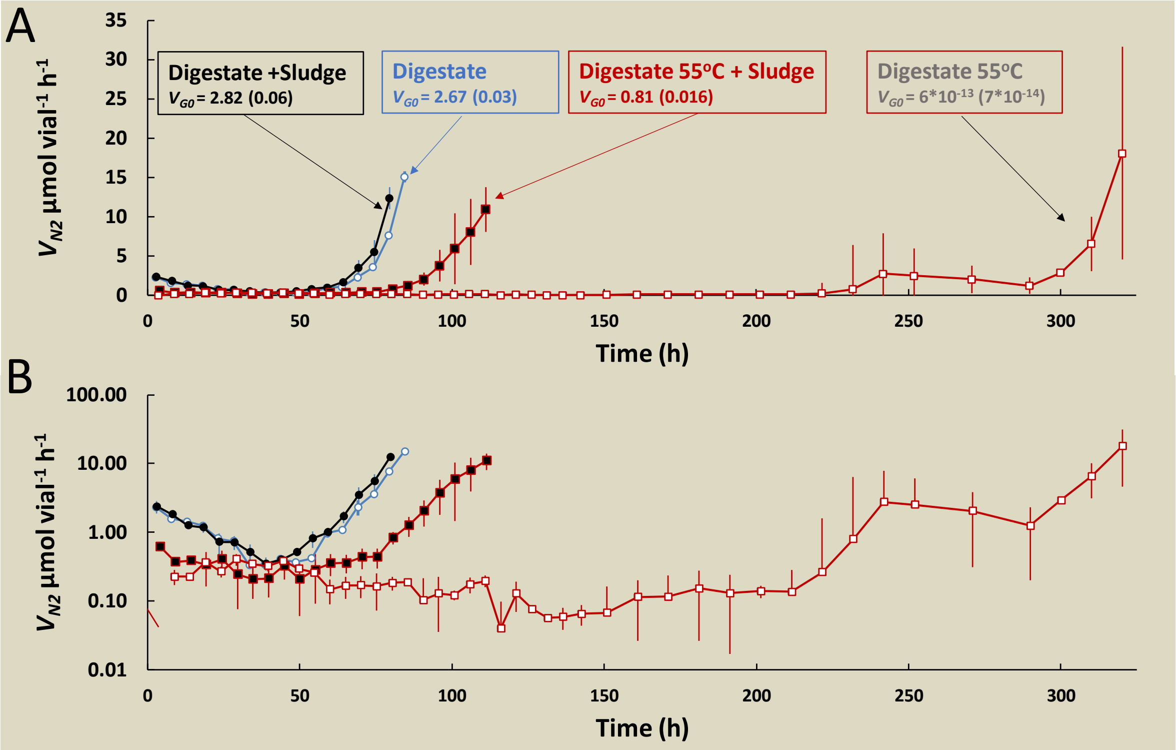


**Figure S6: Comparison of $N_2O$ reducing bacteria in raw sludge and the digestate.** We hypothesized that only a fraction of the $N_2O$-reducing bacteria in the sludge would survive the passage through the anaerobic digester, and checked this by investigating the $N_2O$ reduction kinetics in enrichment cultures with digestate, and digestate which had been heated to 55 $^0$C, with and without the addition of raw sludge (50 mL digestate +/- 1 mL raw sludge in each vial). The panel shows the measured rate of $N_2$ production (with standard deviation, n=3) for the four treatments. As in Figure S1, we estimated the initial rate of $N_2O$ reduction as a proxy for the density of $N_2O$-reducing bacteria, using the same model:

$$V_{N2} = V_D + V_G = V_{D0} * e^{-dt} + V_{G0} * e^{\mu t}$$

The model was fitted to the data for each single vial, using the Generalized Reduced Gradient Solver in Excel, resulting in three independent estimates of $V_{G0}$ for each treatment (one for each replicate vial). Panel A shows measured $V_{N2}$ for all treatments plotted against time (average values, standard deviation as vertical lines, n=3). The boxes show the average estimated initial rates $V_{G0}$ as µmol N vial$^{-1}$ h$^{-1}$, with standard deviations in parenthesis (n=3). Panel B shows the same data with a log scaled Y-axis.

$V_{G0}$ in the 1 mL sludge added can be estimated by the increase in $V_{G0}$ by adding 1 mL sludge to the digestates:

Unheated digestate: $V_{G0\_sludge}$ = $V_{G0\_digestate+sludge}$ – $V_{G0-digestate}$ = 0.15 µmol N mL$^{-1}$ sludge h$^{-1}$

Heated digestate: $V_{G0\_sludge}$ = $V_{G0\_digestate55+sludge}$ – $V_{G0-digestate55}$ = 0.81 µmol N mL$^{-1}$ sludge h$^{-1}$

The two $V_{G0\_sludge}$ estimates are very different, but they are both much higher than $V_{G0-digestate}$, which was 2.67 µmol N vial$^{-1}$ h$^{-1}$ (panel A) = 0.05 µmol N mL$^{-1}$ digestate h$^{-1}$. The fraction of $N_2O$ reducers which survives the passage of the anaerobic digester is estimated by **F=** $V_{G0-digestate}/V_{G0\_sludge}$, and F= 0.33 and 0.06 (based on the two widely different estimates of $V_{G0\_sludge}$. A reasonable conclusion is that ≤1/3 of the viable $N_2O$-reducing organisms in the sludge survived the anaerobic digestion.

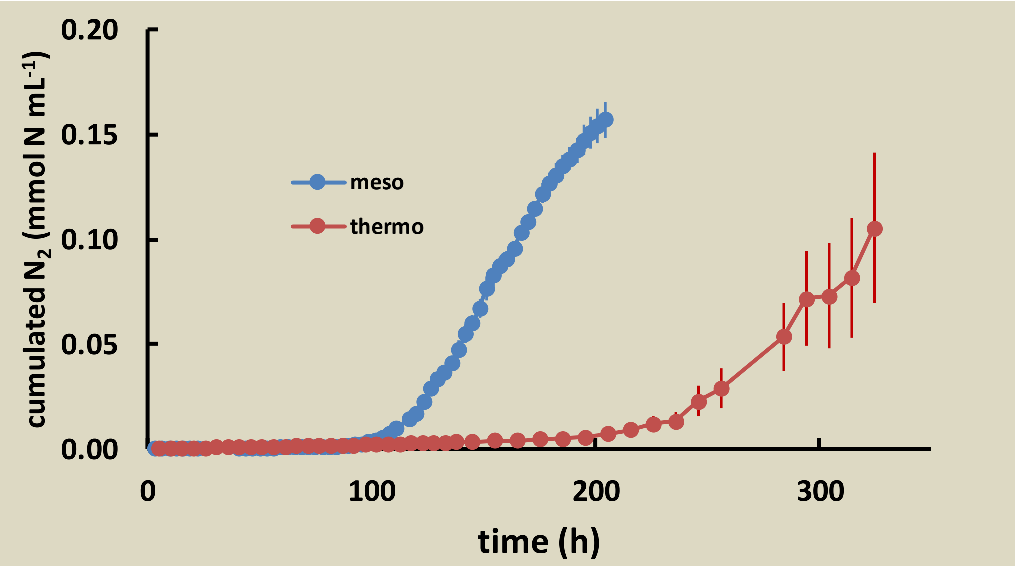


**Figure S7: Comparison of digestates from a mesophilic and a thermophilic digester.** We sampled digestates from the mesophilic and thermophilic digesters (37 °C and 52 °C respectively, both fed with the same sewage sludge), and incubated them anaerobically with $N_2O$ at 20 ºC as in previously presented experiments (**Figure 2** main paper, **Figures S1-5**). Measurements of $N_2$ production (this Figure) showed that the thermophilic digestate contained orders of magnitude lower number of $N_2O$-respiring organisms than the mesophilic digestate.

### B. Growth and decline of members of the microbial consortium

This section provides data regarding the growth or decline of different members of the microbial consortium of the digestate based on the abundance of individual MAGs in the metagenomes (**Supplementary Data S2)** and metaproteomes (**Supplementary Data S1)** at three timepoints during the enrichment (0, 115 and 325 hours; see Figure 2 and 3 in main paper).

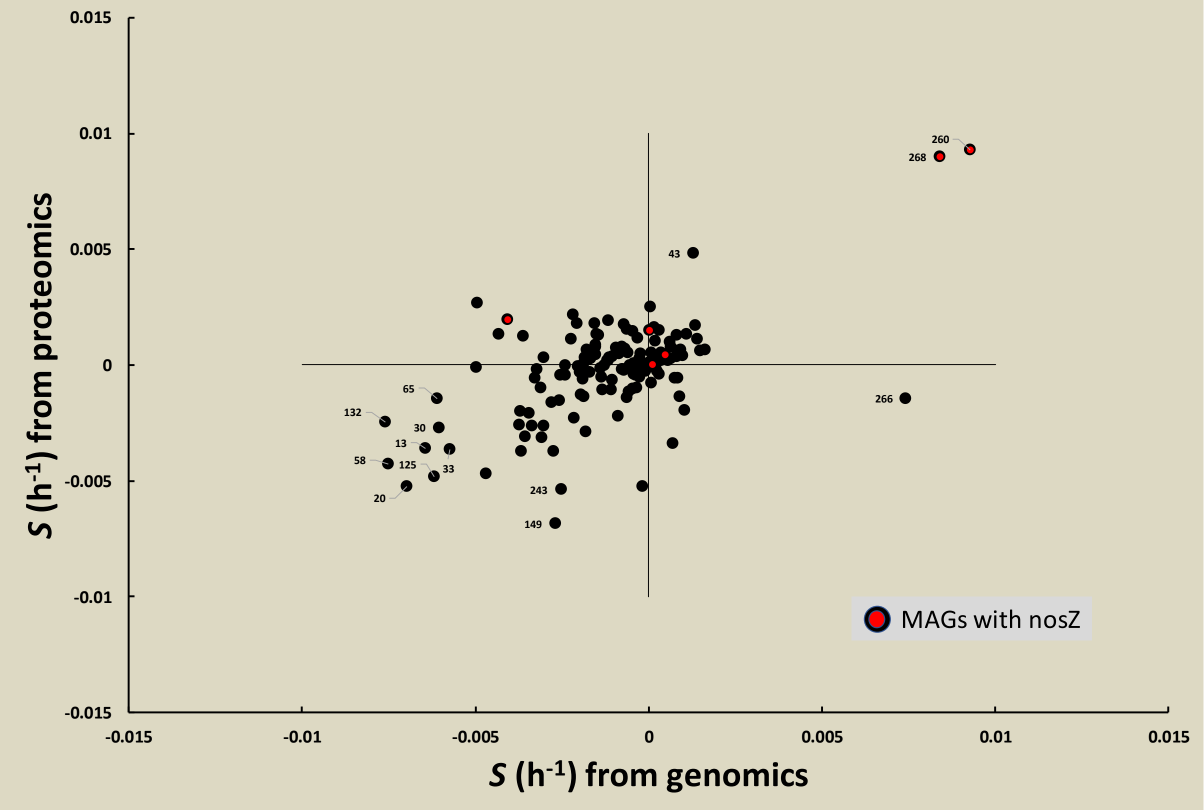


**Figure S8: Abundance of MAGs during enrichment culturing.** To assess how enrichment culturing with $N_2O$ affected the abundance of members of the microbial consortium, we used the metagenomic and metagenomic data to calculate the relative increase of each MAG: $S = \frac{q}{\bar{x}}$ , where $q$ is the regression coefficient for $x$ (x=coverage for genomics, sum of LFQ for proteomics) against time and $\bar{x}$ is the average for the MAG (all three time points), thus the unit for S is $h^{-1}$. The plot shows *S* from proteomics against *S* based on genomics for each MAG. MAGs with *nosZ* are marked with red circles. For the majority of MAGs, S ranged from -0.005 to +0.005 $h^{-1}$, regardless of the assessment method. The identity of the MAGs is shown for MAGS with at least one S-value outside this range. Only two of the MAGs with *nosZ* had S > 0.005 $h^{-1}$ (MAG260 & MAG268). For the organisms without *nosZ*, the average S was -0.0013 $h^{-1}$ for metagenomics data and -0.00045 $h^{-1}$ for metaproteomics data.

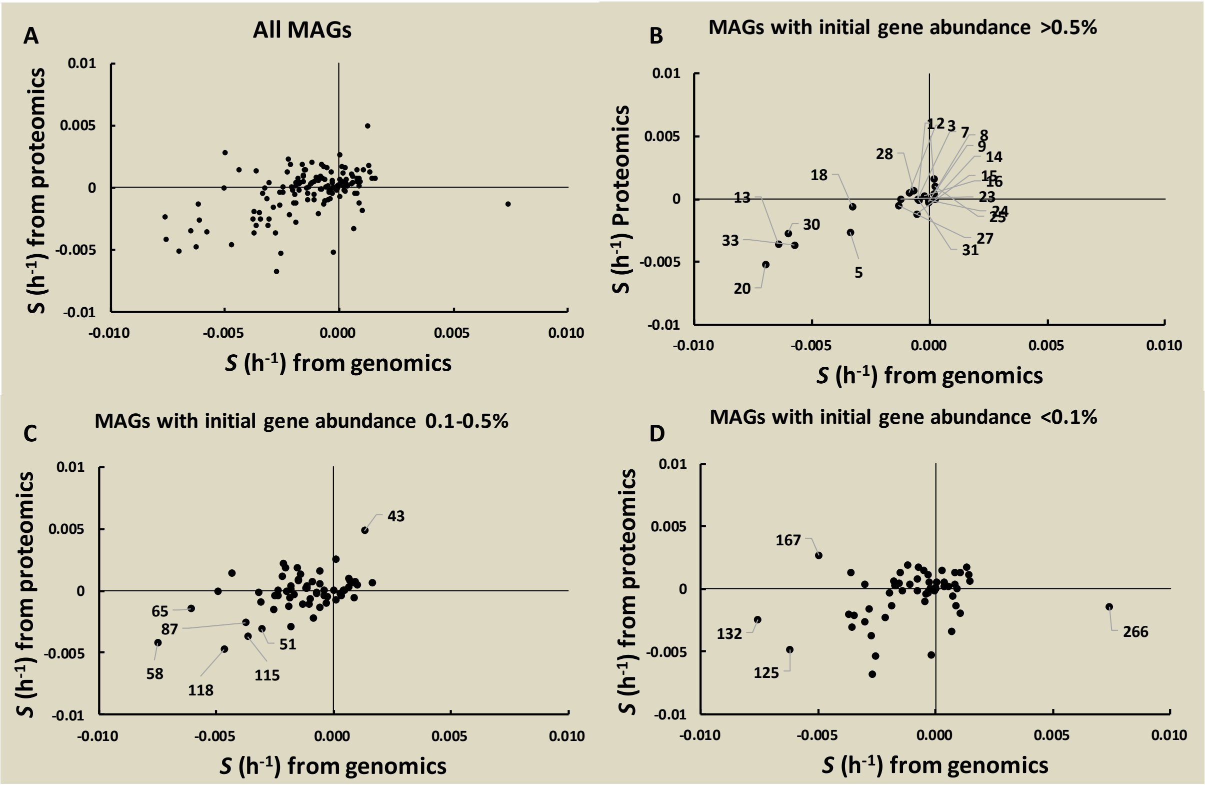


**Figure S9: Evaluation of growth/decline of MAGs without *nosZ*, stratified according to relative abundance.** To inspect if the growth/decline differed depending on the initial population size, we stratified the MAGs into three groups, i.e. MAGs with initial genomic abundance > 0.5 %, 0.1-0.5 % and < 0.1 %, and plotted the relative increase, *S* (as calculated for Figure S8) as calculated from proteomics against that from genomics. Panel A shows the plot of for all MAGS, and Panels C-D shows the plots for the three strata. The stratification demonstrated no clear relationship between initial abundance and the apparent ability to survive during the enrichment culturing: within each group, the majority of MAGs clustered around zero, while a minority showed a declining trend, both for the genomics and the proteomics.

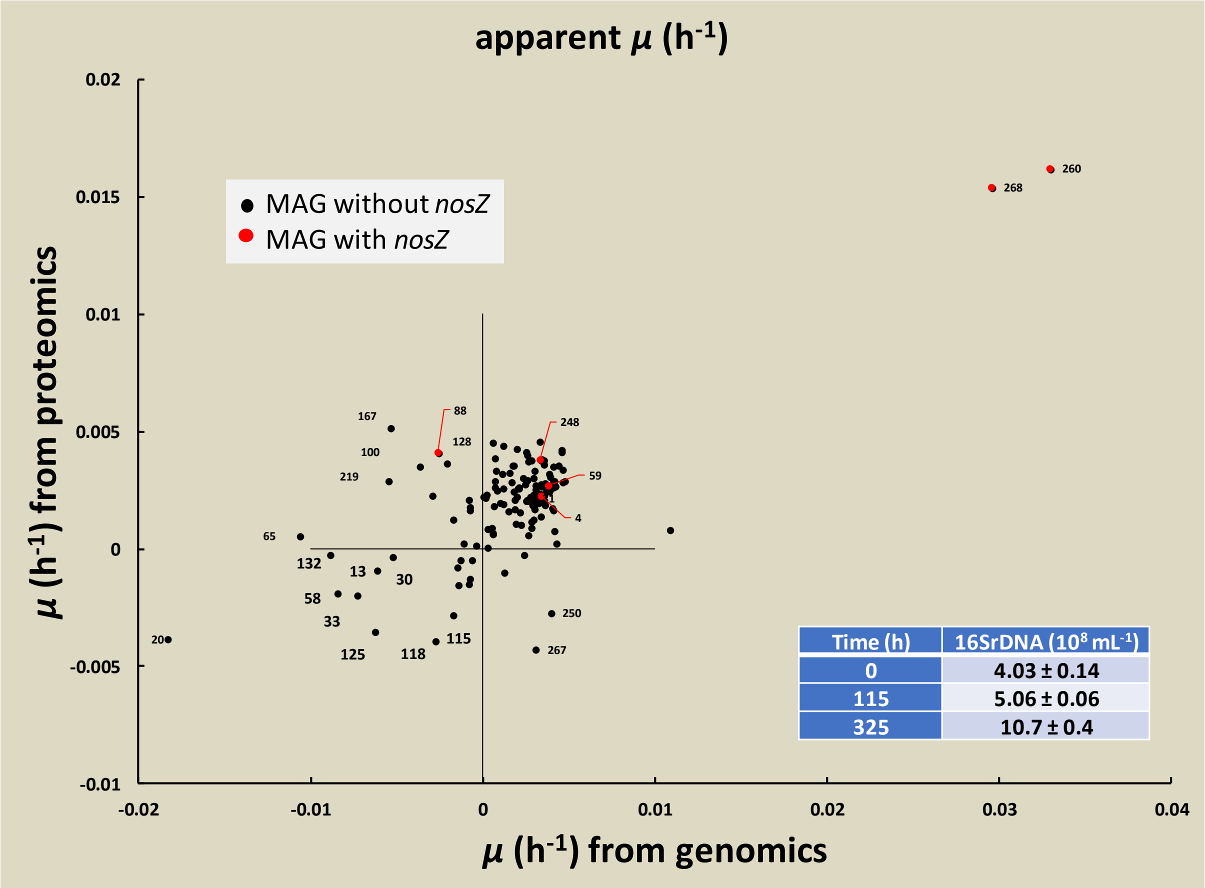


**Figure S10: Apparent rate of growth/decline of MAGs by combining -omics and measured abundance of 16SrDNA.** As a final approach to evaluate growth and decline of specific populations during the enrichment culturing, we combined omics data with 16SrDNA abundance (16S copies mL$^{-1}$ digestate measured by digital droplet PCR, using universal primers), to assess the abundance of individual MAGs and to calculate the apparent growth rates (or decline) during enrichment culturing. Average total 16S rDNA abundance at each time point is shown in the inserted in panel, with standard error (n=3). For each time point (t= 0, 115 and 325 h), the cell density of each MAG was assessed by $N_{it}=S_t*C_{it}/\sum C_t$ where $S_t$ is the measured 16SrDNA abundance at time ***t***, and $C_{it}/\sum C_t$ is the MAG's relative abundance at time t. For genomics, $C_{it}$ = the MAG's coverage at time t, $\sum C_t$ = the total read coverage of all 149 MAGS at time. For proteomics, $C_{it}$ = the pooled LFQ value for the MAG at time t, $\sum C_t$ = the sum of pooled LFQ for all MAGS at time t. The apparent growth/death rate was estimated by the slope of **ln($N_{it}$)** against time (linear regression). The results indicate slight growth (0-0.005 h$^{-1}$) for the majority of organisms (upper right quadrant), consistent strong growth for *nosZ* encoding MAG260 and MAG268, and consistent decline for 9 MAGs (lower left quadrant). The genomics- and proteomics-based μ were inconsistent for 8 MAGs (upper left and lower right quadrant). The relative abundance of the 9 declining MAGs is listed in **Table S1**). We did not find any convincing common traits between the MAGs that could explain their decline. One possible reason for their declining abundance could be inability to adapt to the lower temperature (20°C in the enrichment culture versus 37°C in the digester).

**Table S1: Relative abundance from % total read coverage and % LFQ for MAGs that declined during the enrichment.** GTDB classifications were assigned at a phylum level. MAG relative abundance was calculated as % of total read coverage of the 149 MAGs and from relative LFQ%, which denotes the relative protein intensity calculated as % of LFQ assigned to an individual MAG relative to the summed LFQ for the 149 MAGs used to construct the metaproteome database.

| **MAG ID** | **GTDB classification (phylum)** | **Relative abundance (%)** | | | **Relative LFQ (%)** | | |
|---|---|---|---|---|---|---|---|
| | | **Time (h)** | | | **Time (h)** | | |
| | | **0** | **115** | **325** | **0** | **115** | **325** |
| MAG20 | Myxococcota | 0.764 | 0.176 | 0.001 | 4.076 | 1.017 | 0.388 |
| MAG13 | Spirochaetota | 1.105 | 0.248 | 0.052 | 0.915 | 0.213 | 0.209 |
| MAG132 | Spirochaetota | 0.082 | 0.004 | 0.001 | 0.050 | 0.026 | 0.016 |
| MAG30 | Spirochaetota | 0.511 | 0.144 | 0.033 | 2.895 | 1.155 | 0.872 |
| MAG58 | Spirochaetota | 0.269 | 0.015 | 0.005 | 1.933 | 0.459 | 0.330 |
| MAG33 | Desulfobacterota | 0.531 | 0.264 | 0.020 | 0.339 | 0.176 | 0.064 |
| MAG125 | Firmicutes | 0.104 | 0.031 | 0.005 | 0.510 | 0.166 | 0.056 |
| MAG115 | Firmicutes | 0.108 | 0.097 | 0.025 | 0.135 | 0.096 | 0.021 |
| MAG118 | Thermotogota | 0.105 | 0.055 | 0.016 | 1.386 | 0.601 | 0.141 |

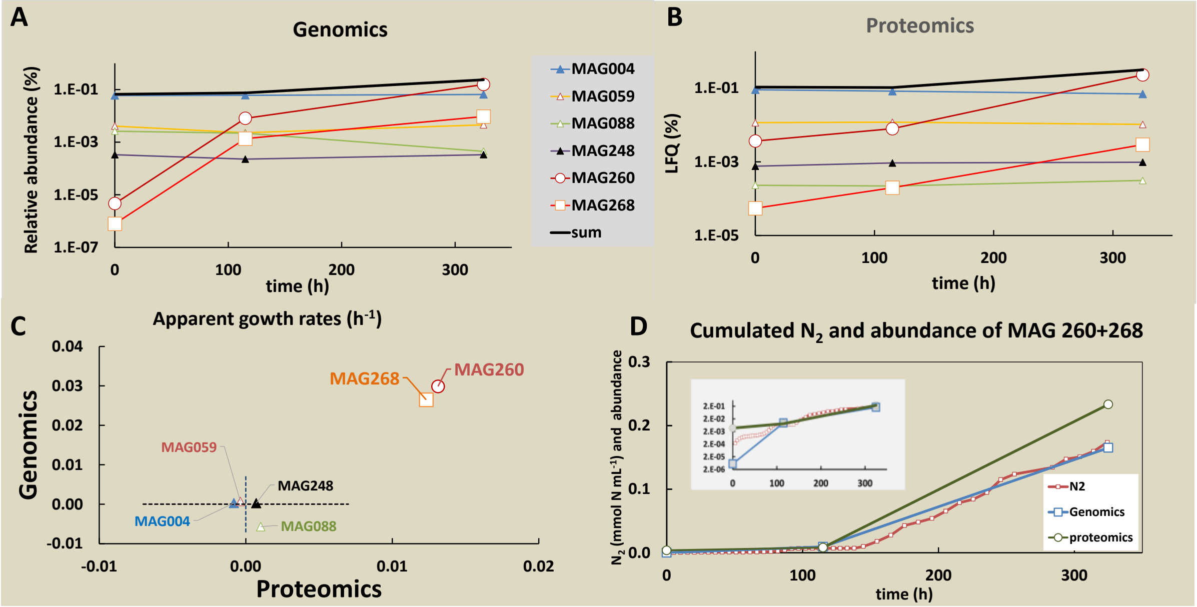


**Figure S11: Abundance of MAGs with *nosZ***. Panel A shows the relative abundance of the *nosZ* containing MAGs based on genomics (reads as fraction of the sum of 149 MAGs). Panel B shows the relative abundance based on proteomics (LFQ as fraction of the sum of 149 MAGs). Panel C shows a crude estimate of apparent growth rates based on genomics and proteomics (= slope of ln(N) against time; N=relative abundance). These results demonstrate substantial growth for MAG260 and MAG268, but not for the other MAGs. In panel D, the sum of MAG260 and 268 is plotted against time, together with cumulated $N_2$ (derived from data shown in Figure 2, main paper). The inserted panel shows the same data on a log scale. This shows that the sum of the abundance of the two MAGs (based on proteomics of genomics) increased as cumulated $N_2O$-reduction to $N_2$ increased.

### C. Metabolism of the methanogenic consortium

Here we present a metaproteome-centric metabolic map of possible substrate flows in the microbial consortium, and experimental evidence for the predicted effects of $N_2O$-inhibition of methanogenesis in this consortium: accumulation of volatile fatty acids and hydrogen.

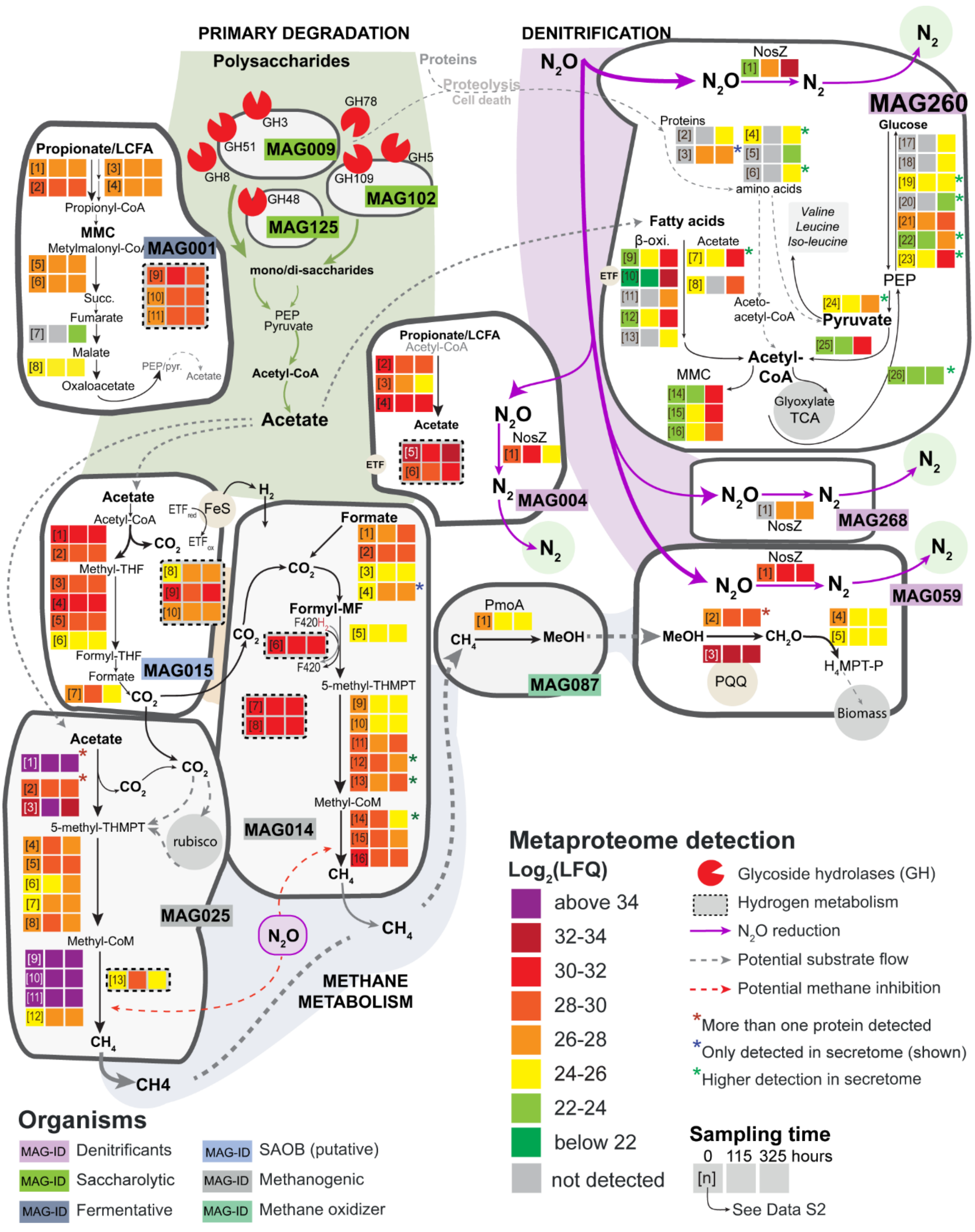


**Figure S12: Metaproteome-centric metabolic map of the substrate flow in the microbial consortium.** For metabolic reconstruction of the substrate flow, including primary degradation of carbon sources and $N_2O$ reduction, we scanned the detected proteins affiliated to each MAG for enzymes involved in specific metabolic pathways. Detected protein levels ($log_2$(LFQ)) for the three sampling timepoints (after 0, 115 and 325 hours) are indicated by colored squares, where the number in the first square corresponds to the number in the first column in **Supplementary Data S2**. A variety of carbohydrate-active enzymes (CAZymes), including those active on cellulose (members of GH5, GH8 and GH48) were detected in the metaproteome. Only a selection is shown in the figure; a complete list of detected proteins annotated as CAZymes can also be found in **Supplementary Data S2**. Multiple MAGs also expressed proteins for fermentation processes and production of acetate and propionate. The metaproteome further supported the assumption that these fermentation products are metabolized by a population which included $N_2O$-reducers, represented by MAG004 and MAG260 in the figure. $N_2O$ has been suggested to inhibit the enzymatic process of methanogenesis (Andalib et al 2011, Kengen et al 1988), which was supported by reduced methane-production rate in the microbial enrichment incubated with $N_2O$ in the current study. Yet, the protein detection level of key enzymes involved in both hydrogenotrophic and acetoclastic methanogenesis (MAG014 and MAG025, respectively) were amongst the highest detected in the metaproteome, even after 115 and 325 hours of incubation. Another MAG with numerous highly detected proteins was affiliated to *Dethiobacteria*, a class recently suggested to encompass syntrophic acetate oxidizing bacteria (SAOBs) (Mosbæk et al 2016, Dyksma et al 2020). This MAG, MAG015, expressed proteins related to the Wood-Ljungdahl pathway (WLP), and combined with the detection of an electron transfer complex (gene cluster encompassing iron-sulfur ferredoxin, coenzyme $F_{420}$ hydrogenase/dehydrogenase, electron transfer flavoprotein) and an aldehyde ferredoxin oxidoreductase for potential acetate activation (Swanson et al 2008, Keller et al 2019) we postulate that also this representative of *Dethiobacteria* might use WLP in a reverse direction to oxidize acetate. This was strengthened by the detection of enzymes central for β-oxidation of longer-chained fatty acids and the detection of the fructose diphosphatase used in anabolic metabolism (i.e., gluconeogenesis). Importantly, SAOBs are depending on an active hydrogen-scavenger population, often hydrogenotrophic methanogens (such as MAG014), to realize the oxidative direction of WLP, which reinforces our observations that the consortium is synergistically producing methane at some capacity. Finally, we detected predicted methane monooxygenase and methanol dehydrogenase proteins from MAG087 and MAG059 (respectively), which leaves tantalizing hypotheses as to the potential role of the methanotrophic community within this enrichment. Methanotrophic processes have recently been shown to be facilitated by the presence of $N_2O$ as a terminal electron acceptor (Valenzuela et al 2020; Cheng et al 2019), but direct links, which may possibly exist within this enrichment, remain to be elucidated.

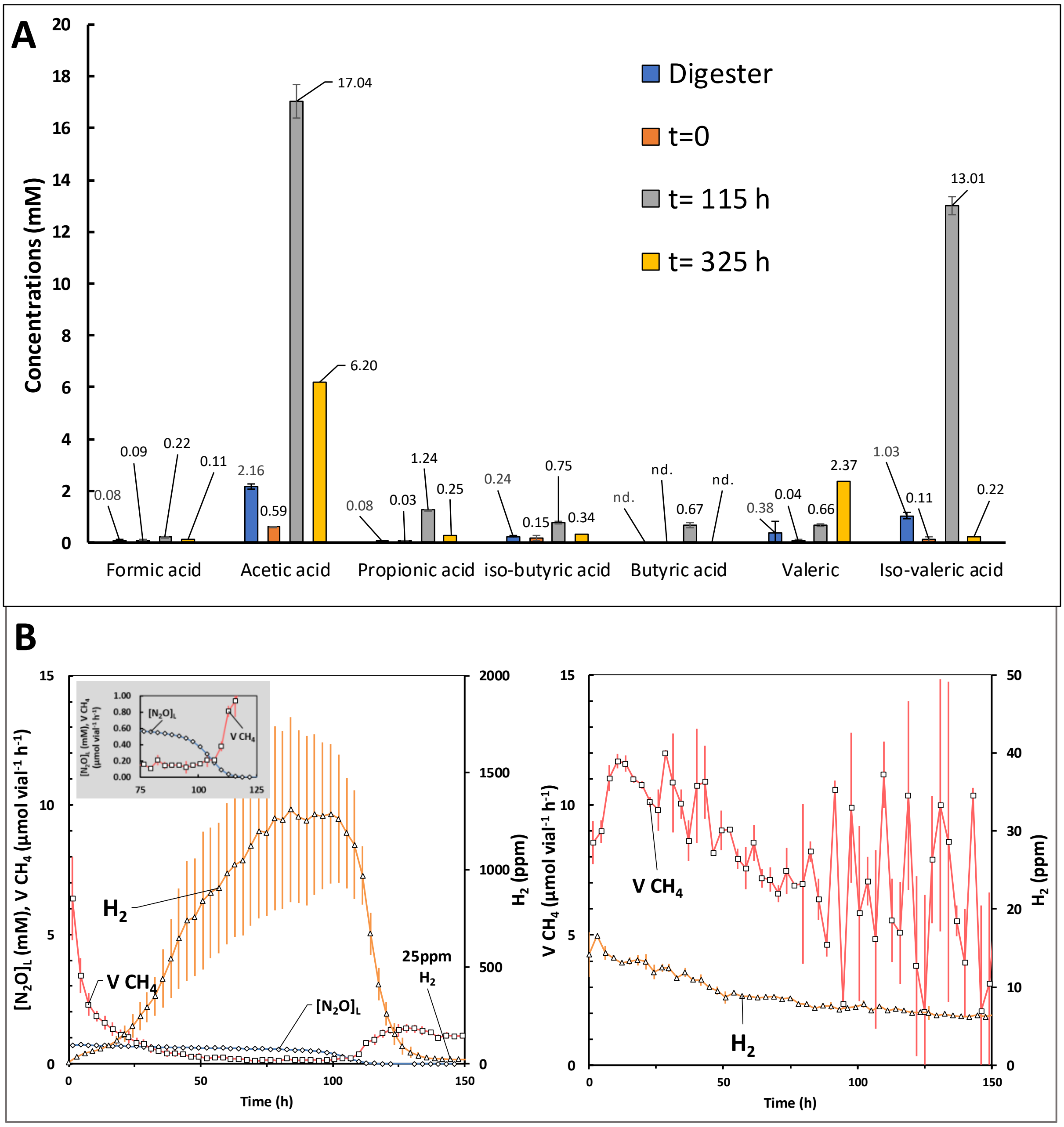


**Figure S13: Quantification of volatile fatty acids (VFA) and $H_2$ during enrichment culturing**. Inhibition of methanogenesis by $N_2O$ could result in transient accumulation of intermediates such as VFA (**Figure S12**), which might last until the $N_2O$-respiring bacteria have become sufficiently numerous to effectively reap these intermediates. $H_2$ might also accumulate, until the partial pressure of $H_2$ reaches levels high enough to sustain hydrogenotrophic acetogenesis (Wood-Ljungdahl pathway).

Panel A shows the VFA concentrations (mmol $L^{-1}$) in samples of digestate directly from the digester (Digester, n=3 replicates, frozen immediately after sampling from the anaerobic digester), and at the three time points (0, 115 and 325 h) of the enrichment culturing experiment number 1, presented in **Figure 2A** in the main paper (n=2 for t=115 h, and 3 for the others). All samples were stored at -80 °C before being prepared for VFA analysis. The lower concentration at the onset of the enrichment culturing (t=0) compared to that in the digester could be due to oxygenation during transport from the WWTP to the laboratory, and to losses due to the He-washing (evacuation and He-filling) prior to enrichment culturing. The results clearly show the expected transient accumulation of VFAs.

Panel B shows the accumulation of $H_2$ in response to $N_2O$-mediated inhibition of methane production. This was measured in a repetition of the enrichment culturing shown in **Figure 2** of the main paper, using an improved version of the incubation robot system which measures $H_2$ by a Plasma Emission detector (PED) (©LDetek). The left panel shows the results for vials with $N_2O$ in the headspace: concentration of $N_2O$ in the liquid, concentration of $H_2$ in the headspace, and the rate of $CH_4$ production ($V_{CH4}$). The insert is scaled to show the onset of $CH_4$ production in response to $N_2O$ depletion. The right panel shows $H_2$ and $V_{CH4}$ in vials without $N_2O$. These results corroborate the hypothesis that $H_2$ accumulates in response to $N_2O$-inhibition of methanogenesis, reaching an apparent steady state concentration around 350 ppm in the headspace ($P_{H2}$=3.5*10$^{-4}$ bar = 0.28 µM $H_2$ in the liquid).

### D. Genetics of isolated organisms

Here, we present the results of genome sequencing of the three isolates, their phylogeny, their core denitrification reductase genes as well as genes coding for peripheral proteins which contributes to the denitrification pathway.

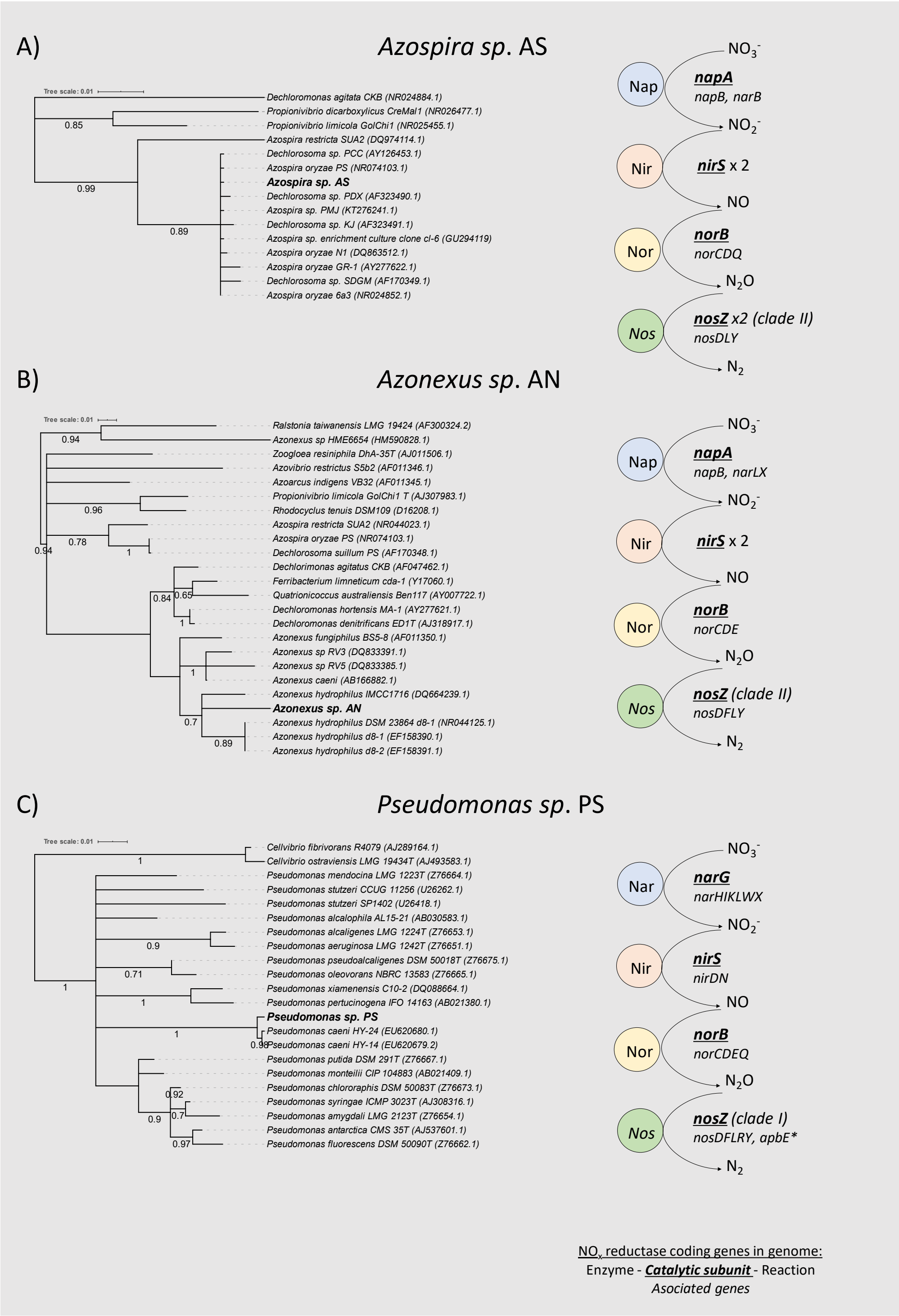


**Fig S14: Phylogeny and denitrification genes annotated in draft genomes of isolated organisms**. The panel shows the maximum likelihood phylogenetic trees of the three isolates, based on full length 16S rRNA DNA sequences (bootstrap values > 0.6 , 100 resamplings), their core denitrification genes coding for the four denitrification reductases (Nar/Nap, Nir, Nor and Nos), as well as a number of genes coding for peripheral proteins that contribute to a fully functional denitrification pathway (Vaccaro et al 2016). One of these is *nosR,* which was only found in *Pseudomonas sp*. PS. NosR is hypothesized to be involved in electron donation to Nos (Wunsch and Zumft 2005; Zhang et al 2017), but apparently only to Nos Clade I, because organisms with *nosZ clade II* often lack *nosR* (Hein et al 2017), which was the case for the two isolates with nosZ Clade II (AS and AN). *Pseudomonas sp.* PS lacked *nosX*, a flavin donor involved in maturation of norR, but insteadthe *apbE* (coding for flavin transferase, EC: 2.7.1.180) that has been suggested as a flavin donor candidate in maturation via covalent flavinylation of NosR in *Pseudomonas stutzeri* (Zhang et al 2017).

**Table S2**: QUAST quality parameters, PROKKA annotation summary, CheckM genome quality parameters and coverage of SPAdes assembled contigs of *Pseudomonas sp.* PS, *Azospira sp.* AS and *Azonexus sp.* AN.

| | *Pseudomonas sp.* PS | *Azospira sp.* AS | *Azonexus sp.* AN |
|---|---|---|---|
| | QUAST quality parameters: | | |
| Contigs total: | 21 | 75 | 59 |
| Largest contig (bp): | 778 581 | 409 855 | 279 444 |
| Contigs (>= 0 bp): | 21 | 75 | 59 |
| Contigs (>= 1000 bp) | 18 | 68 | 48 |
| Contigs (>= 10 000 bp) | 13 | 44 | 26 |
| Contigs (>= 100 000 bp) | 9 | 11 | 13 |
| Total length (bp): | 3 378 613 | 3 810 942 | 2 882 318 |
| N50: | 440 975 | 133 847 | 176 807 |
| L50: | 3 | 9 | 7 |
| Predicted genes (>= 300 bp): | 2349 | 3019 + 11 partially | 2364 + 6 partially. |
| GC (%): | 47.89 | 65.42 | 60.82 |
| Mismatches: | | | |
| # N's | 195 | 199 | 109 |
| # N's per 100 kbp | 5.77 | 5.22 | 3.78 |
| | SPAdes output | | |
| Coverage (k-mer): | 163.8x | 59.72x | 90.12x |
| | Prokka annotation summary: | | |
| Number of genes predicted: | 3153 | 3518 | 2766 |
| Number of protein coding genes: | 3104 | 3461 | 2713 |
| Number of genes with non-hypothetical function: | 2201 | 2404 | 1839 |
| Number of genes with EC-number: | 1223 | 1247 | 988 |
| Number of genes with Seed Subsystem Ontology: | 972 | 995 | 807 |
| Average protein length: | 323 | 327 | 319 |
| | CheckM genome quality parameters: | | |
| Marker Lineage | C_Gammaproteobacteria | C_Betaproteobacteria | C_Betaproteobacteria |
| #Genomes | 263 | 223 | 233 |
| #Markers | 507 | 424 | 425 |
| #Marker sets | 232 | 211 | 211 |
| 0 | 6 | 0 | 4 |
| 1 | 498 | 422 | 421 |
| 2 | 3 | 2 | 0 |
| 3 | 0 | 0 | 0 |
| 4 | 0 | 0 | 0 |
| 5+ | 0 | 0 | 0 |
| Completeness | 97.7 | 100.0 | 98.1 |
| Contamination | 0.89 | 0.26 | 0.00 |

### E. Carbon substrate utilization by isolated organisms

Here we show the results of the testing C substrate utilization by the three isolated organisms. The result show that PS (P*seudomonas sp.*) could utilize a wide specter of substrates, although it's capacity to utilize polymers was marginal. In contrast AN (*Azonexus sp.*) and AS (*Azospira sp.*) utilized very few C substrates, primarily intermediates of anaerobic fermentation of a methanogenic consortium.

**Table S3: Screening the three isolates for C utilization** using Biolog Phenotype MicroArray™ plates PM1 and PM2, which tests the isolates capacity to utilize various C substrates. Positive wells are marked with + (n=3 replicate plates)

| **PM1** | **PS** | **AS** | **AN** | **PM2** | PS | AS | AN |
|---|---|---|---|---|---|---|---|
| A2 L-Arabinose | | | | A2 Chondroitin Sulfate C | | | |
| A3 N-Acetyl-DGlucosamine | +++ | | | A3 α-Cyclodextrin | | | |
| A4 D-Saccharic Acid | | | | A4 ß-Cyclodextrin | | | |
| A5 Succinic Acid | +++ | +++ | +++ | A5 γ-Cyclodextrin | | | |
| A6 D-Galactose | | | | A6 Dextrin | | | |
| A7 L-Aspartic Acid | +++ | +++ | | A7 Gelatin | | | |
| A8 L-Proline | +++ | | | A8 Glycogen | | | |
| A9 D-Alanine | +++ | | | A9 Inulin | | | |
| A10 D-Trehalose | | | | A10 Laminarin | +++ | | |
| A11 D-Mannose | +++ | | | A11 Mannan | | | |
| A12 Dulcitol | | | | A12 Pectin | | | |
| B1 D-Serine | | | | B1 N-Acetyl-DGalactosamine | | | |
| B2 D-Sorbitol | | | | B2 N-AcetylNeuraminic Acid | | | |
| B3 Glycerol | +++ | | | B3 ß-D-Allose | | | |
| B4 L-Fucose | | | | B4 Amygdalin | | | |
| B5 D-Glucuronic Acid | | | | B5 D-Arabinose | | | |
| B6 D-Gluconic Acid | +++ | | | B6 D-Arabitol | | | |
| B7 D,L-α-GlycerolPhosphate | +++ | | | B7 L-Arabitol | +++ | | |
| B8 D-Xylose | | | | B8 Arbutin | | | |
| B9 L-Lactic Acid | +++ | +++ | | B9 2-Deoxy-DRibose | +++ | | |
| B10 Formic Acid | | | | B10 i-Erythritol | | | |
| B11 D-Mannitol | | | | B11 D-Fucose | | | |
| B12 L-Glutamic Acid | +++ | +++ | +++ | B12 3-0-ß-DGalactopyranosylD-Arabinose | | | |
| C1 D-Glucose-6-Phosphate | +++ | | | C1 Gentiobiose | | | |
| C2 D-Galactonic Acid-γ-Lactone | | | | C2 L-Glucose | | | |
| C3 D,L-Malic Acid | +++ | +++ | +++ | C3 Lactitol | | | |
| C4 D-Ribose | +++ | | | C4 D-Melezitose | | | |
| C5 Tween 20 | | | | C5 Maltitol | | | |
| C6 L-Rhamnose | | | | C6 α-Methyl-DGlucoside | | | |
| C7 D-Fructose | +++ | | | C7 ß-Methyl-DGalactoside | | | |
| C8 Acetic Acid | +++ | +++ | +++ | C8 3-Methyl Glucose | | | |
| C9 α-D-Glucose | +++ | | | C9 ß-Methyl-DGlucuronic Acid | | | |
| C10 Maltose | | | | C10 α-Methyl-DMannoside | | | |
| C11 D-Melibiose | | | | C11 ß-Methyl-DXyloside | | | |
| C12 Thymidine | +++ | | | C12 Palatinose | | | |
| D1 L-Asparagine | +++ | | | D1 D-Raffinose | | | |
| D2 D-Aspartic Acid | | | | D2 Salicin | | | |
| D3 D-Glucosaminic Acid | | | | D3 Sedoheptulosan | | | |
| D4 1,2-Propanediol | | | | D4 L-Sorbose | | | |
| D5 Tween 40 | | | | D5 Stachyose | | | |
| D6 α-Keto-Glutaric Acid | +++ | +++ | | D6 D-Tagatose | | | |
| D7 α-Keto-Butyric Acid | | | | D7 Turanose | | | |
| D8 α-Methyl-DGalactoside | | | | D8 Xylitol | | | |
| D9 α-D-Lactose | | | | D9 N-Acetyl-DGlucosaminitol | | | |
| D10 Lactulose | | | | D10 γ-Amino Butyric Acid | | | |

| | | | | | | | |
|---|---|---|---|---|---|---|---|
| D11 Sucrose | | | | D11 δ-Amino Valeric Acid | | | |
| D12 Uridine | +++ | | | D12 Butyric Acid | | +++ | +++ |
| E1 L-Glutamine | +++ | | | E1 Capric Acid | +++ | | |
| E2 m-Tartaric Acid | | | | E2 Caproic Acid | +++ | +++ | |
| E3 D-Glucose-1-Phosphate | +++ | | | E3 Citraconic Acid | | | |
| E4 D-Fructose-6-Phosphate | +++ | | | E4 Citramalic Acid | | | |
| E5 Tween 80 | | | | E5 D-Glucosamine | +++ | | |
| E6 α-Hydroxy Glutaric Acid-γLactone | | | | E6 2-Hydroxy Benzoic Acid | | | |
| E7 α-Hydroxy Butyric Acid | +++ | | | E7 4-Hydroxy Benzoic Acid | | | |
| E8 ß-Methyl-DGlucoside | | | | E8 ß-Hydroxy Butyric Acid | | +++ | +++ |
| E9 Adonitol | +++ | | | E9 Glycolic Acid | | | |
| E10 Maltotriose | | | | E10 α-Keto-Valeric Acid | | | |
| E11 2-Deoxy Adenosine | +++ | | | E11 Itaconic Acid | | | |
| E12 Adenosine | +++ | | | E12 5-Keto-DGluconic Acid | | | |
| F1 Glycyl-L-Aspartic Acid | +++ | | | F1 D-Lactic Acid Methyl Ester | | | |
| F2 Citric Acid | +++ | | | F2 Malonic Acid | | | |
| F3 myo-Inositol | | | | F3 Melibionic Acid | | | |
| F4 D-Threonine | | | | F4 Oxalic Acid | | | |
| F5 Fumaric Acid | +++ | +++ | +++ | F5 Oxalomalic Acid | | | |
| F6 Bromo Succinic Acid | +++ | +++ | +++ | F6 Quinic Acid | | | |
| F7 Propionic Acid | +++ | +++ | | F7 D-Ribono-1,4- Lactone | | | |
| F8 Mucic Acid | | | | F8 Sebacic Acid | | | |
| F9 Glycolic Acid | | | | F9 Sorbic Acid | | | |
| F10 Glyoxylic Acid | | | | F10 Succinamic Acid | | | |
| F11 D-Cellobiose | | | | F11 D-Tartaric Acid | | +++ | |
| F12 Inosine | +++ | | | F12 L-Tartaric Acid | | | |
| G1 Glycyl-LGlutamic Acid | +++ | | | G1 Acetamide | | | |
| G2 Tricarballylic Acid | | | | G2 L-Alaninamide | | | |
| G3 L-Serine | +++ | | | G3 N-Acetyl-LGlutamic Acid | | | |
| G4 L-Threonine | +++ | | | G4 L-Arginine | +++ | | |
| G5 L-Alanine | +++ | | | G5 Glycine | | | |
| G6 L-Alanyl-Glycine | +++ | | | G6 L-Histidine | +++ | | |
| G7 Acetoacetic Acid | +++ | | | G7 L-Homoserine | | | |
| G8 N-Acetyl-ß-DMannosamine | | | | G8 Hydroxy-LProline | +++ | | |
| G9 Mono Methyl Succinate | | | | G9 L-Isoleucine | | | |
| G10 Methyl Pyruvate | +++ | +++ | | G10 L-Leucine | +++ | | |
| G11 D-Malic Acid | +++ | | | G11 L-Lysine G | | | |
| G12 L-Malic Acid | +++ | +++ | +++ | 12 L-Methionine | +++ | | |
| H1 Glycyl-L-Proline | +++ | | | H1 L-Ornithine | | | |
| H2 p-Hydroxy Phenyl Acetic Acid | +++ | | | H2 L-Phenylalanine | +++ | | |
| H3 m-Hydroxy Phenyl Acetic Acid | +++ | | | H3 L-Pyroglutamic Acid | +++ | | |
| H4 Tyramine | +++ | | | H4 L-Valine | | | |
| H5 D-Psicose | | | | H5 D,L-Carnitine | | | |
| H6 L-Lyxose | | | | H6 Sec-Butylamine | | | |
| H7 Glucuronamide | | | | H7 D,L-Octopamine | | | |
| H8 Pyruvic Acid | +++ | +++ | | H8 Putrescine | | | |
| H9 L-Galactonic Acid-γ-Lactone | | | | H9 Dihydroxy Acetone | | | |
| H10 D-Galacturonic Acid | | | | H10 2,3-Butanediol | | | |
| H11 Phenylethylamine | | | | H11 2,3-Butanedione | | | |
| H12 2-Aminoethanol | | | | H12 3-Hydroxy-2- Butanone | | | |

### F. Denitrification phenotypes of isolated organisms

Here we present a series of experiments with each of the three isolated organisms, designed to characterize their denitrification regulatory phenotype, with emphasis on regulatory traits that could determine their capacity to function as sinks for $N_2O$ in soil. The section starts with a synopsis of the results, with references to the subsequent figures showing the results of individual experiments.

In these experiments, cells were raised under strict aerobic conditions to secure negligible amounts of denitrification reductases in the cells. They were then inoculated to 120 mL vials with He + ~1vol% $O_2$ (with or without $N_2O$) in the headspace, containing 50 mL of Sistrom's succinate medium, either with $NO_3^-$ or $NO_2^-$ (1 mM), and with a Teflon-coated magnetic bars). The vials were placed in the thermostatic water bath (20 °C) of the incubation robot, stirred continuously at high speed (700 rpm), and monitored for gas kinetics ($O_2$, NO, $N_2O$ and $N_2$) by frequent sampling of the headspace as the culture grows by aerobic respiration, depletes the oxygen and is forced to switch to denitrification. For each gas sample withdrawn, an equal volume of He is returned, and this dilution by sampling is taken into account when estimating the rates of gas production/consumption. Miniscule leakage of $N_2$ during sampling (40 -100 nmol) is also taken into account. In addition to the automatized gas sampling, small liquid volumes 20-100 µL were withdrawn manually (syringe) for determining the concentration of $NO_2^-$. The measured concentration of each gas in the headspace is used to calculate its concentration in the liquid, and the molar amount per vial (see Molstad et al 2007).

Each experiment is normally continued until metabolism comes to a halt due to depletion of all electron acceptors, i.e. that the only N-gas present is $N_2$, and that the production of $N_2$ comes to a halt (cumulative $N_2$ reach a stable plateau). NB: cumulative $N_2$ is total amount of $N_2$ produced at any time $t$ is $N_t = N_t - N_0 + SN_t - LN_t$, where $N_t$ is measured amount of $N_2$ (vial$^{-1}$) at time $t$, $N_0$ is the measured initial $N_2$ in the vial, $SN_t$ is the amount of $N_2$ removed by all samplings prior to $t$ and $LN_t$ is the amount of $N_2$ leaked into the vial prior to sampling at time $t$.

Since the initial concentration of $NO_3^-$, $NO_2^-$ and $N_2O$ in each vial is known, and that all N-gases (NO, $N_2O$ and $N_2$) are quantified, N-mass balance can be calculated throughout each experiment. For such mass balance, the sampling loss of $N_2O$ and NO is also taken into account). This is useful for two purposes: 1: to check if the initial amounts of $N_2O$+$NO_3^-$+$NO_2$-N is recovered as $N_2$-N at the end, i.e. when cumulated $N_2$ reach a plateau, 2: to estimate the concentration of $NO_3^-$ (or $NO_2^-$) throughout the incubation by mass balance calculation. For obvious reasons 2) can only be done with confidence if 100 % conversion to $N_2$ is confirmed, which was the case for all experiments (+/- 5%, ascribed to experimental error).

The convention when reporting the results is to express the amounts of each N-species as molar amounts if N per vial (2 mol N per mol $N_2O$ and $N_2$!), to make the presentations more transparent with respect to N mass balance (1 mol $NO_3^-$ is converted to 0.5 mol $N_2$, but 2 mol $N_2$-N). The concentrations in the liquid are reported conventionally however (ex: 1 nM $N_2O$ is 1 nmol $N_2O$ $L^{-1}$).

The elaborated routines for calculating rates of production/consumption of each gas has been explained in detail by Molstad et al (2007), and the excel program is freely available (Bakken 2020)

**Synopsis of the results**

***Azonexus sp*. (AN) Fig S15-17. AN** reduced $NO_3^-$ quantitatively to $N_2$, with miniscule transient accumulation of $N_2O$ (**Fig S15).** When provided with both $N_2O$ and $NO_3^-$, all electrons were directed to $N_2O$ reductase until the external $N_2O$ was depleted (**Fig S15ABD**). This was expected since the nitrate reductase in **AN** is periplasmic (Nap), and the study of other organisms with Nap has demonstrated that the electron flow to $N_2O$ reductase (Nos) outcompetes that to Nap when $N_2O$ is available in excess (Mania et al 2020). **AN** was also apparently *bet hedging*: The electron flow rate declined as the culture switched from oxic to anoxic respiration, and increased exponentially therafter (**Fig S15C**), which is the typical pattern for a denitrifying organism that performs *bet hedging*. Such organisms express one (or several) of the denitrification enzymes only in a minority of the cells, as demonstrated for *Paracoccus denitrificans* (Lycus et al 2018). The denitrification kinetics indicate that **AN** is *bet hedging* with respect to nitrate reductase (Nap) (**Fig S15, 16**), i.e. that a minority of cells express Nap, while all cells express Nos and Nir, which was corroborated by proteomic analyses which showed very high Nos/Nap protein abundance ratio after transition to anoxic respiration (**Fig S17**). As a consequence, the majority of cells can only reduce (not produce) $N_2O$ The *bet hedging* with respect to Nap and the strong competitive edge of Nos versus Nap for electrons explains the cultures capacity to keep $N_2O$ extremely low when respiring $NO_3^-$ (**Fig S15**), while producing >3 orders of magnitude more $N_2O$ when provided with $NO_2^-$ (**Fig S16**).

***Azospira* sp. (AS) Fig S18-21.** The phenotype of **AN** was similar to that of **AS**: marginal transient $N_2O$ accumulation when provided with $NO_3^-$ (**Fig S18**), preferential electron flow to Nos versus Nap (**Fig S20**), but not versus Nir (**Fig S21**), and hence higher $N_2O$ accumulation, by 2 – 3orders of magnitude, when provided with $NO_2^-$ compared to $NO_3^-$ (**Fig S19**). The electron flow rate during the transition from oxic to anoxic respiration of $NO_3^-$ showed a modest decline in response to oxygen depletion, suggesting that at least 50% of the cells expressed nitrate reductase (**Fig S18**). In contrast, the transition from oxic to anoxic respiration of $NO_2^-$ was "seamless" (i.e. no depression, **Fig S19**), suggesting that all cells expressed nitrite reductase.

***Pseudomonas sp*. (PS) Fig S22-24.** The electron flow rates in **PS** during the transition from oxic to anoxic respiration suggested *bet hedging* with respect to the expression of nitrite reductase (**Fig S22 panel B2**), but not nitrate reductase (**Fig S22 panel A2**), and the isolate demonstrated fast depletion of externally provided $N_2O$ both in the presence of $NO_3^-$ (**Fig S23**) and $NO_2^-$ (**Fig S24**). The gas kinetics indicate that $N_2O$ reductase in this organism is a very strong sink for electrons, outcompeting both nitrite and nitrate reductase. The steady state $N_2O$ concentration during anaerobic respiration was low: 50 nM whe respiring $NO_3^-$ and 200 nM when respiring $NO_2^-$ (**Fig S24**).

Based on the above phenotypes, **PS** stands out as the most robust $N_2O$ sink in a complex environment like soil, where $NO_2^-$ inevitably will be produced by other organisms, in response to oxygen depletion.

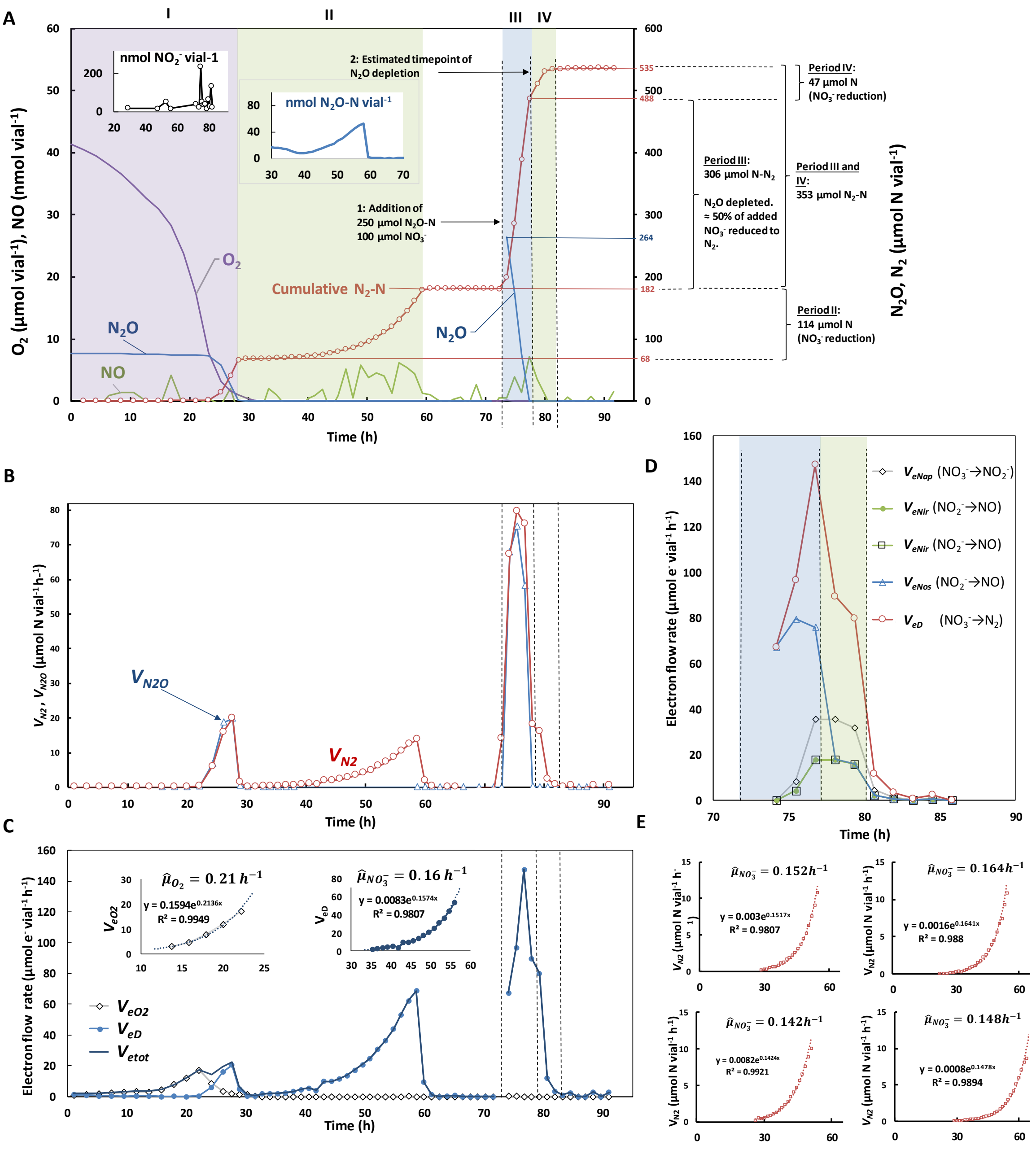


**Figure S15: Denitrification phenotype of *Azonexus sp.* (AN) when provided with $N_2O$ and $NO_3^-$.** The panels A-D show kinetics of gases and $NO_2^-$, calculated electron flow rates and estimation of growth parameters for **AN** grown in gas tight 120 mL vials, initially supplemented with 1 mL $O_2$, 1 mL $N_2O$ and 2mM $NO_3^-$ in 50 mL Sistrom's succinate medium (headspace volume = 70 mL). The vails were incubated at constant temperature and stirring (20 °C, 700 rpm), and given a dose of 250 µmol $N_2O$-N, and 100 µmol $NO_3^-$ after 72 hours. All gases are reported in molar amounts per vial.

Panels A-D show results for a single vial (2 replicate vials gave very similar results, except for a time frameshift with respect to $NO_3^-$ reduction). Panel A: measured $O_2$, NO and $N_2O$ and cumulative $N_2$ throughout the incubation (cumulative $N_2$ is the measured $N_2$ corrected for leakage and loss of $N_2$ by sampling, see materials and methods). Inserted panels show measured $NO_2^-$ (nmol vial$^{-1}$) and $N_2O$ (nmol N vial$^{-1}$). The panel highlights four periods: I, reduction of initial $O_2$ and $N_2O$; II, reduction of initial $NO_3^-$; III, reduction of the injected 250 µmol $N_2O$-N ; IV, subsequent reduction of the remaining $NO_3^-$ (100 µmol $NO_3^-$ was injected together with $N_2O$ at the beginning of period III). Panel B: $N_2$ production rate (***$V_{N2}$***) and $N_2O$-reduction rate (***$V_{N2O}$***; this is the rate at which the externally provided $N_2O$ was reduced). Panel C: Electron flow rates: ***$V_{eO2}$*** is the electron flow rate to terminal oxidases (electron acceptor = $O_2$), $\boldsymbol{V_{eD}}$ is the electron flow rate to denitrification reductases (electron acceptors= $NO_3^-$, $NO_2^-$, NO, and $N_2O$), $\boldsymbol{V_{etot}}$ = $\boldsymbol{V_{eO2}}$ + $\boldsymbol{V_{eD}}$. The inserted panels show exponential regression of $\boldsymbol{V_{eO2}}$ and $\boldsymbol{V_{eD}}$ against time, thus estimating the aerobic and anaerobic growth rates ($\hat{\mu}_{O2}$ =0.21 $h^{-1}$, $\hat{\mu}_{NO3}$ =0.16 $h^{-1}$). Panel D: Electron flow rates to individual N-reductases (and the sum of all) during the periods III and IV, illustrating the preferential electron flow to Nos ($N_2O \rightarrow N_2$).

Panel E shows the result of a separate experiment; four replicate vials supplemented with 2 mM $NO_3^-$ and 1 mL $O_2$ (no external $N_2O$ supplied), and initial $OD_{660nm}$ = 0.0064 (inoculum 1 mL, $OD_{660nm}$= 0.32). The panels show exponential regression of $N_2$ production rates for each individual vial. The kinetics of $O_2$-reduction (not shown) and $N_2$-production were used to estimate the fraction of cells expressing Nap ($F_{den}$), using the model of Hassan et al (2016). The $F_{den}$ estimates for the individual vials were 0.12 (vial 1), 0.04 (vial 2), 0.14 (vial 3), and 0.02 (vial 4). The values indicate that **AN** is *bet hedging* with respect to expression of Nap, but that the fraction of cells that express Nap ($F_{den}$) varied grossly between vials. Estimated $F_{den}$ for 9 individual vials (same type of experiment, results not shown) were done, and the $F_{den}$ estimates ranged from 0.006 to 0.24, average=0.07, stdev=0.08 (result not shown). The NO concentrations during denitrification were invariably low: 0-7 nmol $vial^{-1}$, which is equivalent to 0-5 nM NO in the liquid (1 nmol vial = 0.71 nM in the liquid at the given temperature (20 °C). Likewise, the concentration of $N_2O$ was extremely low during denitrification: in the vials with $NO_3^-$ only (panel E), the $N_2O$-level was 2-4 nmol $N_2O$ $vial^{-1}$ (=6.4-12.8 nM $N_2O$ in the liquid, 1 nmol $N_2O$-N vial = 3.27 nM $N_2O$ in the liquid) during the early phase of $NO_3^-$ reduction, increasing gradually to 5-9 nmol $N_2O$ $vial^{-1}$ during the period with exponentially increasing rates of $NO_3^-$ reduction (results not shown). Such gradual increase in $N_2O$ concentration is expected for a *bet hedging* organism which expresses $N_2O$ reductase in all cells and nitrate reductase only in a minority (see Hassan et al 2016).

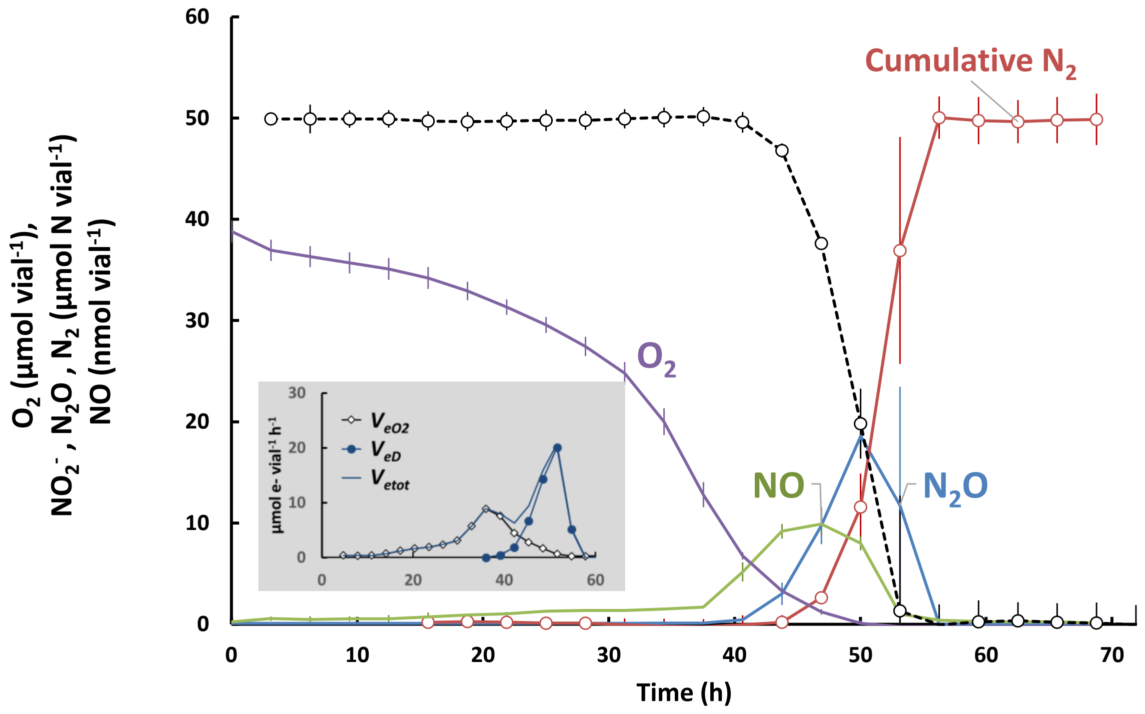


**Fig S16**: **Denitrification phenotype of *Azonexus sp*. (AN) when provided with $NO_2^-$.** The experimental conditions were as for **Fig S15**, but with 1 mM $NO_2^-$ = 50 µmol $NO_2^-$ $vial^{-1}$ (no $NO_3^-$) in the medium. The panel shows measured $O_2$, NO and $N_2O$ and cumulative $N_2$ throughout the incubation (cumulative $N_2$ is the measured $N_2$ corrected for leakage and loss of $N_2$ by sampling, see materials and methods), and $NO_2^-$ calculated by N-mass balance (initial $NO_2^-$-N minus N recovered as NO+$N_2O$ + $N_2$-N), all with standard deviation shown as vertical lines (n=2). Peak NO concentrations were ~10 nmol $vial^{-1}$ (~7 nM in the liquid), which is slightly higher than that in the $NO_3^-$ -fed cultures (**Fig S15A**), while the peak $N_2O$ (20 µmol $N_2O$-N $vial^{-1}$) is >3 orders of magnitude higher than during denitrification of $NO_3^-$ (**Fig S15A**). The inserted panel shows the electron flow rates; $\boldsymbol{V_{eO2}}$ is the electron flow rate to terminal oxidases (electron acceptor = $O_2$), $\boldsymbol{V_{eD}}$ is the electron flow rate to denitrification reductases (electron acceptors= $NO_2^-$, NO, and $N_2O$), $\boldsymbol{V_{etot}}$ = $\boldsymbol{V_{eO2}}$ + $\boldsymbol{V_{eD}}$. The relatively seamless transition to anoxic respiration (indicated by the only marginal depression in $\boldsymbol{V_{etot}}$ at oxygen depletion) suggests that the majority of cells expressed nitrite reductase.

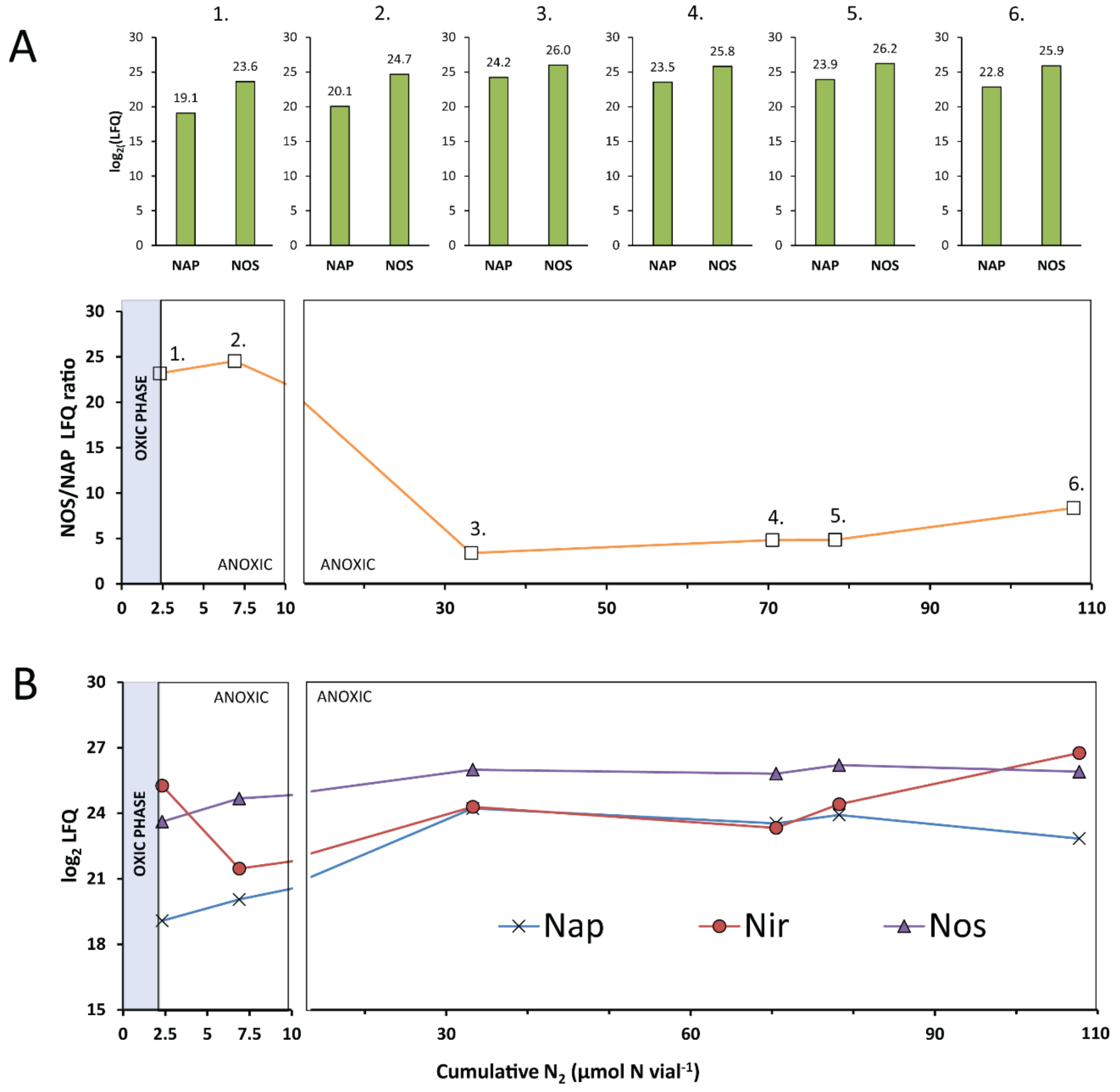


**Figure S17**: **Quantification of denitrification reductases for *Azonexus* sp. (AN) by proteomics.** Aerobically grown **AN**- cells were inoculated in replicate vials with 50 mL Sistrom succinate medium supplemented with 2 mM $NO_3^-$ and 1 mL $O_2$ (Initial $OD_{660}$ = 0.003 vial$^{-1}$), as for the experiment shown in **Figure S15.** Single vials where periodically subjected to destructive sampling and proteomic analysis throughout the incubation (six vials in total, numbered 1- 6). Vial 1 was analyzed at the oxic/anoxic transition (0.7 µM $O_2$ in the liquid). Relative LFQ values were corrected for nitrogen reductases with multiple identical subunits (NOS and NIR). Panel A: $log_2$(LFQ) values for Nap and Nos (bars) and their ratio, for each sample plotted against the vial-specific cumulative $N_2$-N (µmol vial$^{-1}$) at the time of destructive sampling. Panel B: $log_2$(LFQ) assigned Nap, Nir and Nos plotted against the cumulative $N_2$. Nor was only detected in the final sample. Each figure so split in two parts, with different scales for X-axis to improve visibility of the initial changes.

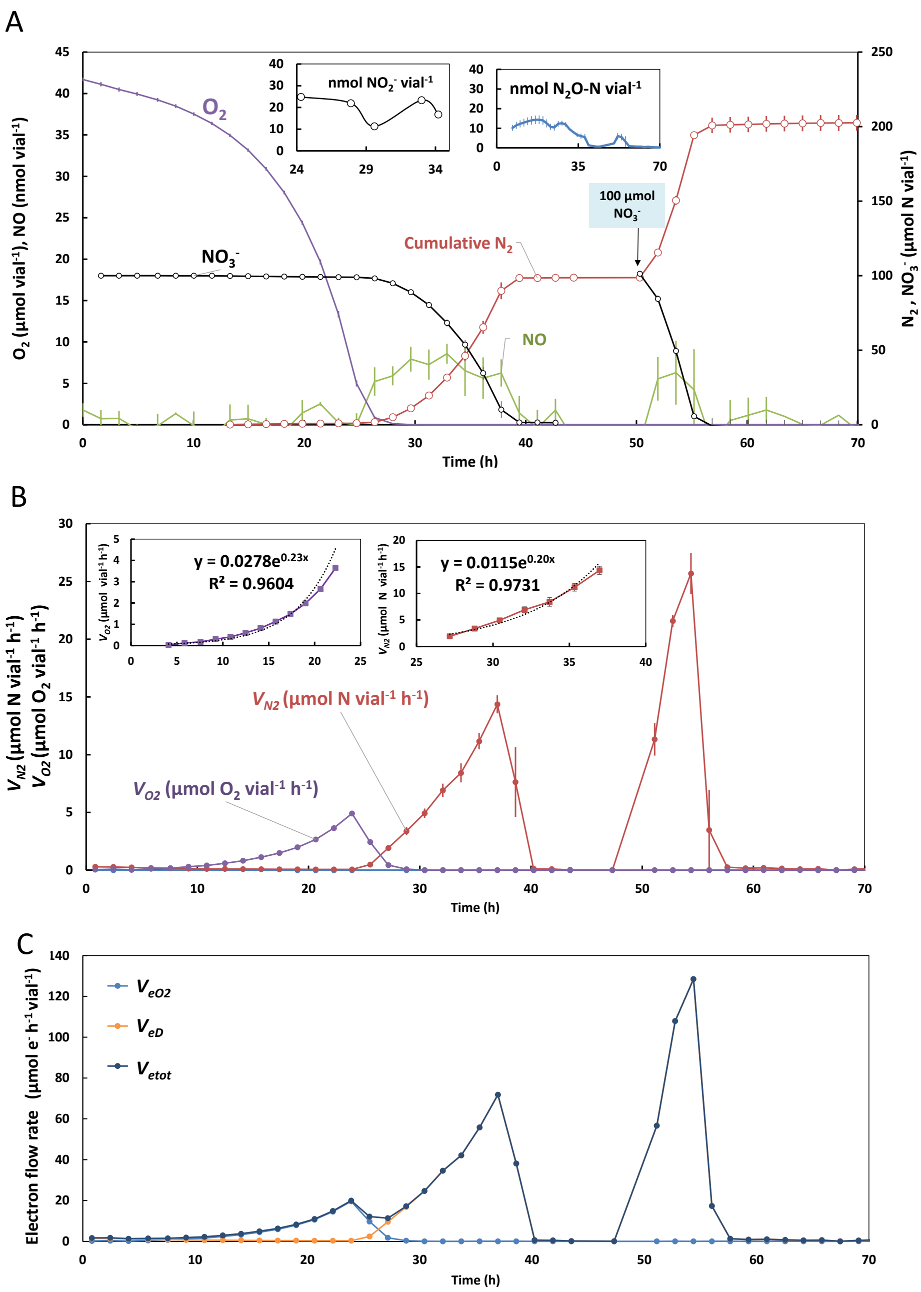


**Figure S18**: **Denitrification phenotype of *Azospira sp.* (AS), provided with $NO_3^-$**. The experimental conditions were as for **Fig S15** but without $N_2O$ in the headspace. After 50 h, 100 µmol $NO_3^-$ (to a final concentration of 2 mM) was injected. The initial inoculum had $OD_{660}$ = 0.030 (1 mL added to the 50 mL medium in the vials). Error bars indicate standard deviations (n = 3). Panel A: measured $O_2$, NO, $N_2O$, cumulative $N_2$ and $NO_3^-$ calculated by N-mass balance (initial $NO_3^-$-N minus N recovered as NO+$N_2O$+$N_2$-N). Inserted panels show measured $NO_2^-$ (nmol vial$^{-1}$) and $N_2O$ (nmol N vial$^{-1}$). All the denitrification intermediates ($NO_2^-$, NO and $N_2O$) were extremely low during denitrification. Panel B: Rates of $O_2$-consumption ($V_{O2}$) and $N_2$- production ($V_{N2}$). The inserted panels show exponential regression of $V_{O2}$ (oxic phase) and $V_{N2}$, estimating aerobic and anaerobic growth rates (0.23 and 0.20 h$^{-1}$, respectively). Panel C: Calculated electron flow rates: $V_{eO2}$ is the electron flow rate to terminal oxidases (electron acceptor = $O_2$), $V_{eD}$ is the electron flow rate to denitrification reductases (electron acceptors= $NO_3^-$, $NO_2^-$, NO, and $N_2O$), $V_{etot}$ = $V_{eO2}$ + $V_{eD}$. The dip in electron flow in response to oxygen depletion suggests some *bet hedging* with respect to expression of nitrate reductase (i.e. that ~50% of the cells express nitrate reductase). In contrast, a transition to anoxic respiration with $NO_2^-$ showed no depression in electron flow (**Fig S19, panel C**).

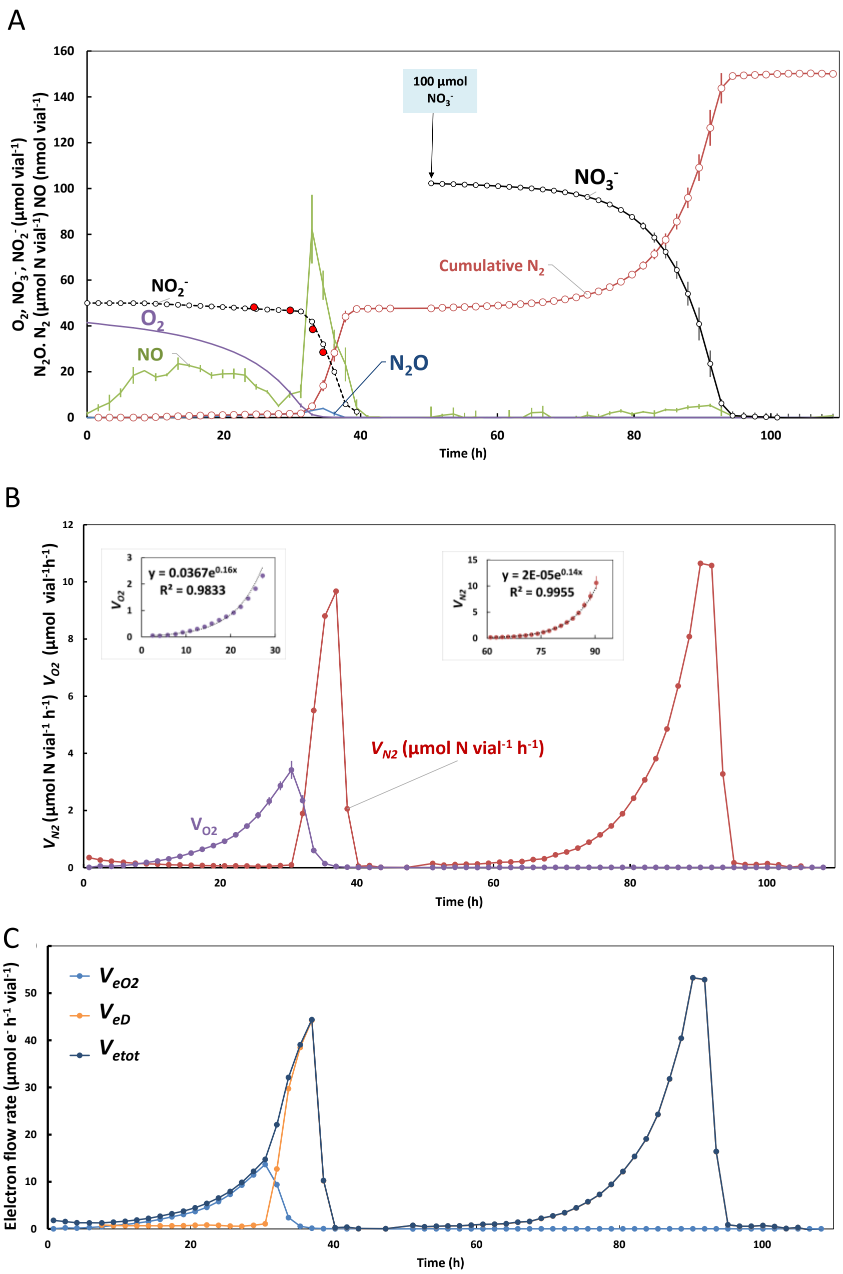


**Figure S19**: **Denitrification phenotype, *Azospira sp.* (AS), provided with $NO_2^-$.** The experimental conditions were as for **Fig S15**, but with 1 mM $NO_2^-$ (no $NO_3^-$). A dose of $NO_3^-$ (100 µmol $NO_3^-$ vial$^{-1}$) was injected after 50 hours. Error bars indicate standard deviations (n = 3). Panel A: measured $O_2$, NO, $N_2O$, cumulative $N_2$, $NO_2^-$ and $NO_3^-$. $NO_2^-$ and $NO_3^-$ were calculated by N-mass balance (initial amounts minus N recovered as NO+$N_2O$+$N_2$). Measured $NO_2^-$ is shown as red dots, which are in good agreement with the values based on mass balance. NO was present throughout the entire oxic phase (~20 nmol vial$^{-1}$ = 15 nM NO in the liquid) and peaked to 80 nmol NO vial$^{-1}$ in response to oxygen depletion. The transient $N_2O$ accumulation during $NO_2^-$ reduction reached 4 µmol $N_2O$-N vial$^{-1}$ (12 µM in the liquid), which is 3 order of magnitude higher than $N_2O$ during denitrification with $NO_3^-$ (**Fig S18**). The onset of $NO_3^-$ reduction after $NO_3^-$ injection was remarkably slow. Panel B: Rates of $O_2$-consumption (***$V_{O2}$***), $N_2$-production (***$V_{N2}$***). The inserted panels show exponential regression of ***$V_{O2}$*** (oxic phase) and ***$V_{N2}$*** (anoxic phase) estimating aerobic and anaerobic growth rates (0.16 and 0.14 h$^{-1}$, respectively). Panel C: Calculated electron flow rates: ***$V_{eO2}$*** is the electron flow rate to terminal oxidases (electron acceptor = $O_2$), ***$V_{eD}$*** is the electron flow rate to denitrification reductases (electron acceptors= $NO_2^-$, NO, and $N_2O$), ***$V_{etot}$*** = ***$V_{eO2}$*** + ***$V_{eD}$***. The seamless transition from oxygen- to nitrite-based respiration (no depression in ***$V_{etot}$***) indicate that all cells express nitrite reductase (no *bet hedging*). The response to the subsequent addition of $NO_3^-$ suggests that only a minority of the cells had expressed nitrate reductase, hence, the majority of the cells were unable to utilize nitrate for anaerobic respiration. These cells (without Nap) were likely entrapped in anoxia, without energy to synthesize Nap.

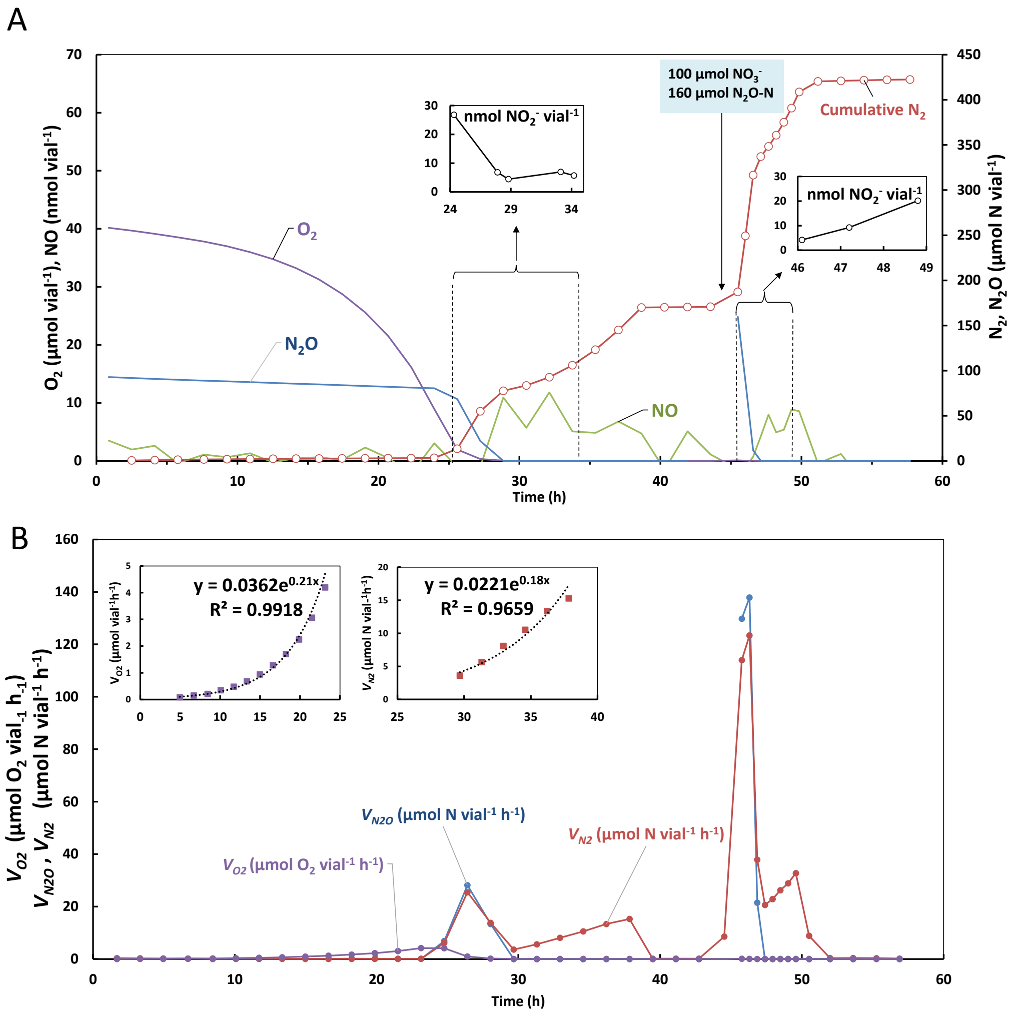


**Figure S20**: **Denitrification phenotype, *Azospira sp.* (AS), provided with $NO_3^-$ and $N_2O$**. The experimental conditions were as for **Fig S15** (2 mM $NO_3^-$ and 1 mL $N_2O$ in headspace). 0.3 mL aerobically raised pre culture ($OD_{660}$ = 0.096) was added to 50 mL medium. After 45 hours, a dose of 100 µmol $NO_3^-$ + 160 µmol $N_2O$-N (per vial) was injected. The initial inoculum had. The panels show results for a single vial. The replicate vial showed very similar gas kinetics, but with a slight time frameshift. Panel A: measured $O_2$, NO and $N_2O$ and cumulative $N_2$ production throughout the incubation. Inserted panels show measured $NO_2^-$ (nmol vial$^{-1}$). Nitrite accumulation was miniscule (inserted panels). Panel B: Rates of $O_2$-consumption ($\boldsymbol{V_{O2}}$), $N_2$-production ($\boldsymbol{V_{N2}}$) and $N_2O$-reduction ($\boldsymbol{V_{N2O}}$). NB: $\boldsymbol{V_{N2O}}$ is the rate of consumption of externally provided $N_2O$ (positive for $N_2O$ consumption). The rates of $N_2$ production ($\boldsymbol{V_{N2}}$) equaled the rates of $N_2O$-consumption ($\boldsymbol{V_{N2O}}$) during depletion of exogenous $N_2O$. This shows that during the transition from oxic to anoxic conditions, $N_2O$ was converted stoichiometrically to $N_2$, while nitrate was only reduced after depletion of $N_2O$. The inserted panels show exponential regression of $\boldsymbol{V_{O2}}$ (oxic phase) and $\boldsymbol{V_{N2}}$ (anoxic phase) estimating aerobic and anaerobic growth rates (0.21 and 0.20 h$^{-1}$, respectively).

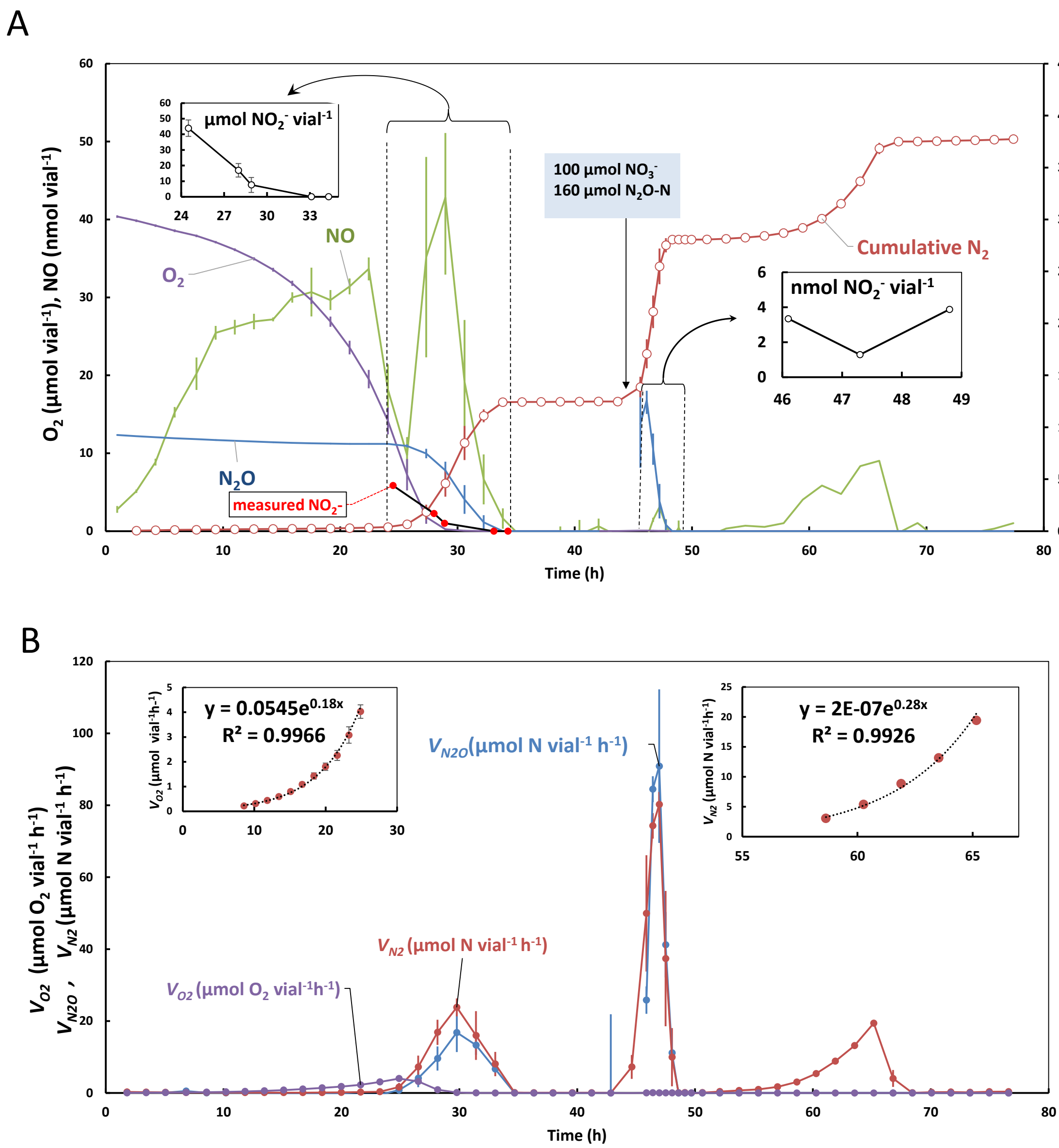


**Figure S21**: **Denitrification phenotype, *Azospira sp*. (AS) provided with $NO_2^-$ and $N_2O$**. Experimental condition as for **Fig S15**, but with 1 mL $N_2O$ and 1 mM $NO_2^-$. After 45 h, a dose of 100 µmol $NO_3^-$ + 160 µmol $N_2O$-N (per vial) was injected. The panels show the average of two replicate vials. Panel A: measured $O_2$, NO, $N_2O$, $N_2$ (cumulative) and $NO_2^-$. Measured $NO_2^-$ during the depletion of externally supplied $N_2O$ is shown in the main panel, while the miniscule $NO_2^-$ measured after injection of $NO_3^-$ is shown in the inserted panel. During the transition from oxic to anoxic conditions $NO_2^-$ - and $N_2O$- was reduced concomitantly. Following addition of 100 µmol $NO_3^-$ and 160 µmol $N_2O$-N *Azospira* sp. AS quickly reduced exogenous supplied $N_2O$, but the immediate $NO_3^-$ reduction rates was miniscule and gradually increasing. Panel B: rates of $O_2$- and $N_2O$-reduction, and $N_2$ production. During the depletion of exogenous $N_2O$ with $NO_2^-$ present (time span 22-30h), $N_2$ production rates clearly exceeded the rates of $N_2$ production, reflecting concomitant reduction of $NO_2^-$ and the exogenous $N_2O$. During the depletion of exogenous $N_2O$ in the presence of $NO_3^-$ (time span 42-47 h), the rate of $N_2$ production did not exceed the rate of $N_2O$ reduction, hence no $NO_3$-reduction took place. Inserted panels: exponential regression (against time) of the rates of $O_2$-reduction (oxic phase) and $N_2$-production.

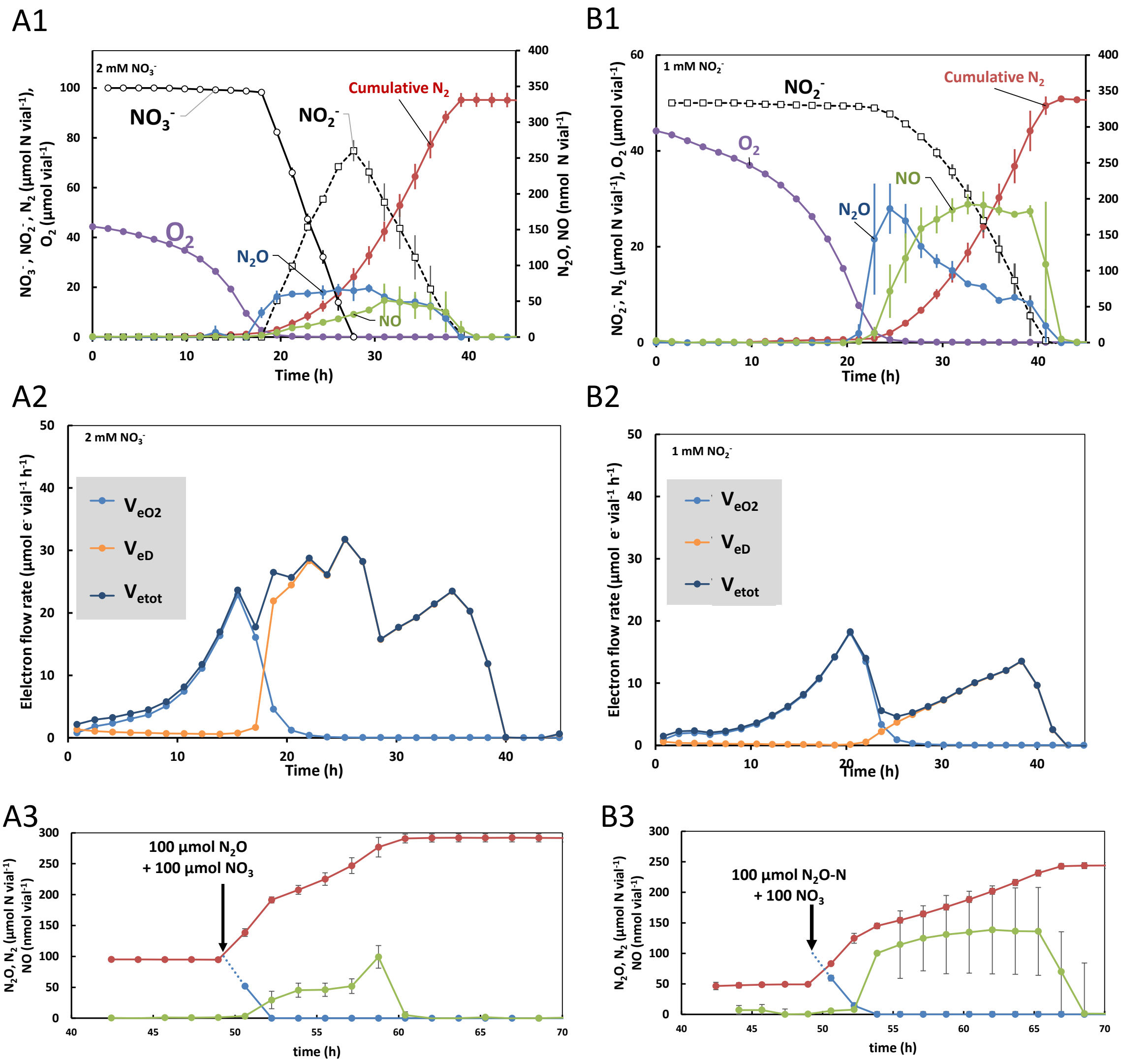


**Figure S22**: **Denitrification phenotype of *Pseudomonas sp*. (PS), provided with $NO_3^-$ or $NO_2^-$.** The experimental conditions were as for **Fig S15**; vials, with 50 mL medium were supplemented with 1 mL $O_2$ and 2mM $NO_3^-$ (=100 µmol $NO_3^-$ vial$^{-1}$ (panels A1&2, n = 3 replicate vials) or 1 mM $NO_2^-$ (Panel B1&2, n = 2 replicate vials). The vials were inoculated with 1 mL of a culture with $OD_{660nm}$ = 0.06. Panel A1: measured gases and nitrogen $NO_2^-$ in vials with 2 mM $NO_3^-$. The panel also shows $NO_3^-$ calculated by N mass balance (initial amount of $NO_3^-$- N minus N recovered as ($NO_2^-$+NO+$N_2O$+$N_2$)-N). The figure shows transient nitrite accumulation to 75 µmol vial$^{-1}$, while the NO and $N_2O$ remained very low (50 nmol NO vial$^{-1}$ ~35 nM in the liquid, 70 nmol $N_2O$-N vial$^{-1}$ ~ 230 nM $N_2O$ in the liquid). Panel A2 shows the calculated electron flow rates to $O_2$ (***$V_{eO2}$***) and to denitrification reductases (***$V_{eD}$***), and the total electron flow rate (***$V_{etot}$=$V_{eO2}$+$V_{eD}$***). The seamless transition from aerobic respiration to respiration by $NO_3^-$ -reduction (marginal reduction of **$V_{etot}$**) suggests that all cells express nitrate reductase. The reduction of **$V_{etot}$** in response to $NO_3^-$ depletion suggest that only a fraction of the cells express nitrite reductase. Panel A3: the vials (Panel A1&2) were given a dose of 100 µmol $NO_3^-$ and 100 µmol $N_2O$ after 49 hours. The kinetics reveal a strong preference for $N_2O$ over $NO_3^-$.

Panel B1: Measured gases in vials with 1 mM $NO_2^-$. The panel also shows $NO_2^-$ calculated by N mass balance (initial $NO_2^-$ minus N recovered as (NO+$N_2O$+$N_2$)-N minus that. The figure shows that transient accumulation of intermediates reached 200 nmol NO vial (~140 nM in the liquid) and 190 nmol $N_2O$-N vial$^{-1}$ (~620 nM in the liquid). Panel B2: calculated electron flow rates to $O_2$ (***$V_{eO2}$***), to denitrification reductases (***$V_{eD}$***), and the total electron flow (***$V_{etot}$***). The dip in the electron flow after the transition from aerobic to anaerobic respiration suggests that only a fraction of cells express nitrite reductase. Panel B3: the vials (B1&2) were given a dose of $NO_3$ and $N_2O$ after 49 hours.

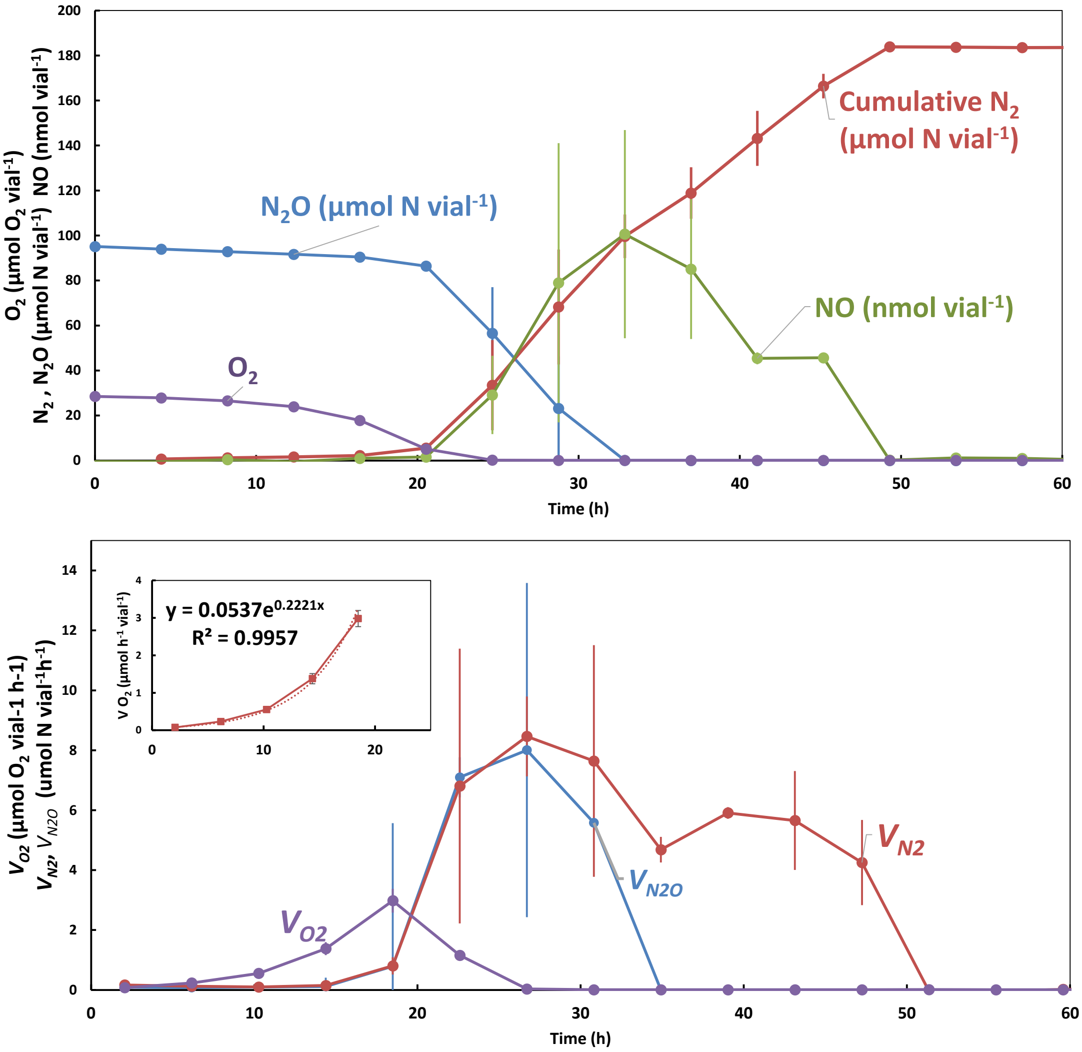


**Figure S23: Denitrification phenotype of *Pseudomonas sp*. (PS), provided with $NO_3^-$ and $N_2O$**. Experimental conditions as in **Figure 15** ($N_2O$ in headspace, 2 mM $NO_3^-$ in the medium). Panel A shows measured $O_2$, NO, $N_2O$, $N_2$. Panel B shows rates of $O_2$-consumption (***$V_{O2}$***), $N_2O$ consumption (***$V_{N2O}$***) and $N_2$-production (***$V_{N2}$***). Error bars: standard deviation, n = 2. The inserted panel shows estimated aerobic growth rate (exponential regression of $O_2$ reduction rate against time). The electron flow rates to the individual steps could not be calculated in this experiment because $NO_2^-$ was not measured.

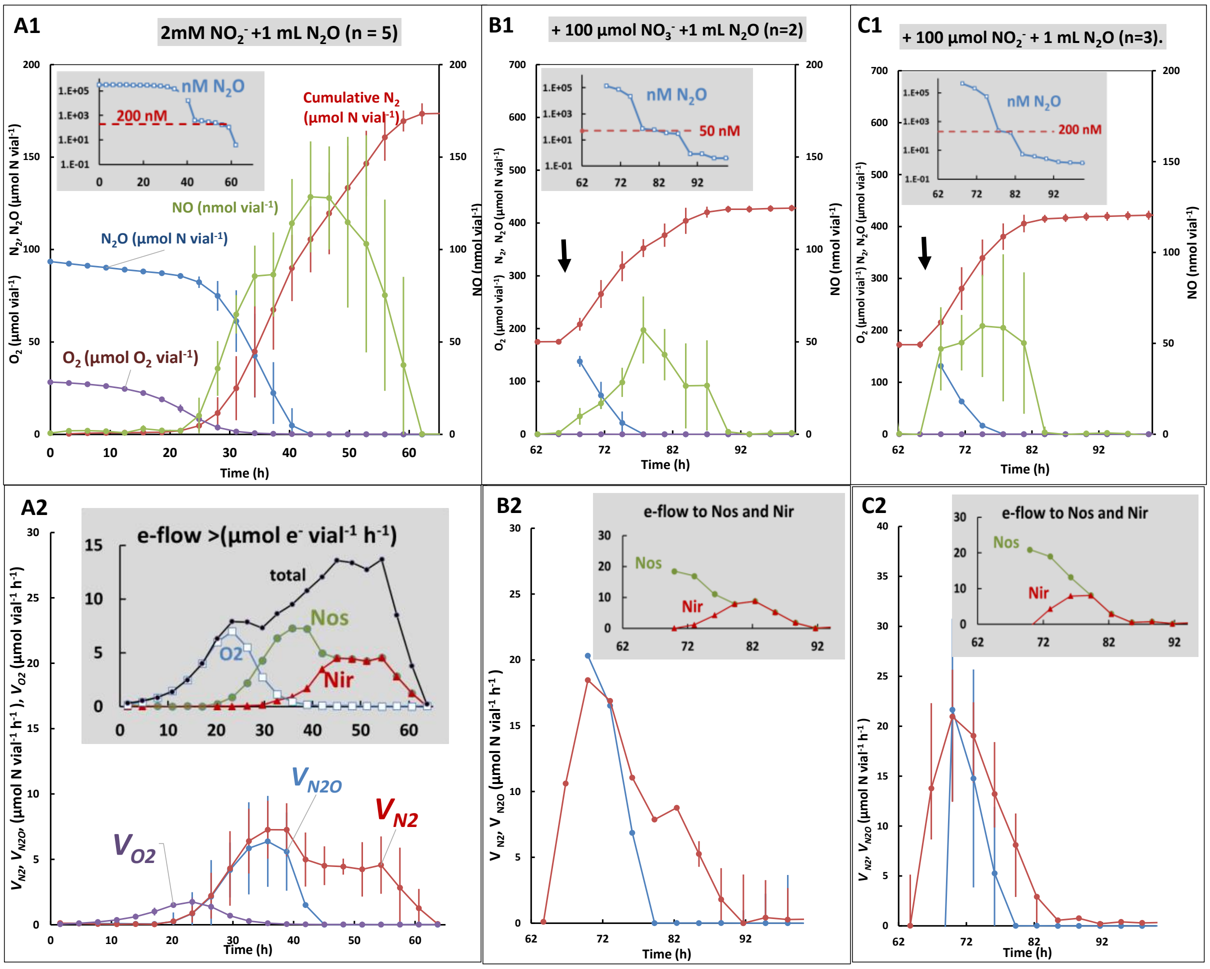


**Figure S24**: ***Pseudomonas sp*. PS, elucidating the preference for $N_2O$ versus $NO_2^-$ and $NO_3^-$**. To assess the preferential reduction of $N_2O$ versus $NO_2^-$ and $NO_3^-$, we set up an experiment with 5 vials with 2mM $NO_2^-$ and 1 mL $N_2O$ (as **Fig S15**) and monitored the gas kinetics. After depletion of all electron acceptors ($N_2O$-N + $NO_2$-N recovered as $N_2$, Panel A1 and A2), the experiment was continued by injecting more electron acceptors: two of the vials received 100 µmol $NO_3^-$ (2 mM $NO_3^-$) + 1 mL $N_2O$ (Panels B1 and B2), while three vials received 100 µmol $NO_2^-$ + 1 mL $N_2O$ (Panel C1 and C2). The time of injections are indicated by black arrows (panel B1&C1). Top panels (A1-C1) show measured gases (molar amounts per vial), with inserted panels showing the $N_2O$ concentration in the liquid (nM, log scale), illustrating steady state $N_2O$ concentrations during respiration based on nitrogen oxyanion-reduction alone, i.e. after depletion of the externally provided $N_2O$ (these steady state concentrations are indicated by dashed red lines). These steady state concentrations were ~200 nM when respiring $NO_2^-$ (panel A1 and C1), and ~50 nM when respiring $NO_3^-$ (panel B1), and again 200 nM when respiring $NO_2^-$. The lower panels (A2-C2) show calculated rates of $O_2$-consumption ($V_{O2}$), $N_2O$-depletion ($V_{N2O}$ = the rate at which exogenous $N_2O$ was depleted), and $N_2$-production ($V_{N2}$). Calculated electron flow rates are shown in the inserted panels. The insert in panel A2 shows the electron flow to terminal oxidases (marked $O_2$), and to the denitrification reductases Nir and Nos and the total (the electron flow to Nor is practically identical with that to Nir since only nanomolar amounts of NO accumulated). The inserts in panel B2 and C2 show electron flow to Nos and Nir only (no oxygen was present in these vials). The electron flow to Nar could not be estimated for (Panel B2) because $NO_2^-$ was not measured.

The initial incubation (Panels A1&A2) demonstrates a strong preference for external $N_2O$ versus $NO_2^-$, although the electron flow to Nir increased gradually as the concentration of exogenous $N_2O$ declined. In response to a second dose of $N_2O$ + $NO_2$ (panel C1 and C2), we see the same preference for $N_2O$ versus $NO_2^-$.

### G. Aerobic growth in sterilized digestate, and the effect of the enriched digestate on $N_2O$ emissions

The cultures where grown in pre-aerated digestate and supplied with $O_2$ as the terminal electron acceptor prior to inoculation in soil (Soil incubations shown in Figure 5 (main paper) and Figure S27 and S28). The pre-aeration, done before inoculation of isolated cultures by blowing air through the sterile digestate suspension for 72 hours, was necessary to secure near-complete abiotic oxidation of the $Fe^{2+}$ in the digestate before inoculation of the cultures ($FeCl_3$ is used at a precipitation chemical at the WWTP, se materials and methods). $Fe^{2+}$ would otherwise obscure the measurements of $O_2$ consumption, and possibly inhibit aerobic respiration due to formation of reactive oxygen species (Winterbourn 1995).

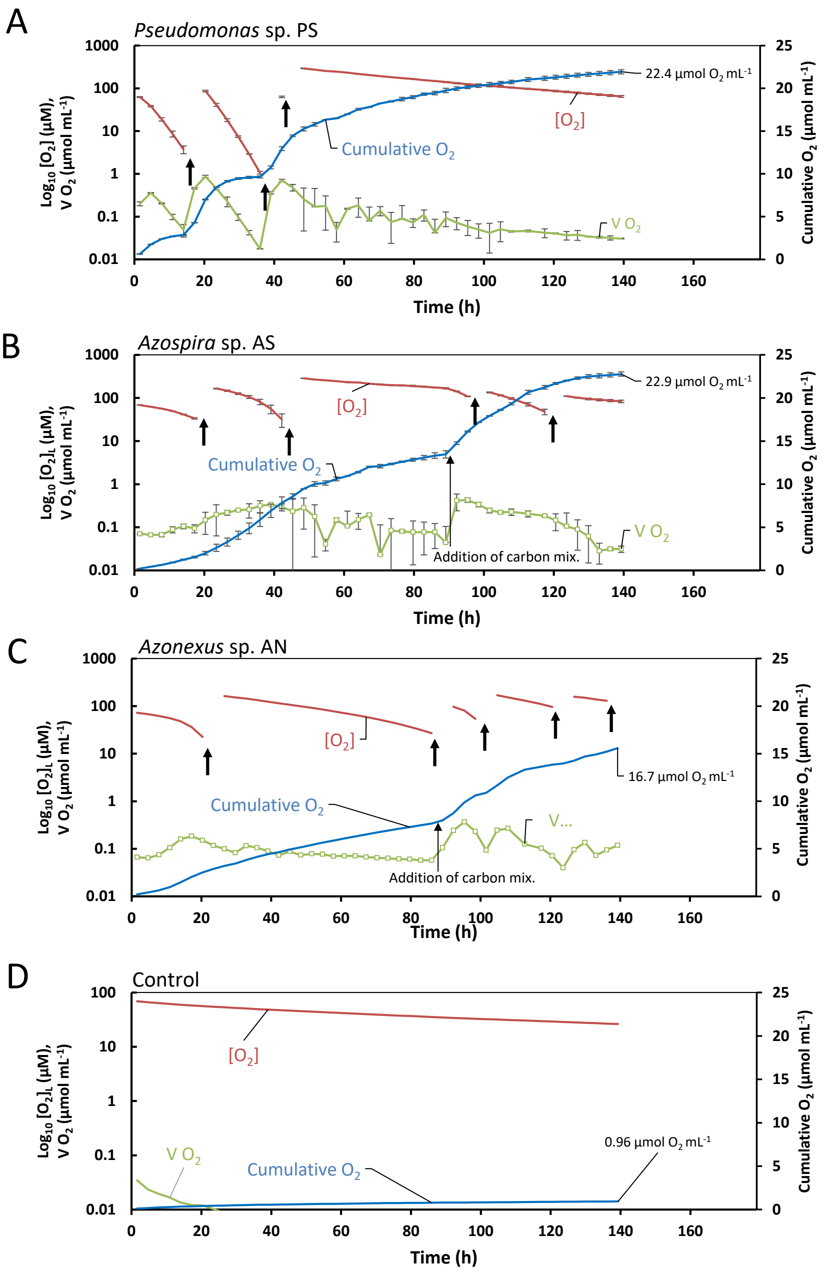


**Figure S25**: **Aerobic growth in autoclaved digestate.** 1 mL cultures of PS, AS and AN, grown oxicly (air) in stirred (700 rpm) Sistrom medium at 20 °C, were inoculated at 20 °C in closed 120 mL stirred vials (600 rpm) containing

50 mL autoclaved, pre-aerated and pH-adjusted (pH=7.5) digestate and monitored for gas concentrations of $O_2$ in the headspace. Inoculum $OD_{660nm}$ in digestate was 0.26, 0.12 and 0.30 for cultures PS, AS and AN, respectively. The vials were helium flushed and 5 mL $O_2$ was added to the headspace before inoculation of the sioaltes. Panels A to D shows calculated liquid concentration of $O_2$ (µM) and rate of oxygen consumption on the primary y-axis ($log_{10}$ scaled), and cumulative $O_2$ consumed (µmol $mL^{-1}$) throughout the incubation for the cultures PS (two replicates), AS (two replicates), AN (single vial) and a control vial containing digestate only (one vial), respectively. Error bars = standard deviation. Fat arrows represent replenishing of oxygen using a syringe piercing the rubber septum of the vials. Liquid concentration of oxygen was not calculable for the timepoint following $O_2$ addition and is therefore removed. Culture PS (Panel A) consumed significantly more $O_2$ in digestate compared to cultures AS (Panel B) and AN (Panel C). We therefore added a 1 mL of a carbon mix to vials of AN and PS (point of addition is indicated in panel B and C). The carbon mix contained 0.5 mM glutamate, 0.5 acetate, 0.5 mM puryvate and 0.5 mM ethanol dissolved in sterile water and pH adjusted to 7. The cumulative end point $O_2$ consumption per mL of digestate suspension is shown in Panel A – D. Assuming growth yields on oxygen to 1.5E14 cells $mol^{-1}$ $O_2$, as determined for *Paracoccus denitrificans* (Bergaust et al 2010, 2012), and correcting for abiotic oxygen consumption in the control (Panel D) a cell density of 3.2E09, 3.3E09 and 2.4E09 $mL^{-1}$ for *Pseudomonas sp*. PS, *Azospira sp*. AS and *Azonexus sp*. AN, respectively, was estimated.

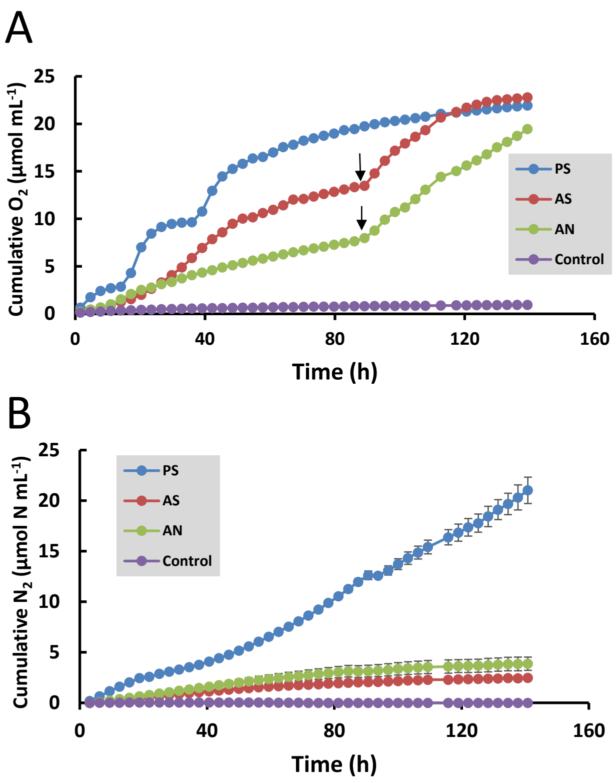


**Figure S26. Aerobic and anaerobic growth of isolatess in autoclaved digestates.** Panel A shows a compilation of the cumulated $O_2$ consumption during aerobic incubations, as shown in more detail in Fig S25. The timepoint of adding a carbon substrate cocktail to AS and AN (see Fig. S25) is indicated by arrows. (standard deviation is not included). Panel B shows the cumulated $N_2$ production in identical corresponding vials, but with a He + $N_2O$ atmosphere. No carbon substrates were added to these vials. Error bars = standard deviation (n = 3).

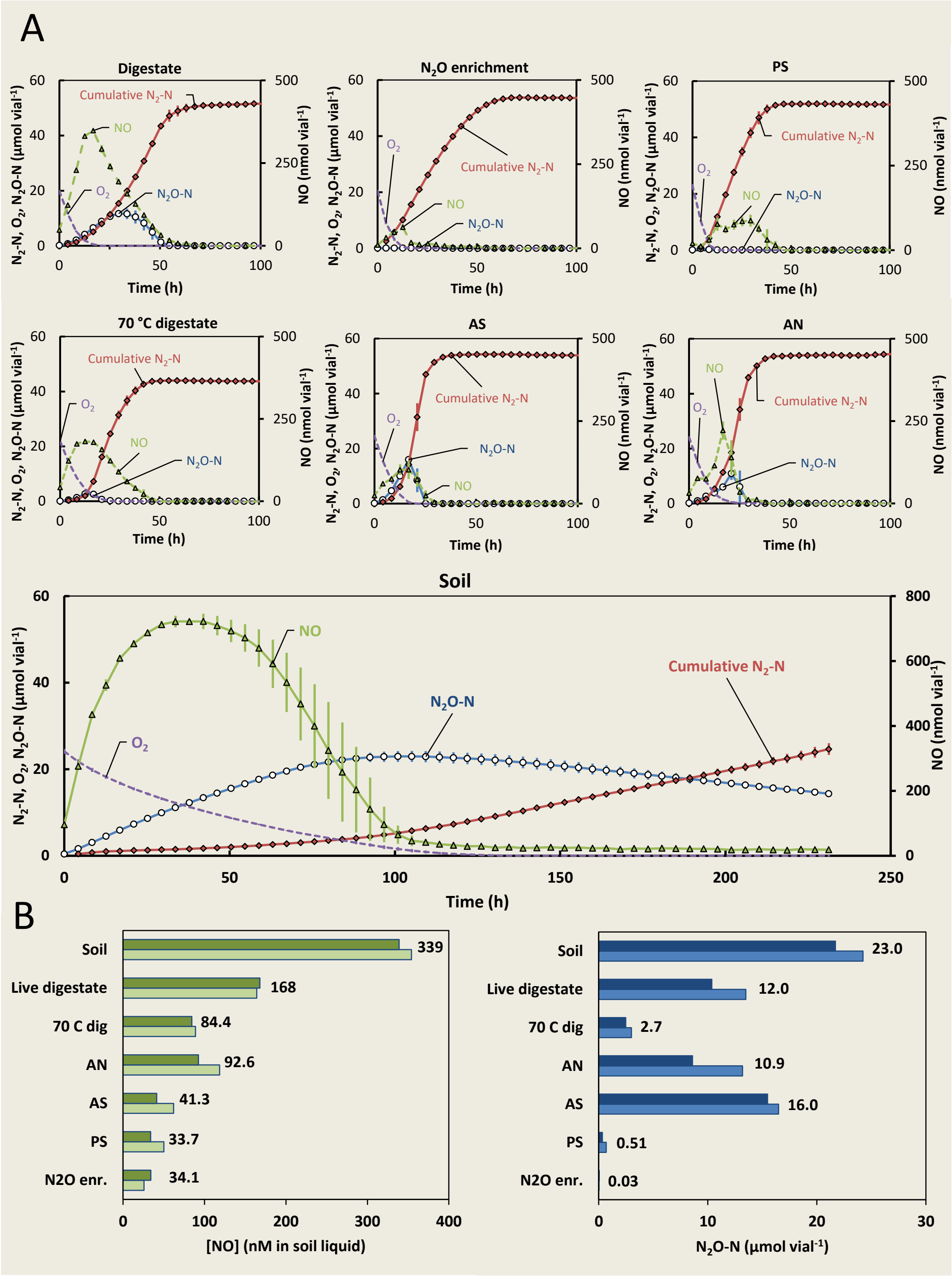


**Figure S27: Incubation of digestates treated in various ways with soil with pH=5.5.** Panel A: kinetics of $O_2$, NO, $N_2O$ and $N_2$ throughout the incubation of soils amended with the various materials (one panel for each amendment) given as molar amounts per vial. The panels show average values (n=2). The initial oxygen (~20 µmol vial$^{-1}$ corresponding to ~0.5 vol% in the headspace) was depleted within the first 20 hours for soils amended with digestates, while soil alone (lower panel) took around 100 hours to deplete $O_2$. This is due to the boost in respiration that occurs when the carbon-rich digestate is added to the soil. The amounts of $O_2$, NO and $N_2O$ are as measured, while "Cumulative $N_2$" denotes the measured $N_2$ that is corrected for leakage and losses by sampling (see Molstad et al 2007). The $N_2$ and $N_2O$ kinetics were used to calculate the $N_2O$ index ($\boldsymbol{I_{N2O}}$), which is the area under the $N_2O$-curve divided by the sum of the areas under the $N_2O$ and $N_2$-curves for a specific time span. $\boldsymbol{I_{N2O}}$ values are shown in **Fig 5** (main paper) and provide a proxy for the propensity of the system to emit $N_2O$. Panel B: peak (maximum) amounts of NO and $N_2O$ (results for single vials, 2 vials per condition; average value indicated). NO is shown as nM in the liquid phase (equilibrium concentrations with measured NO in headspace), while $N_2O$ is shown as µmol $N_2O$-N vial$^{-1}$.

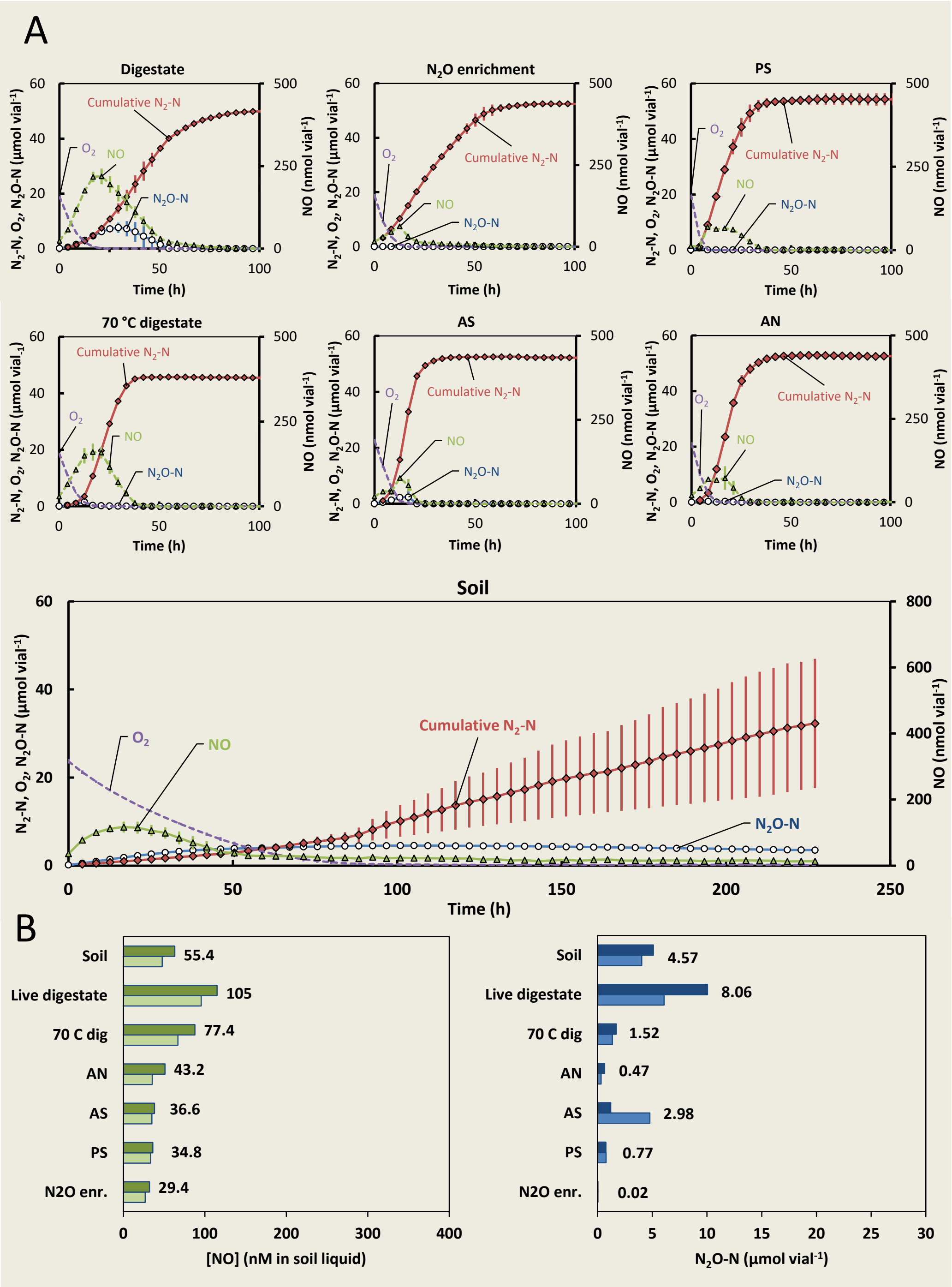


**Figure S28: Incubation of isolates and $N_2O$ enriched digestate in soil with pH=6.6.** Panel A: kinetics of $O_2$, NO, $N_2O$ and $N_2$ throughout the incubation of soils amended with the various materials (one panel for each amendment). Average values shown (n=2). The initial oxygen (~20 µmol vial$^{-1}$, ~0.5 vol% in the headspace) was depleted within the first 20 hours for soils amended with digestates, while soil alone (lower panel) took around 100 hours to deplete $O_2$. The amounts of $O_2$, NO and $N_2O$ are as measured, while "Cumulative $N_2$" denotes the measured $N_2$ that is corrected for leakage and losses by sampling (see Molstad et al 2007). The $N_2$ and $N_2O$ kinetics were used to calculate the $N_2O$ index ($I_{N2O}$), which is the area under the $N_2O$- curve divided by the area under the $N_2O+N_2$ -curve for a specific time span. $I_{N2O}$ values are shown in **Fig 5** (main paper) and is a proxy for the propensity of denitrification to emit $N_2O$. Panel B: peak (maximum) amounts of NO and $N_2O$ (results for single vials). NO is shown as nM in the liquid phase (equilibrium concentrations with measured NO in headspace), while $N_2O$ is shown as µmol $N_2O$- N vial$^{-1}$.

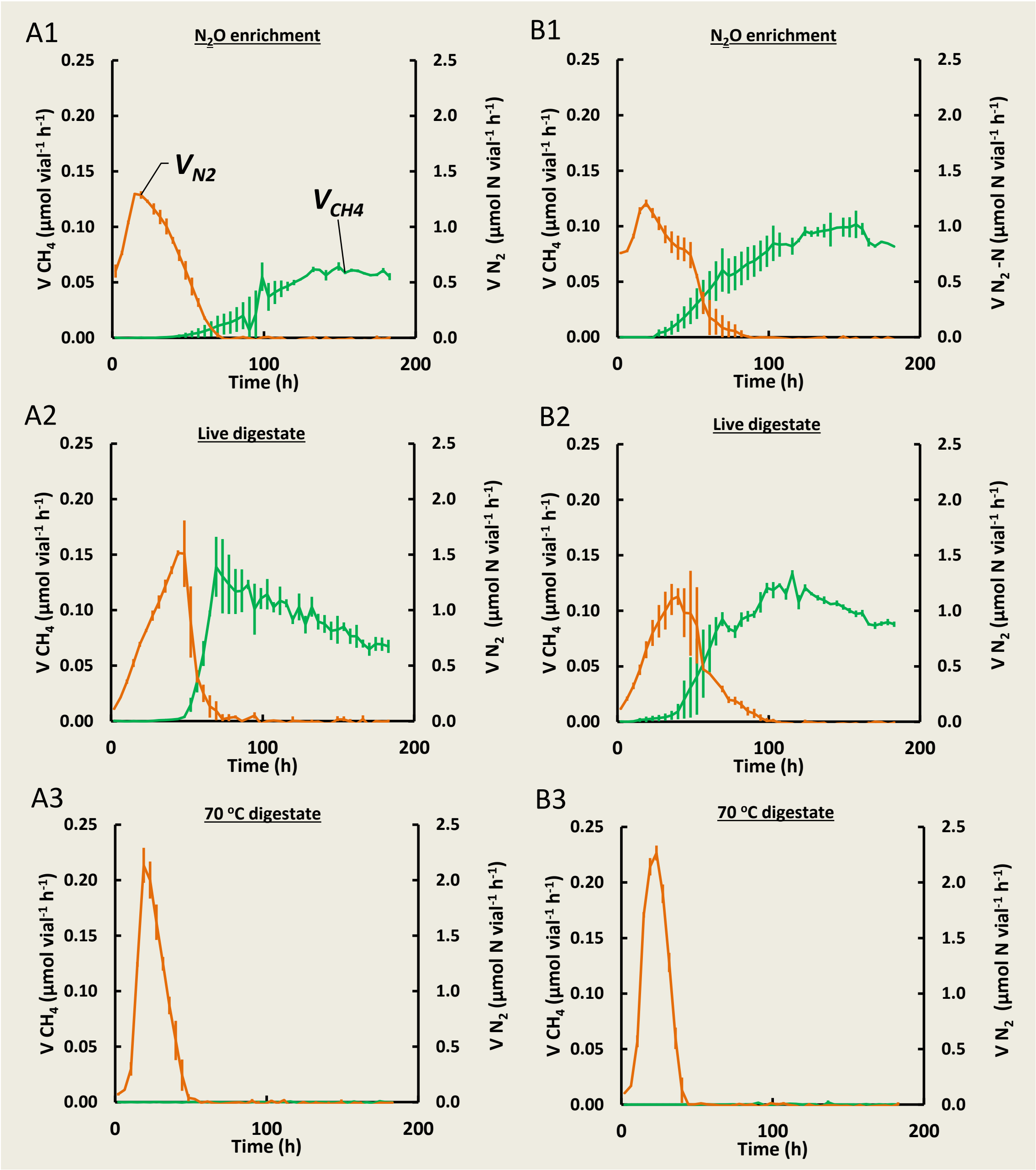


**Figure S29: Potential methanogenesis in soil amended with digestates.** The panels show the rate of $N_2$-production (***$V_{N2}$***) and rate of methane production (**$V_{CH4}$**) during incubations of soils amended with digestates (see **Figures S27 & S28**). Panels A1-A3 show results for soil with pH 5.5, and panels B1-B3 show results for soil with pH 6.6. The three panels for each soil show the results for 1) soils amended with digestate enriched with $N_2O$-reducing bacteria "$N_2O$-enrichment" (Fig 2B, main paper) 2) soils amended with "live digestate" (i.e. digestate taken directly from the anaerobic digester and 3) soils amended with digestate that had been heated to 70 °C for 2 hours. The results demonstrate that the methanogenic consortium of the digestate is alive and active in soil, but evidently suppressed by denitrification: once the nitrogen containing electron acceptors are depleted ($V_{N2}$ approaches zero), methanogenesis resumed. This did not happen in soils amended with the digestate that had been heated to 70 °C and thus likely did no longer contain live methanogenic bacteria.

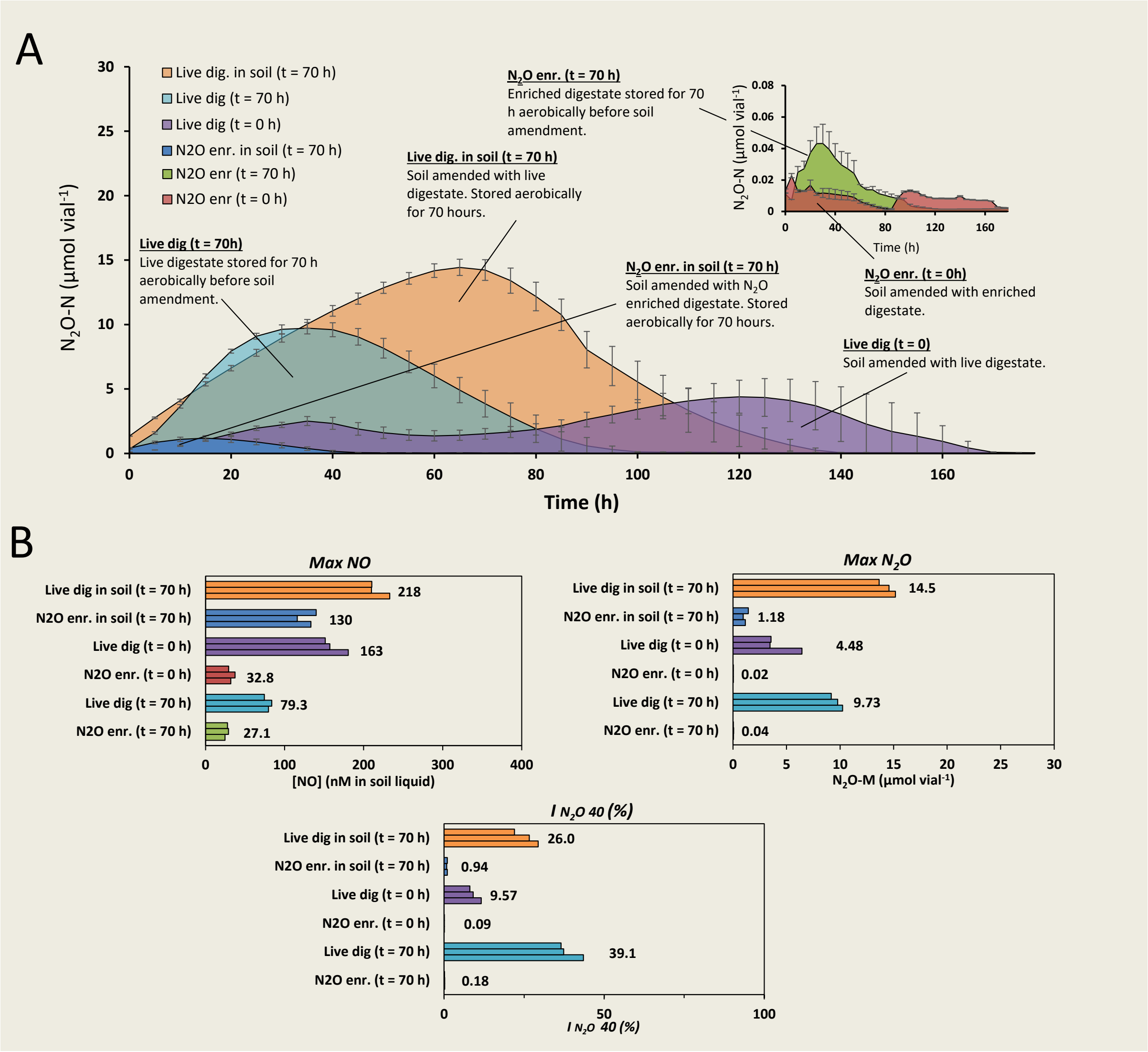


**Figure S30: Storage experiment.** To test the short term survival of the $N_2O$-scavenging capacity of $N_2O$ reducers, we added 3 mL freshly sampled live digestate or 3 mL $N_2O$ enriched digestate from the enrichment shown in Figure 2B in main paper to 10 g soil (pH 6.6) in 6 vials of which 3 were incubated anaerobically immediately for measurement of gas kinetics ("Live dig t = 0h" and "N2O enr. t = 0h"), and 3 were stored aerobically (open vials) for 70 hours in soil ("Live dig in soil t = 70h" and "N2O enr. in soil t = 70h") before being incubated anaerobically in the automated incubation and gas analysis system. In parallel, we also stored 50 mL of the freshly sampled digestate and 50 mL enrichment culture aerobically in open vials for 70 hours from which 3 mL digestate was then applied to 10 g soil (pH 6.6) ("N2O enr. t = 70 h" and "Live dig t = 70h"). Aerobic storage of digestates and digestate amended soil was conducted at 20 °C. Monitoring of gas kinetics was conducted at 20 °C. All treatments were supplemented with 50 µmol $NO_3^-$ (50 µL 1 M $KNO_3$) just before gas analysis in.

Panel A shows the $N_2O$ kinetics for all six soil treatments; the inserted panel shows the results for the two treatments with so low $N_2O$ levels that they would be invisible in the main panel (n=3 for all treatments, standard deviations are shown as vertical bars). Panel B shows maximum NO, $N_2O$ and the $N_2O$ index for 40 % recovery of $NO_3^-$ as N-gases ($I_{N2O40\%}$). NO is shown as nM in the liquid phase (equilibrium concentrations with measured NO in headspace), while $N_2O$ is shown as µmol $N_2O$- N vial$^{-1}$. The results show that within a time frame of 70 h under aerobic conditions, be it as intact enrichment or after amendment to soil, the capacity to reduce the $N_2O/N_2$ product ratio of denitrification in soil is sustained.
